# Simultaneous optimization of donor and acceptor substrate specificity for transketolase by a small but smart library

**DOI:** 10.1101/719906

**Authors:** Haoran Yu, Roberto Icken Hernández López, David Steadman, Daniel Méndez-Sánchez, Sally Higson, Armando Cázares-Körner, John M. Ward, Helen C. Hailes, Paul A. Dalby

## Abstract

A narrow substrate range is a major limitation in exploiting enzymes more widely as catalysts in synthetic organic chemistry. For enzymes using two substrates, the simultaneous optimization of both substrate specificities, is also required for the rapid expansion of accepted substrates. Transketolase catalyses the reversible transfer of a C_2_-ketol unit from a donor substrate to an aldehyde acceptor and suffers the limitation of narrow substrate scope for widely industrial applications. Herein, transketolase from *E. coli* was engineered to simultaneously accept both pyruvate as a novel donor substrate, and unnatural acceptor aldehydes, including propanal, pentanal, hexanal and 3-formylbenzoic acid. Twenty single-mutant variants were firstly designed and characterized experimentally. Beneficial mutations were then recombined to construct a small but smart library. Screening of this library identified the best variant with a 9.2-fold improvement in the yield towards pyruvate and propionaldehyde, relative to WT. Pentanal and hexanal were used as acceptors to determine stereoselectivities of the reactions, which were found to be higher than 98% *ee* for the *S* configuration. Three variants were identified to be active for the reaction between pyruvate and 3-formylbenzoic acid. The best variant was able to convert 47% of substrate into product within 24 h, whereas no conversion was observed for WT. Docking experiments suggested a cooperation between the mutations responsible for donor and acceptor acceptances, that would promote the activity towards both the acceptor and donor. The variants obtained have the potential to be used for developing catalytic pathways to a diverse range of high-value products.

## Introduction

The limited degree of substrate acceptance has been a major limitation in exploiting enzymes as catalysts in synthetic organic chemistry [1]. The fourth wave of biocatalysis is approaching [2] and the ability to discover and engineer new enzymes with expanded substrate scope is now proceeding faster than ever before [3]. For enzymes having two substrates, the simultaneous optimization of both substrate specificities is desired for rapidly expanding the scope of substrate acceptance, and range of products formed. Few studies have reported the engineering of both substrate specificities.

Transketolase (TK) catalyses the reversible transfer of a C_2_-ketol unit from D-xylulose-5-phosphate to either D-ribose-5-phosphate or D-erythrose-4-phosphate, linking glycolysis to the pentose phosphate pathway in all living cells [4, 5]. The stereospecifically controlled carbon-carbon bond forming ability of TK makes it very promising as a biocatalyst in industry, for the synthesis of complex carbohydrates and other high-value compounds [6–8]. However, to achieve industrial viability in large-scale processes, a robust transketolase toolbox is desired, in which the variants should be able to accept a diverse range of substrates including both acceptors and donors.

Wild-type TK accepts a narrow range of hydroxyaldehydes as the aldol-acceptor substrate, and with strict (*2R*)-specificity [9–11]. TK from *E. coli* has been engineered successfully for improved and reversed enantioselectivity [12], as well as an expanded aldol-acceptor substrate range including polar aliphatic [13], non-polar aliphatic [14], heteroaromatic [15, 16] and aromatic aldehyde derivatives [17, 18]. Related variants were also introduced by others, into a thermostable TK from *Geobacillus stearothermophilus* (TK_gst_) [19–22].

Wild-type TKs have very limited tolerance for different donor substrates, which require both an oxo group adjacent to the C–C bond undergoing cleavage, and a hydroxyl group at C-1. They also show strong preference towards the trans-configuration of hydroxyls at the asymmetric C-3 and C-4 positions [11]. In addition to its natural phosphorylated donors, TK can also accept hydroxypyruvate (HPA). The use of β-hydroxypyruvate as the ketol donor renders the donor half-reaction irreversible, thus increasing the atom efficiency of the reaction favourably for industrial syntheses. *E. coli* TK accepts HPA with 30-fold higher activity than other orthologs such as from yeast and spinach, thus resulting in higher specific enzyme activity [23].

Pyruvate **2d** differs from hydroxypyruvate only by the absence of the C-3-hydroxyl group, and yet cannot function as a substrate in the transketolase reaction, indicating the critical role of the hydroxyl group in serving as a donor substrate [24]. Unlike TK, the closely related enzyme 1-deoxy-D-xylulose-5-phosphate synthase (DXS), catalyzes the transfer of the acetyl group from pyruvate **2d** to glyceraldehyde-3-phosphate to yield 1-deoxy-D-xylulose 5-phosphate. This was used recently as a template to guide the engineering of TK_gst_ for acceptance of pyruvate and aliphatic analogues, as the donor substrate [25]. However, this enzyme variant has not been shown to work also with the aliphatic or aromatic aldehydes as acceptor substrates. The best variants of TK_gst_ were able to catalyse the preparative scale reactions between novel donors and hydroxylated acceptors with isolated yields of 48 – 88 %. However, the acceptor substrate used during the process of directed evolution was the hydroxylated aldehyde, glycolaldehyde **1**. Therefore, as the wild-type TK already utilises hydroxylated acceptors with high activity, it remains unknown whether TK variants can be engineered to accept a combination of both pyruvate **2d** as donor, and aliphatic or aromatic aldehydes as acceptors. The reactions between pyruvate **2d** and aliphatic or aromatic aldehydes would allow access to a widely diverse range of products such as analogues of phenylacetylcarbinol (PAC), an important pharmaceutical intermediate, and precursors to drugs such as spisulosine and phenylpropanolamine.

In this study, we engineered *E. coli* TK variants that simultaneously accept both novel donor pyruvate **2d** and unnatural aliphatic and aromatic acceptor aldehydes, including propanal **5a**, pentanal **5b**, hexanal **5c** and 3-formylbenzoic acid (3-FBA **7**). Single variants capable of accepting pyruvate **2d** were firstly obtained using a structure-guided strategy and previous knowledge. A small but smart library was then constructed by combining the single mutants showing improved activity towards pyruvate **2d** and glycolaldehyde **1** or propionaldehyde **5a**. After screening this library, we successfully obtained the TK variants that could catalyse the reactions between sodium pyruvate **2d** and propionaldehyde **5a**, or 3-FBA **7**, with significantly improved efficiency compared to WT. The potential mechanisms for the enhanced activity was then evaluated using molecular docking simulations.

## Results and discussion

### Design of single mutants

Several enzymes are capable of accepting pyruvate **2d** for carboligation reactions among the superfamily of ThDP-dependent enzymes [26, 27]. For example, pyruvate decarboxylase (PDC) can catalyse a carboligation side reaction that has been exploited in the asymmetric synthesis of phenylacetyl carbinol from pyruvate **2d** and benzaldehyde. However, it also efficiently catalyses the decarboxylation of pyruvate **2d** into acetaldehyde and CO_2_ showing catalytic promiscuity [28, 29]. Therefore, we attempted to design variants of TK based on a structural alignment with PDC.

The *E. coli* TK crystal structure complexed with the natural donor substrate, D-xylulose-5-phosphate and cofactor ThDP has been solved previously [30]. Hydroxypyruvate readily acts as a donor substrate, whereas pyruvate 2d cannot, indicating a critical role of the C-3-hydroxyl group for catalysis. The TK structure in complex with the donor substrate showed that it is anchored by hydrogen bonds between the donor C-3-hydroxyl group, and His100 and His473 in the enzyme active site (Fig. 1A). Therefore, in order to stabilize new donor substrates lacking the C-3-hydroxyl group, new interactions may be required with other local residues, such as hydrophobic interactions with the methyl group in pyruvate. Therefore, we identified nine residues (His26, His66, His100, Leu116, Ile189, His261, Ser385, Asp469, His473) within 5 Å of the first two carbons of the donor substrate, as potential mutation sites (Fig. 1A).

**Fig. 1.**
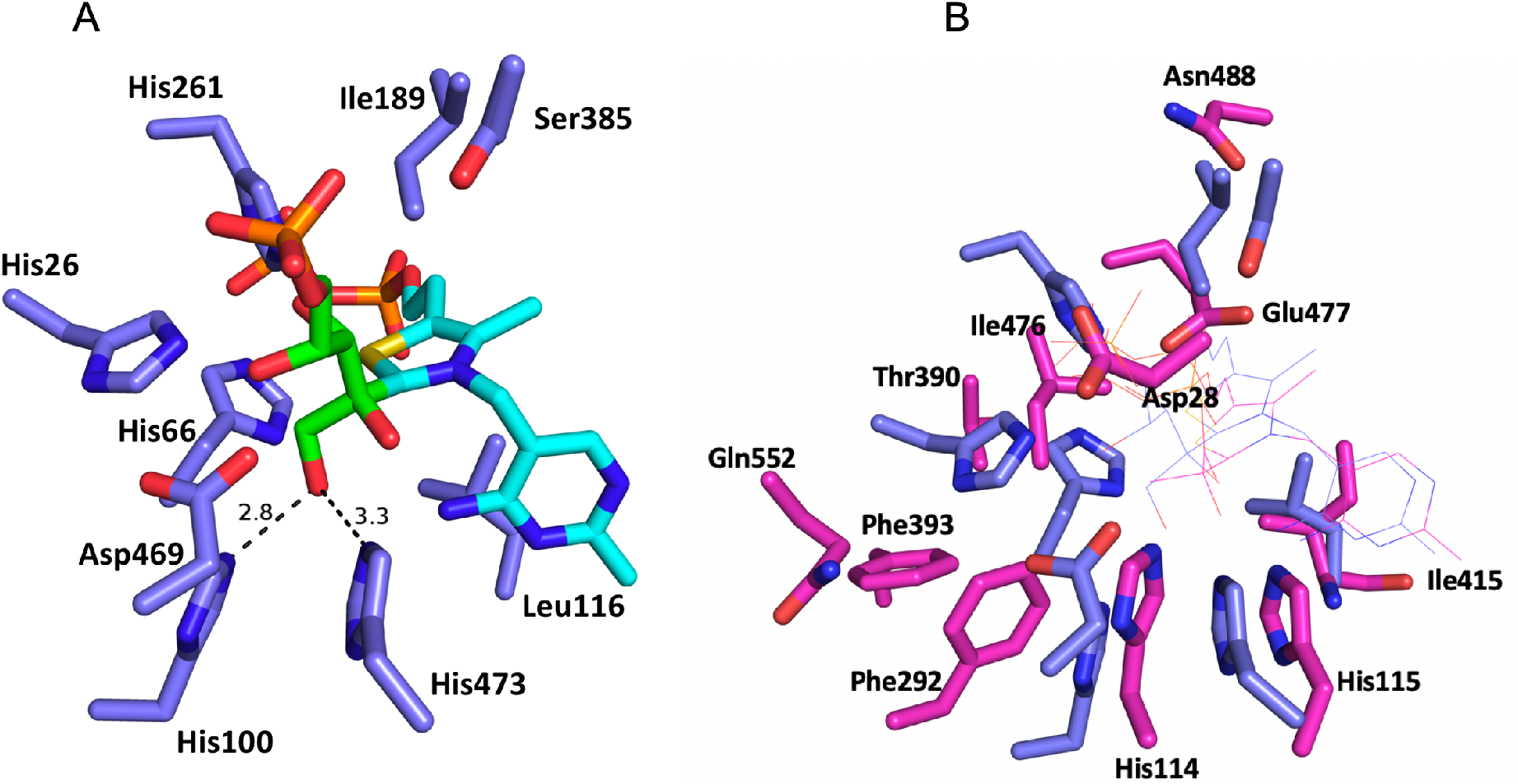
Structure alignment between transketolase (2R8O.pdb) and pyruvate decarboxylase (2VK1.pdb). A, Structure of TK in the complex with D-xylulose-5-phosphate: green, the substrate; cyan, thiamine pyrophosphate (ThDP). Distances are shown in Å. B, Active-site alignment between transketolase (blue) and pyruvate decarboxylase (pink). Only the residues of pyruvate decarboxylase are labelled. The cofactor ThDP of two enzymes are shown as lines.

Although the TK and PDC studied only shared 6.7% sequence identity, 559 pairs of residues that are structurally equivalent were found using the TopMatch tool (Fig. S1, Table S1). The root-mean-square error of the superposition for the structurally equivalent Cα atoms was 2.66 Å and the sequence identity in the equivalent regions increased to 12.3% (Table S1). Interestingly, only three target residues Leu116, Ser385 and His473 were located in the structurally equivalent regions (Fig. S1B). Based on structure alignment, Leu116 of TK is equivalent to Ile415 of PDC, and so the mutation L116I was proposed. However, for the Ser385 and His473, the equivalent structural alignment with PDC showed the same residues at the equivalent positions (Fig. S1B). Ser385 of TK is equivalent to Ser32 of PDC, which is close to Asp28 in both the sequence and 3-D structure (Fig. 1B). Asp28 in PDC was reported previously to play a critical role in the stabilization of the enzyme-bound cofactor, and in the stereochemical control of the decarboxylation reaction [31], and so the mutation S385D was proposed. His473 formed a hydrogen bond with the C-3-hydroxy group in TK and played a role in anchoring the donor substrate. The equivalent residue in TK_gst_ was His474, and the mutations H474S and H474N were previously found respectively to give a 6-fold and 3-fold improved activity towards glycolaldehyde **1** and pyruvate **2d**. Therefore, we included the corresponding mutations H473S and H473N in *E. coli* TK to investigate whether these mutations would also improve the activity of *E. coli* TK towards pyruvate.

As for the target residues that were not in structurally equivalent regions, we proposed a range of mutations at each target site based on the types of residues found in the equivalent spatial location in PDC to enhance the chances of success. The active site residues around the cofactor and donor substrate for the aligned TK and PDC, are shown in Fig. 1B. Ile189 of TK was located in a space occupied by Glu477 in PDC, and also close to Asn488, and so the mutations I189N/E were proposed (Fig. 1B). I189Q was also included as glutamine has a similar structure to asparagine. Ile476 is a conserved hydrophobic residue among PDCs from various organisms and plays an important role in substrate recognition and enantioselectivity [32]. Three target histidines 26, 66 and 261 were all within 2 Å of Ile476 in PDC, and so the mutations H26I, H66I and H261I were also proposed. In addition, the mutations H26Q, H66T and H261E were proposed approximately corresponding to residues Gln552, Thr390 and Glu477 of PDC.

His100 in TK also formed a hydrogen bond with the C-3-hydroxyl group of the donor substrate and its mutation to alanine previously had little effect on the *K*_m_ for acceptor substrates, but significantly increased that of the donor substrate, suggesting its important role in accepting the donor substrate [33]. Based on the structural alignment to PDC, His100 was close to residues His114 and Phe292 of PDC (Fig. 1B), and so we proposed the mutation H100F. Considering the importance of this site, we also included alternative mutations. H100Y was included as tyrosine has a similar structure to phenylalanine. In addition, the TK_gst_ mutations H102F/L/T previously showed improved performance towards glycolaldehyde **1** and sodium pyruvate **2d** [25]. Therefore, we included corresponding mutations H100L and H100T in *E. coli* TK to investigate whether these mutations would improve activity in *E. coli* TK towards glycolaldehyde **1** or propionaldehyde **5a**, and pyruvate **2d**.

When predicting the potential mutations, previous knowledge was also considered. Asp469 is a hotspot for engineering TK towards novel acceptor substrates. While using HPA as the donor, D469E led previously to increased activity towards aliphatic aldehyde acceptors, and D469T increased the activity towards both aliphatic and aromatic aldehydes [17, 34]. Preliminary work suggested that these two variants may have accepted pyruvate **2d** as donor substrates with pentanal as the aldehyde acceptor, although with a very low reaction yield. Therefore, in addition to the D469H mutation suggested by the proximity of His114 in the structurally aligned PDC, we also included D469E and D469T into our screens. In total, we designed and constructed 20 single mutants (summarised in Table S2), and measured their activities towards glycolaldehyde **1** or propionaldehyde **5a** and pyruvate **2d** experimentally.

### Characterization of single variants

#### Reaction with glycolaldehyde and pyruvate

Clarified lysate preparations of the twenty single mutants were first tested for catalytic efficiency towards pyruvate **2d**, paired with glycolaldehyde **1** as this is known already to be one of the best acceptor substrates for wild-type TK. This allowed us to evaluate the acceptance of the novel donor substrate in the context of a favourable hydroxylated aldehyde acceptor. The target product, 3,4-dihydroxy-2-butanone (DHB) 3 of the reaction between glycolaldehyde **1** and pyruvate **2d** was detected by HPLC at a retention time of 13.7 min (Scheme 1). However, erythrulose **4** was also detected as a side product at the retention time of 11.4 min (Scheme 1, Fig. 2A), which was not reported in any previous mutagenesis studies, although the reaction of two glycolaldehydes to form erythrulose is a known slow side reaction for wild-type TK [35].

**Fig. 2.**
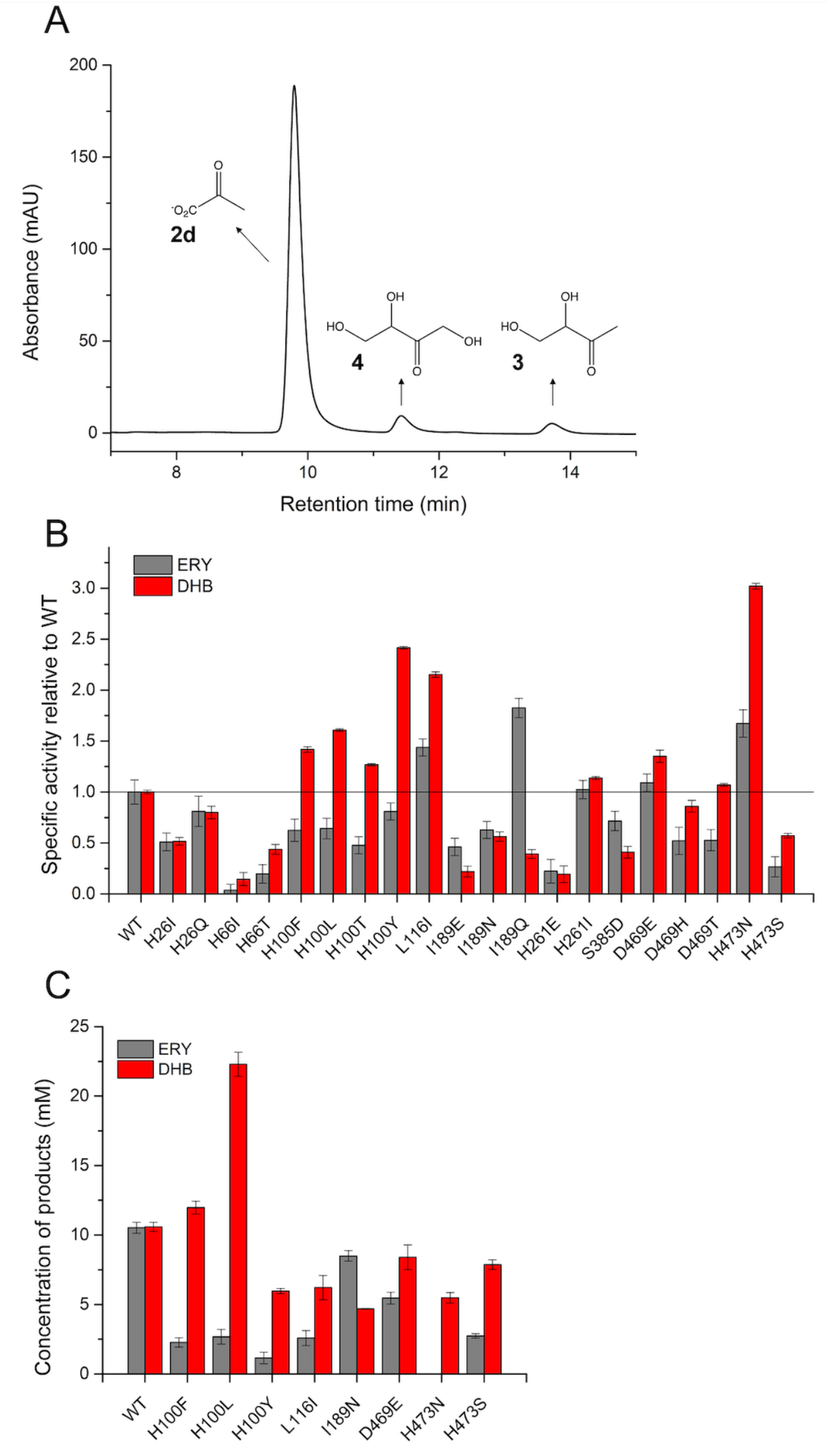
Mutants catalysing the reaction between glycolaldehyde **1** and sodium pyruvate **2d**. A, HPLC chromatography of the reaction glycolaldehyde **1** and sodium pyruvate **2d**. Retention time for erythrulose (ERY) **4** and 3,4-dihydroxy-2-butanone (DHB) **3** were 11.4 min and 13.7 min, respectively. B, Specific activity (yield at 24 h normalised to TK concentration) of TK variant lysates, relative to WT, for reaction of glycolaldehyde **1** and sodium pyruvate **2d**. C, Final concentrations of the two products for the 24-h reactions catalysed by the purified variants. Enzyme reactions were carried out at 50 mM glycolaldehyde **1**, 50 mM sodium pyruvate **2d**, 50 mM Tris-HCl, pH 7.0, and 0.13 mg/mL purified enzyme or enzyme from 1.3 mg/mL lysate, in triplicate vials, at 30 °C for 24 h.

Considering the possible difference in expression level of the variants, the yields of both products determined after 24-h reaction, were normalised by the TK concentration as determined from the relative density of SDS-PAGE bands, to give an approximate measure of specific activity (Fig. S2). Each of the mutations tested influenced the yields of both the target product and the erythrulose byproduct, and also the ratio of the two products, relative to WT (Fig. 2B). The variants H100F/L/T/Y, L116I, D469E and H473N in particular, gave markedly increased reaction yields of the target product relative to that of WT. All of these improved variants were subsequently purified and re-evaluated, except for H100T which was the lowest ranked. Two negative variants I189N, H473S were also purified and tested as controls (Fig. 2C). Interestingly, H100Y, L116I, D469E and H473N no longer showed an improvement in activity in their pure enzyme form, compared to wild type (WT) (Fig. 2B). The same variants also had low protein expression levels (Fig. S2) and so their poor specific activity when purified may have been due to an adverse impact of the mutations on enzyme stability, or otherwise resulted from the increased error in measuring low protein concentrations by densitometry on lysates.

The yield of reaction catalysed by H100L was 22.3 mM (45%) (Table 1), a 2.1-fold improvement compared to WT, consistent with previous observations with equivalent TK_gst_ variants [25]. H100F also improved the catalytic efficiency by 1.2-fold. However, H473N and H473S did not improve the target product yield in *E. coli* TK, in contrast to the effect of the equivalent mutations in TK_gst_ (Fig. 2C). The contextual structure differences between the active sites of *E. coli* TK and TK_gst_ are a likely reason for these differences. However, it should be noted that differences in the activity assays used in our work compared to that in previous reports with TKgst, could also lead to similar observations. The pH-sensitive colorimetric assay used for TK_gst_ would not have detected the erythrulose byproduct, nor distinguish between the decarboxylation of pyruvate **2d** versus true transketolase function (decarboxylation followed by ketone unit transfer to acceptor) [25, 36].

**Table 1.**
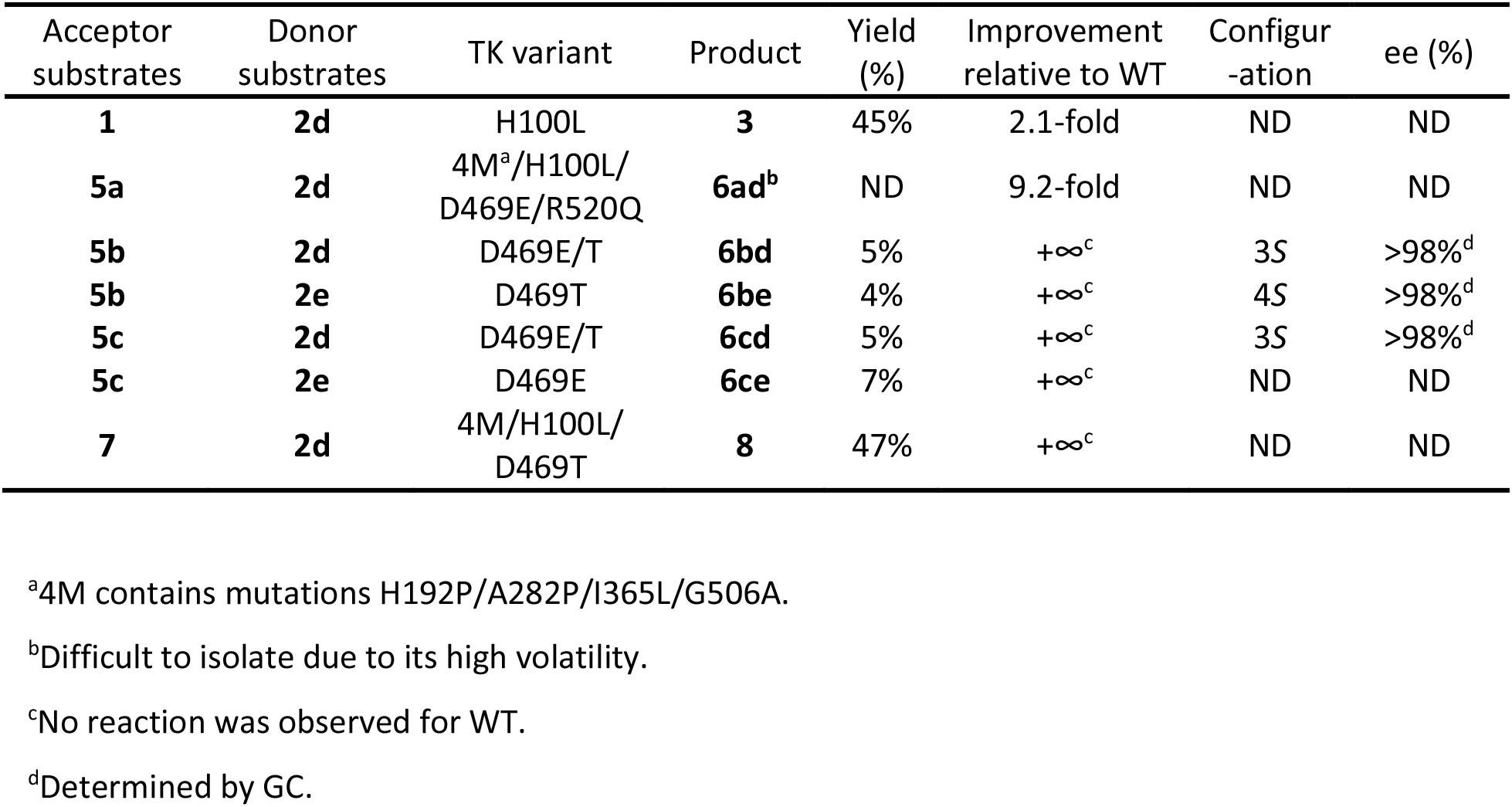
Reactions between novel acceptor and donor substrates catalysed by TK variants

The variants also changed the relative reaction yields for the two products. WT produced equal concentrations of target product and byproduct, at 10.5 mM (21%), whereas all purified variants except I189N, gave higher yields of the target product DHB **3**, than the byproduct erythrulose **4**. H473N gave the highest selectivity, with no byproduct detected after 24-h reaction, even though the target product accumulated to 5.5 mM. The ratios of the target product DHB **3** relative to byproduct were 8.3, 5.3, 5.2, 2.9, 2.4, and 1.5, for H100L, H100F, H110Y, H473S, L116I, and D469E, respectively.

Most of the variants based on structural comparison to PDC, failed to improve the yield of the target product 3,4-dihydroxy-2-butanone **3** for the reaction between glycolaldehyde **1** and pyruvate **2d**, which is perhaps not surprising given only 6.7% sequence identity between TK and PDC, and the likely need for multiple simultaneous mutations to generate a change in function.

#### Reaction with propionaldehyde and pyruvate and extension to other aliphatic analogues to determine stereoselectivities

All variants were also tested for their activities towards propionaldehyde **5a** and sodium pyruvate **2d**. Wild-type *E. coli* TK can accept a range of non-hydroxylated aliphatic aldehyde substrates in the presence of hydroxypyruvate. However, the activity is typically only 5–35% of those for hydroxylated substrates such as glycolaldehyde **1**. Propionaldehyde **5a** is therefore a more challenging acceptor for TK than glycolaldehyde **1**, especially when using pyruvate 2d as the donor substrate. Initially the 12 variants not already purified above (Fig. 2C), were first tested for their activity in lysates, to identify any to add into the pool of purified variants. After reaction for 24 h using the cell lysates (Fig. 3A), the target product 3-hydroxy-2-pentanone **6ad** was detected indirectly using the colorimetric assay. Four variants, H26Q, H100T, H261I and D469T gave 2-to 4-fold improved specific activities relative to WT (Fig. 3A). These four variants were subsequently purified and re-evaluated together with the other 8 variants already purified above.

**Fig. 3.**
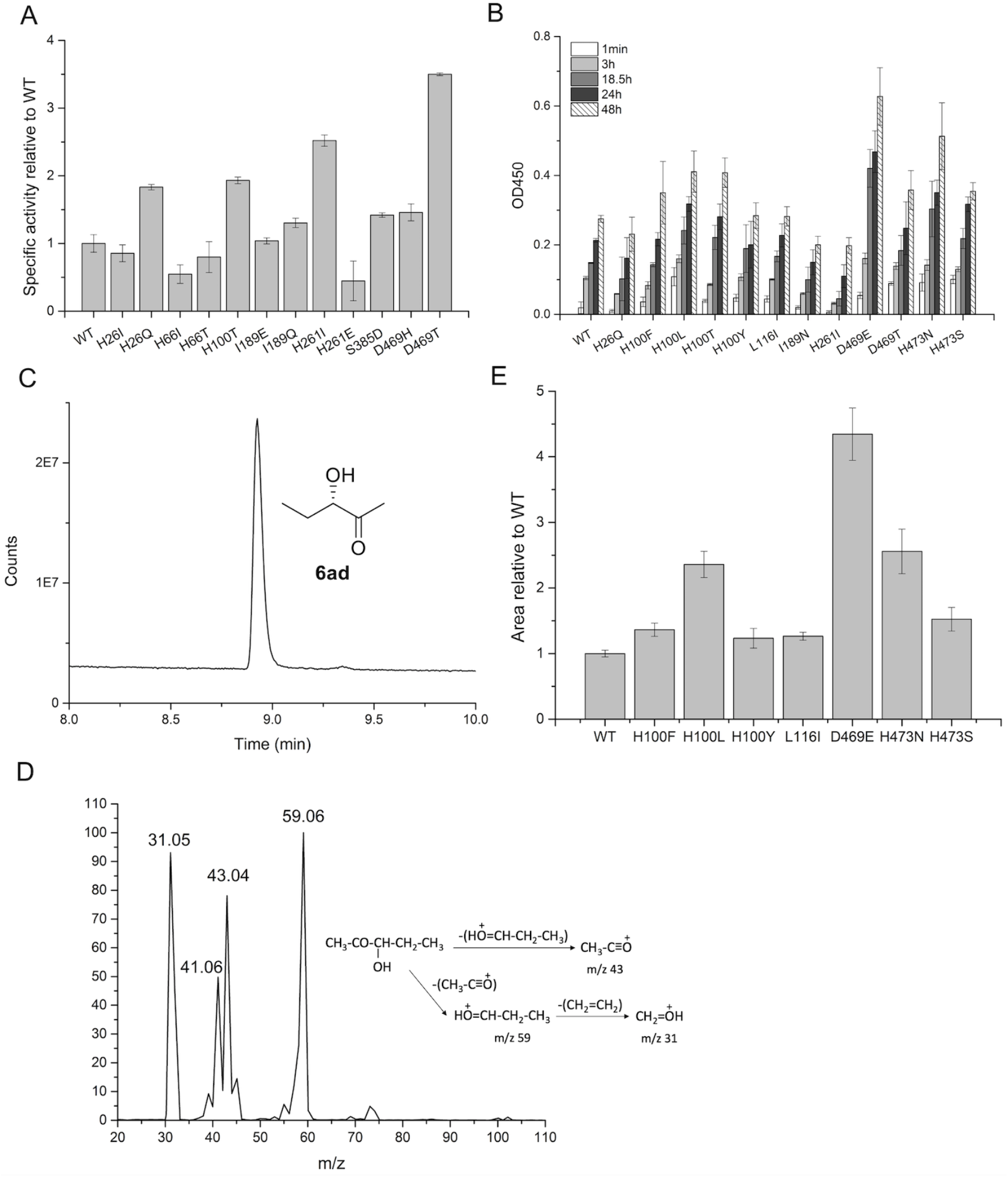
Activity of TK variants towards propionaldehyde **5a** and sodium pyruvate **2d**. (A), Specific activity (yield at 24 h normalised to TK concentration) of TK variant lysates, relative to WT, for reaction of propionaldehyde **5a** and sodium pyruvate **2d**. (B), the yield of enzyme reactions catalysed by purified proteins based on the colorimetric assay. (C), the peak of 3-hydroxy-2-pentanone **6ad** identified by GC for WT. (D), mass spectrum of the 3-hydroxy-2-pentanone **6ad** GC peak fraction. (E), yield of 48-h reaction quantified based on GC peak area. Enzyme reactions were carried out at 50 mM propionaldehyde **5a**, 50 mM sodium pyruvate **2d**, 50 mM Tris-HCl, pH 7.0, and 0.13 mg/mL purified enzyme or enzyme from 1.3 mg/mL lysate, in triplicate vials, at 30 °C for 24 h or 48 h.

The absorbance at 450 nm increased with reaction time indicating a significant time-dependent formation of 3-hydroxy-2-pentanone **6ad** (Fig. 3B). However, the purified variants H26Q and H261I no longer improved the activity, even though their lysates had shown improved performance, which could also be attributed to their low expression as discussed above. Seven variants including H100F, H100L, H100T, D469E, D469T, H473N and H474S gave improved activity relative to WT, with the most active variant D469E having more than 2-fold improvement in the catalytic efficiency compared to WT (Fig. 3B).

Based on the colorimetric assay, WT also appeared to show activity towards propionaldehyde **5a** and pyruvate **2d**, which had never been observed previously. In order to confirm the reaction product, GC-MS was applied to analyse the reaction mixture after 24-hours reaction. A peak was identified by GC with the retention time of 8.9 min (Fig. 3C). After ionization, the compound 3-hydroxy-2-pentanone **6ad** revealed three expected fragments, having mass-to-charge ratios of 43, 59, and 31, respectively, indicating the successful generation of the target product (Fig. 3D).

The peak area in GC was used to quantify the relative amounts of product generated by the enzyme reactions. We carried out the enzyme reactions for 48 hours using seven variants, including both the improved and non-improved ones in their purified form. The GC peak area and OD450 gave a good linear correlation with an R^2^ of 0.93, reflecting that the colorimetric assay was a reliable activity measurement method (Fig. S3). The negative intercept on the GC peak area axis indicated that some of the colorimetric assay response was due to background. This would reduce the apparent activity of WT considerably, although the presence of a peak by GC confirmed that it could indeed form the target product. Consistent with the measurements of the colorimetric assay, the GC peak area of the 3-hydroxy-2-pentanone **6ad** showed that D469E was the most active variant, with a 4.3-fold improvement in yield relative to that of WT. H100F, H473S, H100L and H473N also revealed 1.4-fold, 1.5-fold, 2.4-fold and 2.6-fold improvements in yield respectively, relative to WT (Fig. 3E). Larger improvements were observed by GC relative to WT, as a result of the better removal of background signal compared to in the colorimetric assay, and so the results from GC would be more accurate.

3-Hydroxy-2-pentanone **6ad** is hard to isolate due to its volatility so larger scale TK reactions were performed with pentanal **5b** and hexanal **5c** as acceptors with a view to isolating the products. Both pyruvate **2d** and keto-butyric acid **2e** were also used as donors to highlight general applicabilities of the reaction at this stage (Scheme 2). Following the screening of variants with propanal and pyruvate described above, D469E was selected for use together with D469T as a lower performing mutant for comparison purposes. As shown in Scheme 2, isolated products **6bd** and **6cd** were formed in reactions with both D469T and D469E in both cases in approximately 5% yield, although some material was lost due to volatility (Table 1). (3R)-Hydroxy-2-octanone **6cd**, the product arising from hexanal **5c** and **2d**, is an insect pheromone produced by males of the longhorn beetle *Anaglyptus subfasciatus* and a Mosher’s derivatisation method has been reported to determine enantiomeric excesses (ees) [37]. Following this precedent the TK product **6cd** was reacted with (2*R*)-3,3,3-trifluoro-2-phenylpropanoyl chloride ((*R*)-MTPA) to give the (2S)-Mosher’s ester. ^1^H NMR analysis revealed a major singlet for the 1-H_3_ group at 2.21 ppm, and a minor peak at 2.13 ppm. The positions of these two signals corresponded to the (2S,3S) and (2S,3R)-esters, respectively, following the literature and indicated that the major (3S)-isomer was formed by the TK mutants: integration of the peaks revealed an ee of >95%. A second set of diastereomeric signals could also be observed for the 8-H_3_ triplet, and integration of the signals at 0.83 ppm (major (2*S*,3*S*)-isomer) and 0.93 ppm (minor (2*S*,3*R*)-isomer) confirmed the *ee* as >95%. The absolute stereochemistry was further confirmed by measurement of the optical rotation compared to reported data (see SI). In addition, racemic samples of **6bd** and **6cd** were also formed via chemical methods as chemical standards (see SI) to determine the isomeric compositions of the products via chiral GC which confirmed the formation of the major isomer in >98% *ee* (Table 1). The reaction product **6bd** formed from pentanal **5b** and pyruvate **2d** was then derivatised using (*R*)-MTPA, and integration of the 1-H_3_ signal again indicated an *ee* of >95% for the (S)-isomer. Chiral GC of the racemic product compared to the TK products with D469T and D469E similarly revealed ees of >98%. These results highlighted the highly stereoselective nature of the reactions.

From these results the same aldehydes were used with keto-butyric acid 2e to give **6**be and **6ce** using D469T and D469E respectively, which gave the highest isolated yields. Racemic samples of **6be** and **6ce** were again prepared to confirm the characterisation data. To determine again the stereoselectivity for one example **6be**, it was analysed by chiral GC and reacted again with (*R*)-MTPA. Analysis of the ester product 8-H_3_ signal revealed two triplets at 0.82 ppm (major (2*S*,4*S*)-isomer) and 0.94 ppm (minor (2*S*,4*R*)-isomer), consistent with the method described above, with integrations indicating the *ee* as >95%, and GC analysis gave an *ee* of >98% (see SI).

The improvements in yield for the reaction between propionaldehyde **5a** and pyruvate **2d** from *E. coli* TK variants D469E, H473N, H100L, H473S and H100F, were particularly interesting given that they typically improved the activity towards glycolaldehyde **1** and sodium pyruvate **2d** by a lesser extent, or even not at all for D469E, H473N and H473S. This highlights the complexity of engineering an enzyme towards two substrates simultaneously, and also that improvements towards the more hydrophobic pyruvate (relative to hydroxypyruvate), may be structurally linked to the better acceptance of the more hydrophobic aldehyde acceptor propionaldehyde **5a** (relative to glycolaldehyde 1).

The catalytic reaction of TK proceeds in two stages. In the first half of the reaction, cleavage of the donor substrate is accompanied with the formation of a covalent intermediate. In the second half of the reaction, nucleophilic attack of the acceptor substrate by the enamine intermediate leads to product formation and release. The improvement of the final yield of the reaction could be attributed to both the first and second halves of the reaction. D469E showed improved reaction yields towards propionaldehyde **5a** and pyruvate **2d**, and yet failed to significantly increase the activity towards glycolaldehyde **1** and pyruvate **2d**. Interestingly, D469E also showed improved activity towards propionaldehyde **5a** but decreased the activity towards glycolaldehyde **1** when HPA was used as the donor [13, 16, 36]. This suggested that D469E only improved the efficiency of the second half of the reaction which involves acceptance of the aldehyde substrate, but did not affect acceptance of pyruvate in the first half. By contrast, the mutations of His100 could have only influenced the first half of the reaction, accepting the pyruvate. We tested the correlation between reaction yields for the purified variants, when reacting with the two different acceptors and sodium pyruvate (Fig. S4). It clearly showed that the mutations of residue His100 were responsible for the enzyme activity towards both the glycolaldehyde and propanal, with an R^2^ of 0.99 when only H100L, H100F, H100Y and WT were considered, suggesting that mutations of His100 were critically required to accept the pyruvate, while not being important for controlling the acceptor specificity, consistent with previous observations [33] (Fig. S4). We thus combined the variants that improved donor acceptance with those that improved acceptor acceptance, to generate a new library.

### Design of ‘a small but smart’ library

In order to further improve the activity of *E. coli* TK towards propionaldehyde **5a** and pyruvate **2d**, eight single variants including H100F, H100L, H100T, D469E, D469T, H473N, H473S and R520Q were selected for recombination to construct the library. Among them, H100F, H100L, H100T, D469E, D469T, H473N, H473S each showed improvement in the activity towards propionaldehyde **5a** and sodium pyruvate **2d**. The R520Q mutation was also included as it was previously found to improve stability, and also the catalytic efficiency towards HPA and propionaldehyde **5a**, within certain other variants [34]. This strategy kept the library size small while still sampling for potential cooperative effects between mutations. In previous work with TK [34], recombination of single active-site mutants, including at Asp469, nearly always led to the loss of soluble expression. Therefore, we used a previously stabilised variant H192P/A282P/I365L/G506A (4M) as the template for constructing the new library to promote evolvability [34, 38, 39]. Introduction of these same stabilising mutations was recently found to restore the stability loss of another TK variant (S385D/D469T/R520Q) that was engineered to accepted aromatic aldehydes [39, 40].

### Improvement of activity towards propionaldehyde and pyruvate

The ‘small but smart library’ was screened towards propionaldehyde **5a** and sodium pyruvate **2d** using the colorimetric assay. Ten promising variants were identified, purified and tested for catalytic activity. The reaction yield relative to WT was determined for the variants using GC after reaction for 24 h (Fig. 4). As the quadruple variant H192P/A282P/I365L/G506A (4M) was used as the template for constructing the library, its activity was also measured. The 4M maintained a comparable activity with WT, whereas its combination with other mutations influenced their impact. The D469E variant previously gave a 4.3-fold improvement in product yield relative to the WT, whereas 4M/D469E had improved the yield by 6.7-fold relative to 4M, demonstrating the benefit of TK stabilisation even for the single active-site mutations (Fig. 3E & 4). However, the addition of the compensatory stabilising 4M mutations did not significantly change the impact of H100L (Fig. 3E & 4), presumably as this variant did not significantly decrease the stability of TK relative to the WT stability. Interestingly, addition of 4M influenced the activity of H473N in a negative way. 4M/H473N only showed 1.2-fold improvement in reaction yield compared to WT, whereas the H473N itself revealed 2.4-fold improvement. Thus, the relationship between template stability and the fitness of the new variant is not always predictable.

**Fig. 4.**
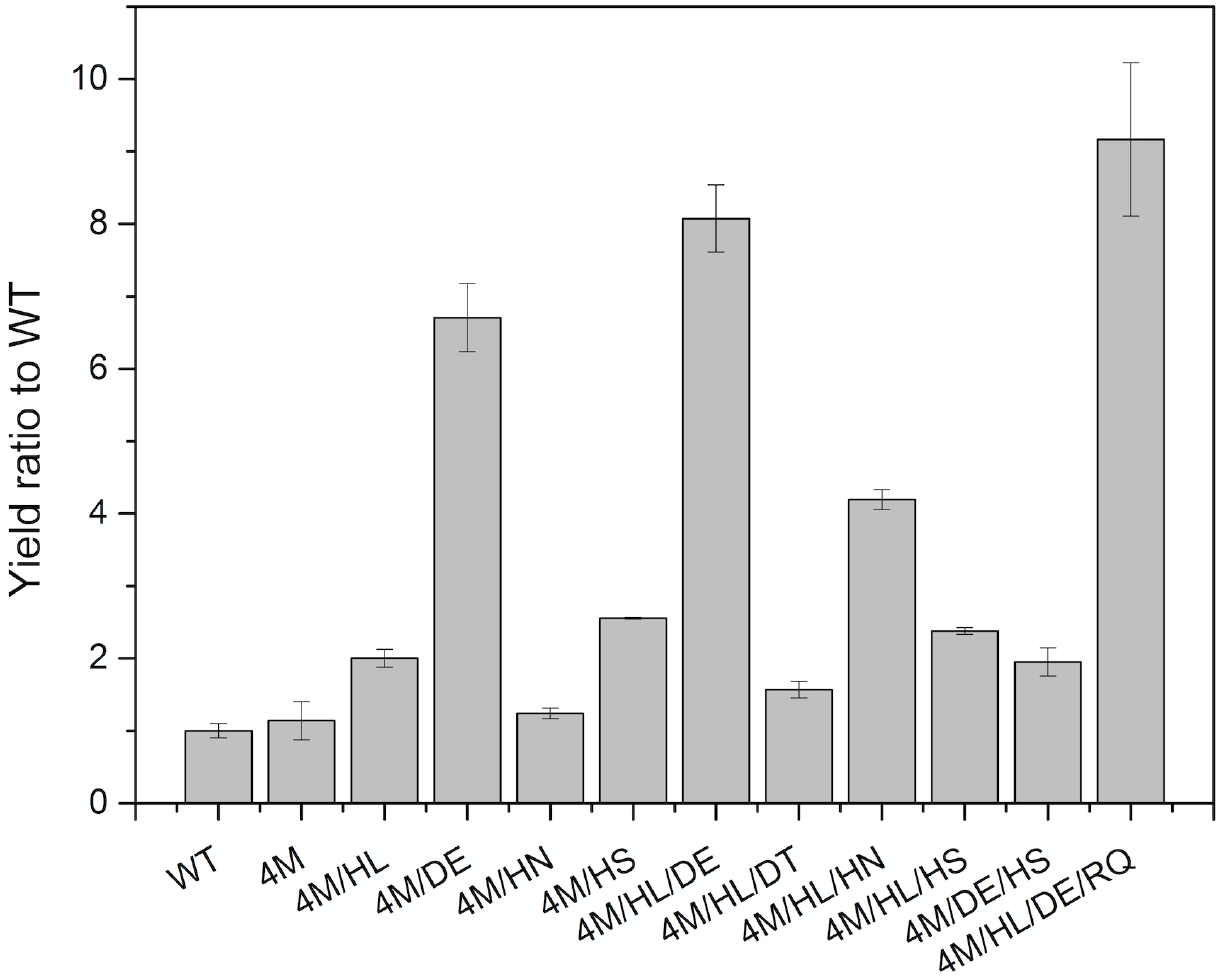
Yield of reactions towards propionaldehyde and pyruvate, catalysed by purified TK variants for 24 h. The yield was calculated based on GC area. 4M is the quadruple mutant H192P/A282P/I365L/G506A. HL, H100L; DE, D469E; HN, H473N; HS, H473S; RQ, R520Q. The catalytic activity was tested towards 50 mM propionaldehyde **5a** and 50 mM sodium pyruvate **2d** in 50 mM Tris-HCl solution at 30 °C. The enzyme concentration used was 0.13 mg/mL.

The combination between H100L and D469E generated the variant 4M/HL/DE that gave an 8.1-fold improvement in the reaction yield compared to WT. Addition of the mutation R520Q to this variant improved the reaction yield further and generated the best variant overall, 4M/HL/DE/RQ, which had a 9.2-fold improvement in the reaction yield compared to WT (Fig. 4, Table 1). R520Q has been reported to increase the activity of D469T towards propionaldehyde 5a and HPA due to improved soluble protein expression. The combination of H100L and H473N also led to a variant showing 4.2-fold improvement in the reaction yield compared to the WT, confirming the important role of H100L in accepting pyruvate **2d**.

### Novel enzyme activity towards 3-FBA and pyruvate

In our previous work, variants showing improved performance towards propionaldehyde **5a** and HPA also improved the catalytic efficiency towards aromatic substrates with HPA, including 3-FBA **7** or 4-FBA [17, 18]. For instance, D469T improved the specific activity towards propionaldehyde **5a** and HPA by 5-fold and also improved conversion yield of 3-FBA **7** and 4-FBA from 0% to 65% and 30% respectively. We therefore explored whether any of the new library variants had improved activity towards the aromatic aldehyde 3-FBA 7 and pyruvate **2d** (Scheme 3).

The whole library was screened for activity towards 3-FBA **7** and sodium pyruvate **2d** as determined by HPLC. Most variants had no activity towards these two substrates, with the exception of 4M/H100L/D469T, 4M/D469T/H473N, and 4M/D469T/H473N/R520Q. After reaction for 24 h, a new peak at the retention time of 4.1 min was identified that correlated with the consumption of the 3-FBA **7** in the HPLC chromatogram for these three variants, but not for WT or any other variants (Fig. 5A).

**Fig. 5.**
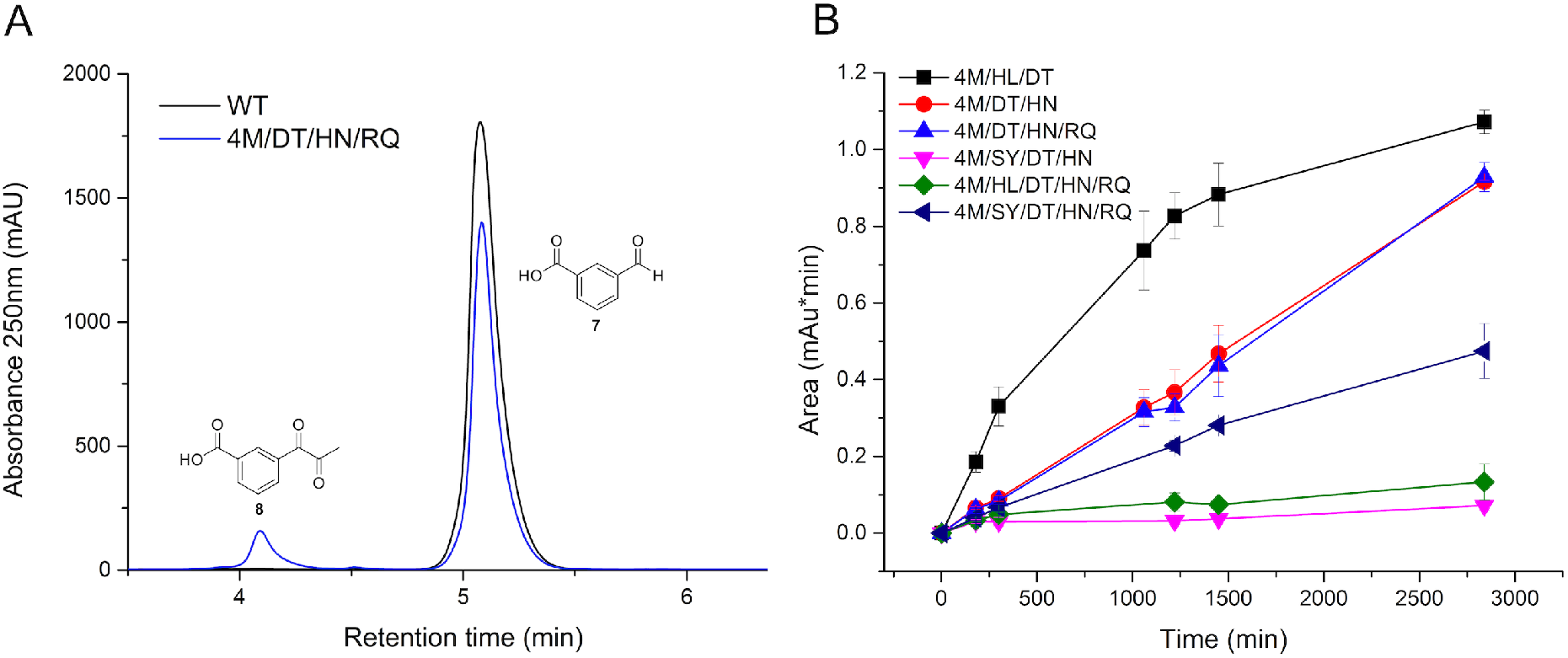
Catalytic reaction between 3-FBA 7 and sodium pyruvate **2d**. A, HPLC chromatograms of reaction between 3-FBA **7** and sodium pyruvate **2d**. B, Catalytic activity of TK variants. 4M is the quadruple mutant H192P/A282P/I365L/G506A. HL, H100L; SY, S385Y; HN, H473N; DT, D469T; RQ, R520Q. The enzyme activity was tested towards 50 mM 3-FBA 7 and 50 mM sodium pyruvate **2d** in 50 mM Tris-HCl solution at 30 °C. And the enzyme concentration used was 0.067 mg/mL.

In order to confirm if this peak was the product expected, the peak was isolated by semipreparative HPLC and characterized by ^1^H NMR and LC-MS. In the LC-MS analysis of the extracts, a peak with an m/z of 195.1 was observed, corresponding to the expected product 3-(1-hydroxy-2-oxopropyl) benzoic acid **8** (Fig. S5A). The NMR spectrum also confirmed the generation of the target product (See SI). These data demonstrated that the *E. coli* TK variants catalysed the unprecedented reaction between the aromatic aldehyde and sodium pyruvate **2d**.

Three variants including 4M/H100L/D469T (4M/HL/DT), 4M/D469T/H473N (4M/HL/DT/HN), 4M/D469T/H473N/R520Q (4M/HL/DT/RQ) were also purified and tested for the conversion of 3-FBA **7** with sodium pyruvate **2d** as the donor substrate. The variant 4M/H100L/D469T showed the highest activity of all, with an initial rate around 2-fold higher than those of 4M/D469T/H473N and 4M/D469T/H473N/R520Q (Fig. 5B). The variant 4M/H100L/D469T was then characterized at higher enzyme concentrations. When increasing the concentration of enzyme from 0.067 mg/mL to 1.33 mg/mL, the conversion from 3-FBA 7 to product 3-(1-hydroxy-2-oxopropyl) benzoic acid 8 within 24 h was increased from 2.5% to 47% based on the substrate depletion (Table 1 & S3). The specific activity, *K*_m_ and *k*_cat_ of 4M/H100L/D469T were also found to be 0.048 μmol mg^-1^ min^-1^, 90.4 mM and 0.22 s^-1^ respectively (Table S3).

Addition of the mutation S385Y into the double mutant D469T/R520Q previously improved the catalytic efficiency towards 3-FBA **7** and HPA by significantly decreasing the Km [18]. We therefore introduced the S385Y mutation into the two mutants 4M/D469T/H473N and 4M/D469T/H473N/R520Q to test whether their activities could be improved. The variant 4M/H100L/D469T/H473N/R520Q was also purified for activity measurements. Interestingly, the mutation S385Y failed to improve the activity for the variants of 4M/D469T/H473N and 4M/D469T/H473N/R520Q (Fig. 5B). The residue Ser385 was within 5 Å around the first two carbons of the donor substrate (Fig. 1A). The mutation S385Y, hence, could easily have interfered with the binding of pyruvate **2d** and decreased the activity towards 3-FBA **7** and sodium pyruvate **2d**. The variant 4M/H100L/D469T/H473N/R520Q also showed limited activity, consistent with the activity measured using lysates (Fig. 5B).

### Elucidating the mechanism of enhanced activity

To gain insights into the influence of the mutations on enzyme activity, the substrates were docked into the binding pocket of the WT and several TK variants using the Autodock Vina program [41]. 4M/H100L/D469E and 4M/H100L/D469T were selected for docking as they had significantly improved enzyme catalytic efficiencies and yet different activities towards propionaldehyde **5a** and 3-FBA **7** with the only difference in their substitution of position 469. The first half of the TK reaction consists of the cleavage of the donor substrate, pyruvate **2d** and the formation of the first product, carbon dioxide and a covalent intermediate, ThDP-enamine. The intermediate was first docked into wild-type and mutant TKs (Fig. S6A). ThDP-enamine had the binding affinity energy of −9.5 kcal mol^-1^ when bound into the WT, 0.3 kcal mol^-1^ higher than when it was bound into the 4M/HL/DE and 4M/HL/DT, indicating that the intermediate had stronger binding with the variants than with the WT. The pose of ThDP-enamine was similar in the binding pockets of 4M/HL/DE and 4M/HL/DT with a RMSD of 0.3 Å, and yet slightly to the binding in WT with a RMSD of 0. 7 Å (Fig. S6A, Table 2). When the pyruvate **2d** instead of hydroxypyruvate was used as the donor substrate, the (α-hydroxyethyl)-ThDP intermediate instead of (α, β-dihydroxyethyl)-ThDP intermediate was generated. In the WT, the C2 methyl group of the hydroxyethyl was located in a highly polar environment surrounded by six charged residues including H26, H66, H100, H261, D469 and H473, which was potentially unfavourable (Fig. S6B). Substitution of histidine with leucine at the position 100 would reduce the polarity of this local region and form a hydrophobic interaction between L100 and C2 methyl group of the intermediate (Fig. S6C&D). This could result in the slight difference of binding orientation of ThDP-enamine, and stronger binding affinity in the two variants compared to WT.

**Table 2.** Substrates binding pose and affinity in wild-type and mutant TKs

|  | ThDP-enamine |  |  | Propionaldehyde |  |  |  | 3-FBA |  |  |  |
| --- | --- | --- | --- | --- | --- | --- | --- | --- | --- | --- | --- |
|  | Affinity (kcal mol<sup>-1</sup>) | RMSD<sup>[a]</sup> (Å) |  | Affinity (kcal mol<sup>-1</sup>) | Distance<sup>a</sup> (Å) | RMSD (Å) |  | Affinity (kcal mol<sup>-1</sup>) | Distance<sup>a</sup> (Å) | RMSD (Å) |  |
| WT | -9.5 | 0 | 0.7 | -2.4 | 5.2 | 0 | 2.0 | -5.4 | 5.0 | 0 | 1.1 |
| 4M/HL/DE | -9.8 | 0.7 | 0 | -2.4 | 4.6 | 2.0 | 0 | -5.6 | 6.1 | 1.1 | 0 |
| 4M/HL/DT | -9.8 | 0.7 | 0.3 | -2.4 | 5.0 | 0.9 | 2.4 | -5.5 | 5.1 | 2.1 | 2.5 |

<sup>a</sup>The RMSD was calculated with WT or 4M/HL/DE as the reference
<sup>b</sup>The distance is the nucleophilic attack distance of α-carbanion of the ThDP-enamine on the aldehyde group of the acceptor substrate

Two acceptor substrates, propionaldehyde **5a** and 3-FBA **7** were then separately docked into the WT and variants containing the ThDP-enamine intermediate. According to the elucidated catalysis mechanisms of TK, the α-carbanion of the (α-hydroxyethyl)-ThDP intermediate would attack the aldehyde group of the acceptor substrates to form the products with a carbon skeleton extended by two carbon atoms. The distances between the C-1 atom of the hydroxyethyl group of the ThDP-enamine and the carbon atom of the aldehyde group of propionaldehyde **5a** were 5.2 Å, 4.6 Å and 5.0 Å for the WT, 4M/HL/DE and 4M/HL/DT respectively, consistent with the activity measurements towards propionaldehyde **5a** and pyruvate **2d** (Table 2, Fig. 4 & 6). The higher activity of 4M/HL/DE thus potentially resulted from the slightly shorter distance between the acceptor substrate propionaldehyde **5a**, and the enamine intermediate, compared to 4M/HL/DT and WT. As shown in Fig. 6 A-C, the D469T and D469E mutations apparently led to differences in the shape of the tunnel around the intermediate. The D469E mutation led to a narrower tunnel because of the longer side chain of glutamic acid. The longer side chain also has greater steric hindrance, which result in C-2, C-3 atoms of propanal have dramatically different positions when docked into WT, 4M/HL/DE and 4M/HL/DT (Fig. S7). Compared to the side chain of threonine, that of glutamic acid will have greater polarity, whereas the C-2, C-3 atoms of propanal are non-polar. The strong polarity and long side chain of Glu469, hence, may lead propanal to have a shorter distance towards the ThDP-enamine compared to WT and 4M/HL/DT (Table 2 Fig. S7).

**Fig. 6.**
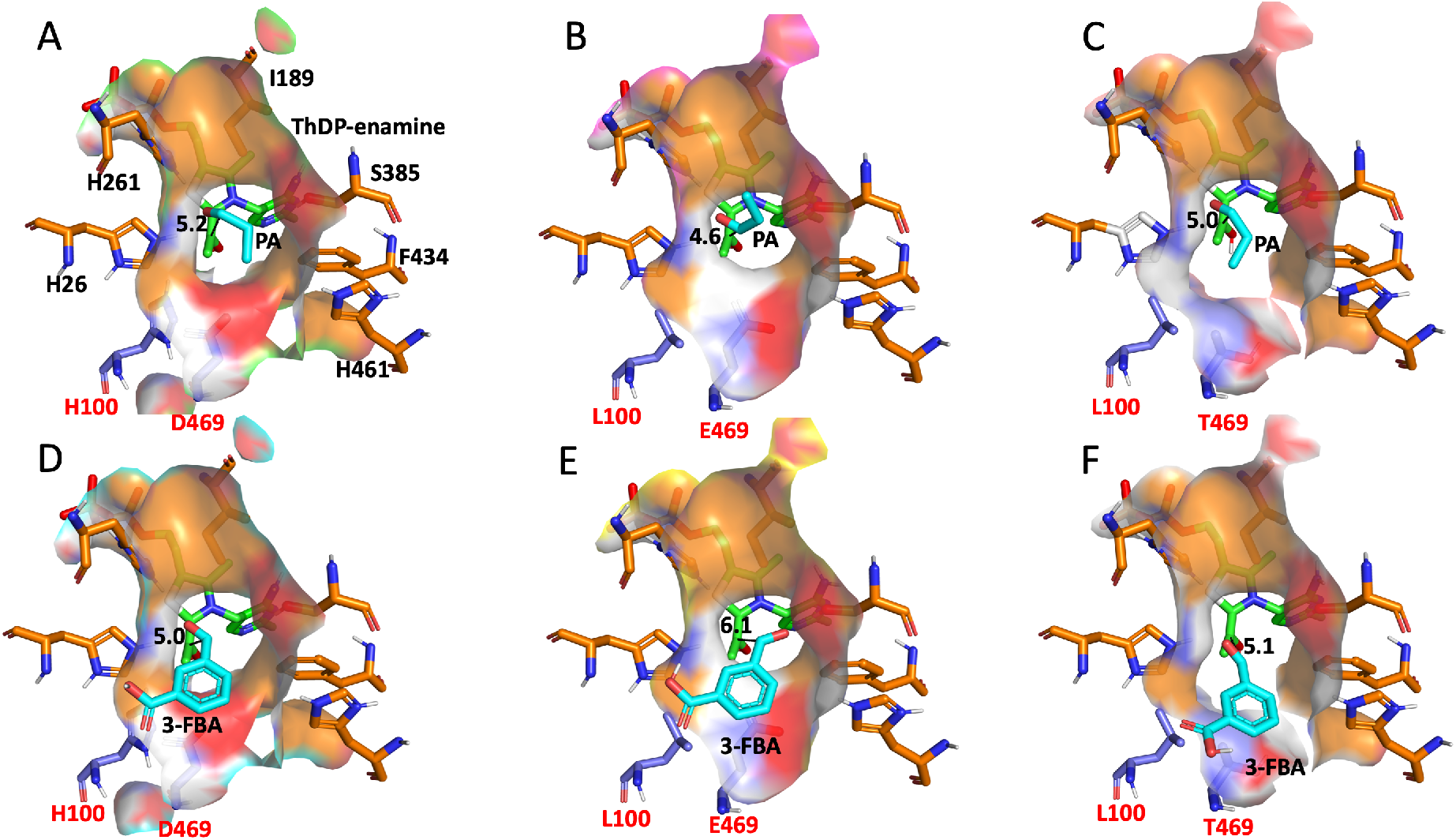
Binding mode analysis of substrates in wild-type and mutant TKs. A, Docking of propionaldehyde **5a** into WT containing the ThDP-enamine intermediate; B, Docking of propionaldehyde **5a** into 4M/HL/DE containing the ThDP-enamine intermediate; C, Docking of propionaldehyde **5a** into 4M/HL/DT containing the ThDP-enamine intermediate; D, Docking of 3-FBA **7** into WT containing the ThDP-enamine intermediate; E, Docking of 3-FBA **7** into 4M/HL/DE containing the ThDP-enamine intermediate; F, Docking of 3-FBA **7** into 4M/HL/DT containing the ThDP-enamine intermediate. Lightblue, mutational residues; Orange, active site residues which were only labelled in WT; Green, ThDP-enamine intermediate; Cyan, acceptor substrates including Propionaldehyde **5a** and 3-FBA **7**. The unit of the distance was angstrom. Residues surrounding the substrate-binding pocket are shown in surface representation.

3-FBA **7** is larger than propanal, and so the narrow substrate tunnel of 4M/HL/DE influenced the access of 3-FBA **7** to the enamine intermediate as evidenced by the distance of 6.1 Å, which was 1 Å and 0.9 Å longer than those for 3-FBA 7 docked within WT and 4M/HL/DT respectively (Table 2). This is consistent with the observation that 4M/HL/DE did not show any activity towards pyruvate **2d** and 3-FBA **7**. By contrast, the D469T mutation expanded the size of the substrate tunnel towards the intermediate, which allowed the occurrence of nucleophilic attack of the intermediate on the aldehyde group of 3-FBA **7**. This was similarly observed previously when we docked the 3-FBA **7** into D469T containing the (α, β-dihydroxyethyl)-ThDP intermediate [17]. 4M/HL/DT also showed a binding affinity of −5.5 kcal mol^-1^, 0.1 kcal mol^-1^ lower than that of WT. A possible explanation is that replacing the negatively charged residue aspartic acid with the more hydrophobic and uncharged threonine provides more favourable interactions with the non-polar acceptor substrates.

Consistent with the activity test, the docking experiments showed a picture of cooperation between different residues to improve the catalytic efficiency towards different substrates. Specifically, combination between H100L, responsible for the donor acceptance and D469E, responsible for propionaldehyde **2** acceptance lead to the mutation showing enhanced activity towards pyruvate and propanal, while the combination of H100L and D469T responsible for 3-FBA acceptance generate the mutation that improves the activity towards pyruvate and 3-FBA.

## Conclusion

In this study, we attempted to engineer the transketolase from *E. coli* to accept both the unnatural acceptor substrates propanal **5a**, pentanal **5b**, hexanal **5c** or 3-FBA **7** and a novel donor substrate, sodium pyruvate **2d** and its analogue keto-butyric acid **2e**, thus decreasing the polarity of both substrates. Twenty single-mutant variants were firstly designed based on structural alignment with PDC, and previous knowledge of variants in *E. coli* and *G. stearothermophilus.* The catalytic efficiency was then measured for these single-mutant variants experimentally. Beneficial mutations were then recombined using a previously obtained stable variant as the template, to construct a small but smart library. Library screening was carried out towards two reactions separately: 1) sodium pyruvate **2d** and propionaldehyde **5a**; 2) sodium pyruvate **2d** and 3-FBA **7**. The best variant 4M/H100L/D469E/R520Q showed a 9.2-fold improvement in the catalytic yield towards sodium pyruvate **2d** and propionaldehyde **5a** compared to WT. Pentanal and hexanal were also used as acceptors in the reactions with pyruvate **2d** or keto-butyric acid **2e** to determine stereoselectivities, which were found to be higher than 98% *ee* for the *S* configuration. Also, three unprecedented variants were identified to be active towards the reaction between sodium pyruvate **2d** and 3-FBA **7**. The best variant 4M/H100L/D469T was able to convert 47% of substrate into product within 24 h, whereas no conversion was observed for WT. Docking was applied to illustrate the cooperation between mutations responsible for donor acceptance and the mutation responsible for acceptor acceptance to promote the activity towards both the acceptor and donor, and this provides a guide for the future work about simultaneously engineering two substrate specificity. The hits obtained in this study have the potential to be used for developing catalytic pathways to a diverse range of products such as the phenylacetylcarbinols, an important pharmaceutical intermediate in the synthesis of drugs including spisulosine and phenylpropanolamine. This study also suggested that, in order to engineer enzymes for simultaneously accepting two novel substrates, a useful strategy is to identify the variants improving the single substrate acceptance first and then combine them together.

## Materials and methods

All chemicals were obtained from Sigma-Aldrich, UK unless mentioned otherwise.

### Preparation of lysates for enzyme reaction

The cell pellets from 12 mL cell culture were suspended using 1.2 mL cofactor solution containing 2.4 mM thiamine diphosphate (ThDP), 9 mM MgCl_2_, 50 mM Tris-HCl pH 7.0. The suspended cells were then transferred to a 2 mL eppendorf tube for cell lysis by sonication. The sonicated cells were centrifuged at 13000 rpm for 15 min to collect the supernatant as the cell lysates. The protein expression level was assessed by 12% sodium dodecyl sulfate-polyacrylamide gel electrophoresis (SDS-PAGE). Target bands were quantified using the GelDoc-It Imaging System (Cambridge, UK). Total protein concentrations in cell lysates were measured using the Bradford method [42]. Cell lysates were used at 2 mg/mL in enzyme reactions.

### Enzyme reactions

Enzyme reactions were carried out with lysates or purified proteins. Five acceptor substrates including glycolaldehyde **1**, propanal **5a**, pentanal **5b**, hexanal **5c** and 3-FBA **7** were used for reaction with the donor substrates sodium pyruvate **2d** or ketobutyric acid **2e**. Reactions between glycolaldehyde **1** and sodium pyruvate **2d** were initiated by adding 50 μL or 100 μL substrate solution containing 150 mM glycolaldehyde, 150 mM sodium pyruvate, 50 mM Tris-HCl, pH 7.0, into 100 μL purified enzyme with the concentration of 0.2 mg/mL or 200 μL enzyme lysates with the whole protein concentration of 2 mg/mL, to give a final substrate concentrations of 50 mM. Triplicate vials containing reaction solutions were incubated at 30 °C for 24 h. Samples were analysed by HPLC (Dionex, CA, USA) as previously [13] to determine the concentration of 3,4-dihydroxy-2-butanone **3** and erythrulose **4** against a standard curve.

Reactions between propionaldehyde **5a** and sodium pyruvate **2d** were initiated by adding 50 μL or 100 μL substrate solution containing 150 mM propanal, 150 mM sodium pyruvate **2d**, 50 mM Tris-HCl, pH 7.0, into 100 μL purified enzyme with the concentration of 0.2 mg/mL or 200 μL enzyme lysates with the whole protein concentration of 2 mg/mL, to give a final substrate concentrations of 50 mM. Triplicate vials containing reaction solutions were incubated at 30 °C for 24 h. A colorimetric assay detected the generation of the reaction product, 3-hydroxy-2-pentanone **6ad** using WST-1 instead of 2,3,5-triphenyltetrazolium chloride reported previously [43]. Compared to HPA, pyruvate does not reduce the tetrazolium red, also removing the need to add carbonate resin for extracting excess HPA as previously [43]. In a 96-well microplate, 50 μL reaction mixture was diluted further with 50 μL 50 mM Tris-HCl pH 7.0, then 20 μL WST-1 (0.2%) solution and finally 10 μL 3 M NaOH with good mixing. After incubation of 15 min at 25 °C, the microplate was scanned in a plate reader (BMG Fluostar Optima, Offenburg, Germany) for absorbance at 450 nm.

Reactions between aromatic aldehyde **7** and sodium pyruvate **2d** were initiated as above by adding solution containing 150 mM 3-FBA **7**, 150 mM sodium pyruvate **2d**, 50 mM Tris-HCl pH 7.0 into purified enzyme solution, to give final concentration of 50 mM 3-FBA **7** and 50 mM sodium pyruvate **2d**. After reaction at 30 °C for 24 h, 20 μL reaction mixture were transferred to 380 μL 0.1% TFA to quench the reaction. Samples were then analysed by HPLC with an ACE5 C18 reverse phase column (150 × 4.6 mm), UV detection at 210 nm and 250 nm, and a mobile phase at a flow rate of 1.0 ml/min starting with 85% of A (0.1% TFA) and 15% of B (acetonitrile). After 0.5 min, mobile phase B was linearly increased to 72% over 9 minutes, followed by a 2-minute re-equilibration at 85% of 0.1% TFA and 15% acetonitrile.

The reactions between acceptor substrates pentanal **5b** or hexanal **5c** and the donor substrates sodium pyruvate **2d** or keto-butyric acid **2e** were also carried out for isolating the products and determining the stereoselectivities. The detailed procedures of TK reactions, isolating the products and measuring stereoselectivities were provided in the supporting information (see SI).

### Site-directed mutagenesis and library construction

Site-directed mutagenesis was carried out with the QuikChange XL Site-Directed Mutagenesis Kit (Agilent Technologies, US), according to the manufacturer’s instructions, on the TK gene in plasmid pQR791 [44], and transformed into XL10-Gold ultracompetent cells contained in the kit. Mutagenic primers were designed using the web-based QuikChange Primer Design Program (www.agilent.com/genomics/qcpd).

A small library was constructed, based on the 8 single mutants H100F, H100L, H100T, D469E, D469T, D473N, D473S, R520Q, and combining them as described below to give 71 variants in total. Three rounds of mutagenesis were applied to construct the library. In the first round, site-directed mutagenesis was used to construct each of the following 12 variants, divided into three groups: group A were H100F, H100L, H100T, group B were D469E, D469T, D473N, D473S, D469E/D473N, D469E/D473S, D469T/D473N, D469T/D473S and group C was R520Q. In the second round, pairwise recombination of the groups as AB, BC and AC, was achieved by mixing templates from one group and primers from the other, to give 35 further variants. For instance, to combine group A and B mutations, the mixed PCR products of group A from the first round were used as the template, and the mixed mutagenic primers of group B were used as the primers for the Quikchange protocol. In the third round of PCR, the 24 possible new recombinants of groups ABC were constructed. PCR products from each round were mixed and transformed into ultracompetent cells. The library was then screened and the hits were sequenced.

### Library screening

Colonies from agar plates were picked and transferred to 2 mL 96 deep-well square plates (DWP) filled with 900 μL LB Amp^+^ medium. Three wells were inoculated with blank cells without the TK gene as the negative control. DWP plates were sealed with breathable sterile film (VMR International, US) and incubated at 37 °C, 400 rpm for 18 h in a humidified shaker. The final OD_600_ was measured from 50 μL cell culture transferred into a 96-well microplate. The remaining cell culture was centrifuged at 4000 rpm for 30 minutes to collect cell pellets, frozen at −80 °C and thawed at room temperature for freeze-thaw lysis of cells. The cell pellets were also suspended in 200 μL lysis buffer containing 9 mM MgCl_2_, 2.4 mM ThDP, 1×bugbuster and 1% lysonase (Merck, Hertfordshire, UK) in 50 mM Tris-HCl, pH 7.0 and then incubated at 25 °C for 30 minutes. Lysates were then transferred to a normal 96-well plate and centrifuged at 4000 rpm for 30 min, and 100 μL supernatant was used in enzyme reactions in glass vials, initiated by adding 50 μL aliphatic substrate solution containing 150 mM sodium pyruvate **2d**, 150 mM propionaldehyde **5a**, 50 mM Tris-HCl, pH 7.0, or 50 μL aromatic substrate solution containing 150 mM sodium pyruvate **2d**, 150 mM 3-FBA **7**, 50 mM Tris-HCl, pH 7.0. Reactions were incubated at 30 °C for 24 h before detection. For reactions with sodium pyruvate **2d** and propionaldehyde 5a, the colorimetric assay described above was used to detect the product. For reactions with sodium pyruvate **2d** and 3-FBA **7**, 20 μL samples were transferred to 380 μL 0.1% TFA to quench the reaction, centrifuged at 13000 rpm for 15 min, then 200 μL analysed by HPLC as above.

### Semi-preparative HPLC

The target peak detected by analytical HPLC was collected using semi-preparative HPLC on an Agilent ZORBAX 300SB-C18 (250 mm × 9.4 mm, 5 μm) reverse-phase column at 25 °C, with 2 mL/min flow rate and detection at 250 nm. The two column mobile phases were water with 0.1% TFA (A) and acetonitrile with 0.1% TFA (B), the column was pre-equilibrated with 85% A, 15% B. The reaction mixture was diluted 5-fold using 0.1 % TFA and then 200 μL loaded onto the column. A gradient was then applied as 85% A, 15% B at 0-5 min, then linearly ramped to 28% A, 72% B over 15 mins, and then linearly back to 85% A, 15% B over 5 mins.

### Gas chromatography

After the enzyme reaction, 500 μL chloroform was added and the mixture agitated, then centrifuged at 4000 rpm for 3 min. The organic phase was then pipetted out carefully and analysed by GC/MS with an ISQ series single Quadrupole GC-MS system (ThermoFisher Scientific, MA, US), equipped with a Rxi-5Sil MS column (30 m × 0.2 mm I.D.), and the helium carrier gas at a flow rate of 1 mL/min. The column temperature was kept at 40 °C for 3 min, then raised to 150 °C at a rate of 2 °C/min, then to 200 °C at a rate of 4 °C/min. The ion source temperature was 230 °C and the range of molecular weight scanned was 20-350 m/z. GC analyses were also carried out with an Agilent 7820A GC-system and a chiral stationary phase: Supelco RT BetaDEX225 column (30 m x 0.25 mm, 0.25 μm) and enantiomeric excess measurements were determined with the following method: Starting temperature 100 °C and an increase of 5 °C per minute until 135 °C, then 40 °C per minute to 210 °C.

### Liquid chromatography-mass spectrometry

MS analysis was performed in the UCL Chemistry Mass Spectrometry Facility. The molecular masses of the compounds were measured on an Acquity UPLC System (Waters, MA, US) with an SQ mass spectrometer (Waters, MA, US). The HPLC was performed with a Thermo Scientific Hypersil Gold^TM^ C4 reverse phase column (50 × 2.1 mm) with 0.2 mL/min flow rate. The mobile phases were 0.1% (v/v) formic acid (A) and 95% acetonitrile-5% water (B), equilibrated at 95% A, 5% B, and then a gradient applied after sample injection to 5% A, 95% B over 4 mins, then back to 95% A, 5% B over 0.5 mins, and held for a further 0.5 mins. Compounds were ionized using electrospray ionization (ESI) and detected in positive mode. The operating condition of ESI interface were set to a capillary temperature of 300 °C, capillary voltage 9 V and spray voltage 4 kV.

### Nuclear magnetic resonance spectroscopy

^1^H NMR spectra were recorded on Bruker Avance 400, 500, 600 and 700 MHz spectrometers at 25 °C, using the residual protic solvent stated as the internal standard. Chemical shifts are quoted in ppm to the nearest 0.01 ppm using the following abbreviations: s (singlet), d (doublet), t (triplet), q (quartet), qn (quintet), sext (sextet), dd (doublet of doublets), dt (doublet of triplets), m (multiplet) defined as all multi-peak signals where overlap or complex coupling of signals make definitive descriptions of peaks difficult. The coupling constants are defined as J and quoted in Hz. ^13^C NMR spectra were recorded at 100, 125, 150 or 175 MHz on Bruker Avance 400, 500, 600 and 700 MHz spectrometers at 25 °C using the stated solvent as standard. Chemical shifts are reported to the nearest 0.1 ppm. Melting points were measured with a Gallenkamp apparatus and were uncorrected.

### Structure alignment

The TK structure 2R8O.pdb and pyruvate decarboxylase structure 2VK1.pdb were aligned using the server, TopMatch-Web [45, 46]. The aligned structure from output was visualized using Pymol (Schrödinger, USA). Sequence identity was calculated online by the web server Sequence Identity And Similarity (SIAS, http://imed.med.ucm.es/Tools/sias.html) with the input of 2R8O.pdb and 2VK1.pdb.

### Computational Molecular Docking

The structures of the variants H192P/A282P/I365L/G506A/H100L/D469E (4M/HL/DE) and H192P/A282P/I365L/G506A/H100L/D469T(4M/HL/DT) were obtained using SWISS-MODEL and 1QGD.pdb as the template [47]. ThDP-enamine, propionaldehyde 5a and 3-FBA 7 structures were drawn in Chem3D Ultra v.13.0 and energy minimised by MM2 calculations.

Autodock Vina was used to sequentially dock the ThDP-enamine and propionaldehyde 5a or ThDP-enamine and 3-FBA 7 into wild-type and mutant TKs with the grid centred at −11.25, 25.858, 40.198 and the exhaustiveness of 24 [48, 49]. The grid sizes of 30 Å × 30 Å × 30 Å, 10 Å × 10 Å × 10 Å and 28 Å × 28 Å × 28 Å were used for the docking of ThDP-enamine, propionaldehyde 5a and 3-FBA 7 respectively. Five replicate dockings were carried out and the conformations with lowest energy were selected for analysis.

## Author Contributions

P.A.D. designed the research. H.Y., R.I.H.L., D.S., D.M.S., S.H., and A.C.K performed the experiments. H.Y., P.A.D. and H.C.H. interpreted the data. P.A.D., H.C.H. and J.M.W. supervised the studies. H.Y., P.A.D. and H.C.H. wrote the paper. All authors reviewed, revised, and approved the manuscript for publication.

## Acknowledgement

This work was supported in part by a Chinese Scholarship Council stipend (to H.Y.) and Engineering and Physical Sciences Research Council Grants EP/N025105/1 and EP/P006485/1. We gratefully acknowledge the Department of Chemistry at University College London for funding S.H. and D.S., the Biotechnology and Biosciences Research Council (BBSRC) (BB/N01877X/1) for funding D.M.S. and CONACYT is thanked for a PhD studentship to A.C.K.

## Schemes

**Scheme 1.**
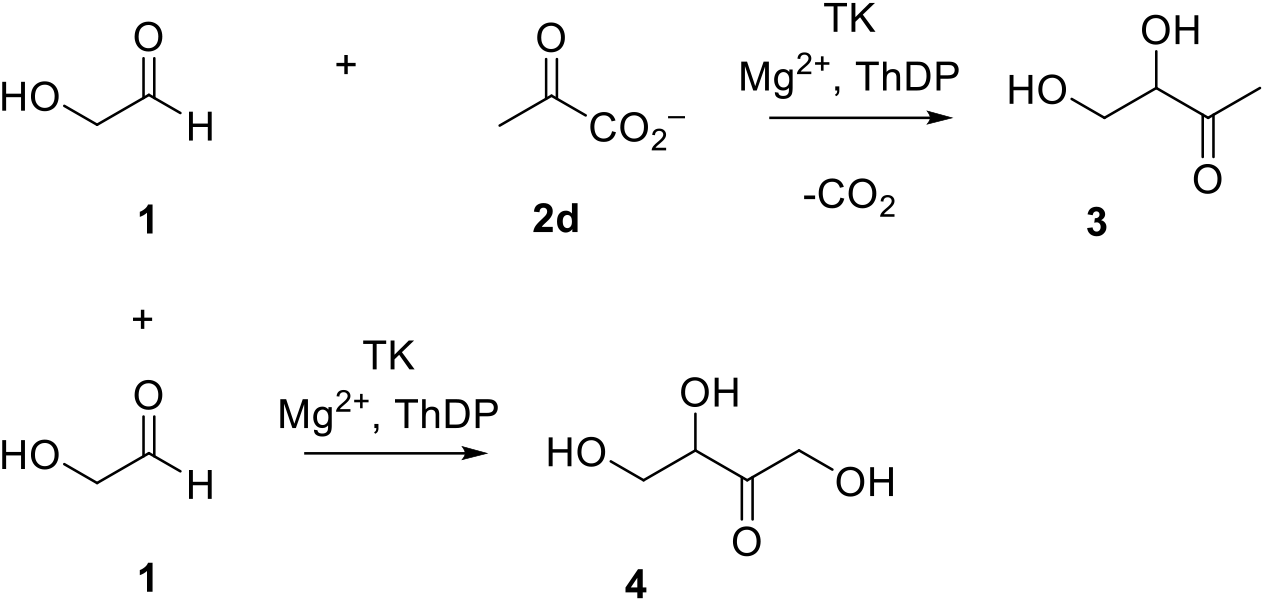
TK mediated the reaction between glycolaldehyde 1 and pyruvate 2d. Target product 3,4-dihydroxy-2-butanone (DHB) **3** and side product erythrulose **4** were generated.

**Scheme 2.**
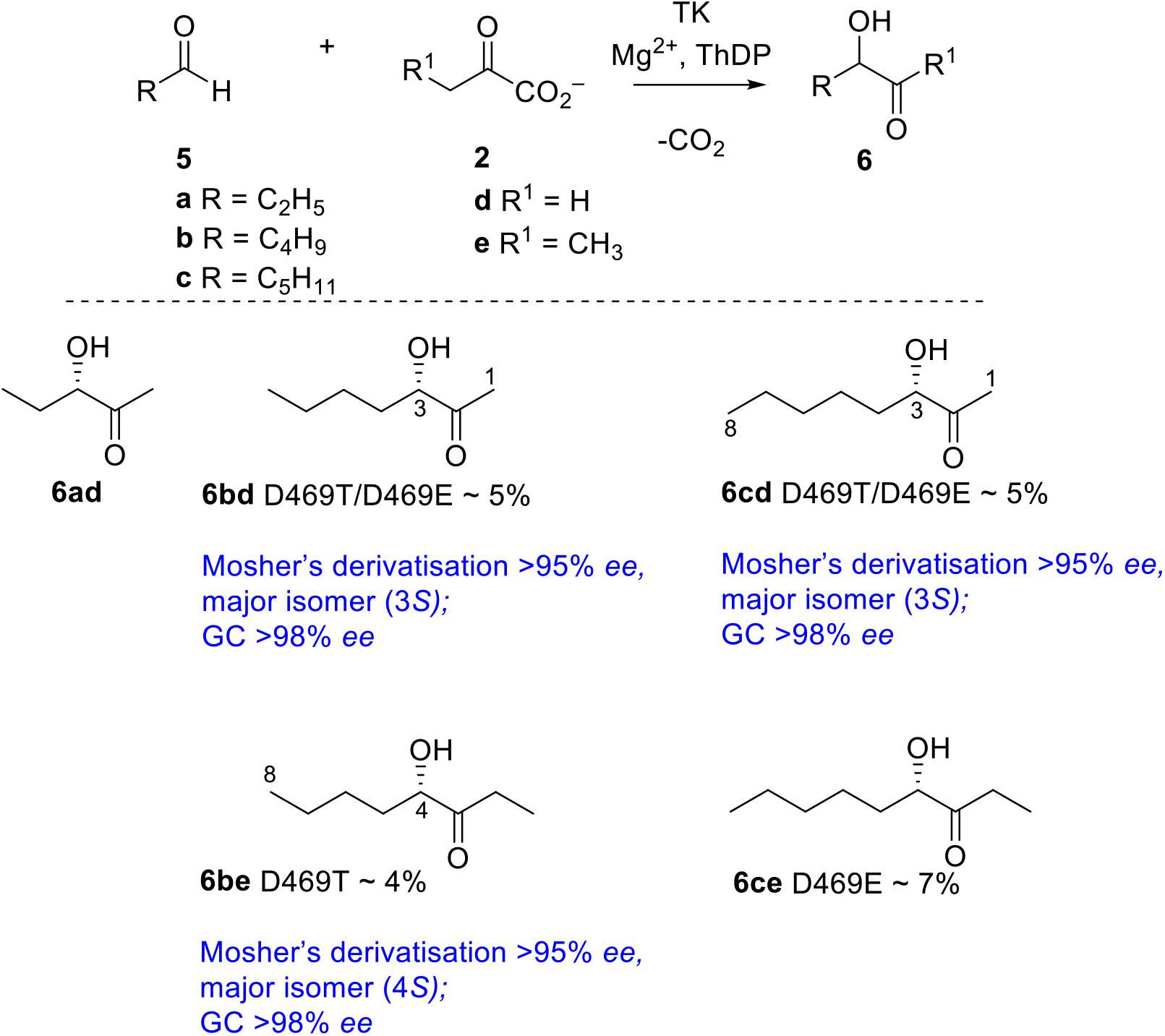
TK mediated the reactions between aliphatic acceptor aldehydes and donor substrates including pyruvate 2d and keto-butyric acid 2e.

**Scheme 3.**
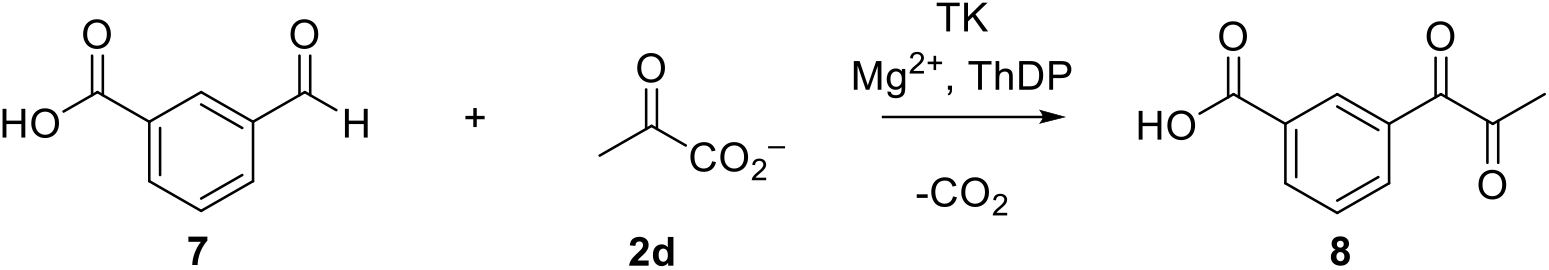
TK mediated the reaction between aromatic aldehyde 3-formylbenzoic acid 7 and pyruvate 2d.

### Supporting information

Additional supporting information may be found online in the supporting information section at the end of the article.

**Table S1** TK (2R8O.pdb) and PDC (2VK1.pdb) structure alignment parameters

**Table S2** Design of single variants

**Table S3** Characteristics of the variant 4M/HL/DT

**Fig. S1** Structure alignment between transketolase (2R8O.pdb) and pyruvate decarboxylase (2VK1.pdb)

**Fig. S2** Densitometry to measure TK variants expression in the lysates with the protein concentration of 2 mg/mL

**Fig. S3** Correlation between colorimetric assay and the GC-MS measurement for the reactions yield

**Fig. S4** Activity correlation between reactions with sodium pyruvate as donor and with glycolaldehyde or propionaldehyde as acceptor

**Fig. S5** LC-MS analysis of the reaction product between 3-FBA and sodium pyruvate

**Fig. S6** Pose of ThDP-enamine intermediate in the binding pockets of wild-type and mutant TKs

**Fig. S7** Structural alignment of TK wild type and two variants 4M/HL/DE, 4M/HL/DT

**Appendix S1** General experimental for synthesis and analysis

**Appendix S2** General procedures for TK reactions

**Appendix S3** Stereoselectivity **measurements**

**Appendix S4 NMR spectra**

