## supplementary information for "Simultaneous optimization of donor and acceptor substrate specificity for transketolase by a small but smart library"

**Table S1 TK (2R8O.pdb) and PDC (2VK1.pdb) structure alignment parameters**

| LEN[a] | QC%[b] | TC%[c] | SCORE[d] | RMS[e] | SI%[f] |
|---|---|---|---|---|---|
| 559 | 42.1% | 24.9% | 378.1 | 2.66 | 12.3 |

[a] Number of residue pairs that are structurally equivalent.
[b] TK structure cover based on similarity score, expressed in percent. (= 100 x SCORE/$Q_n$, where $Q_n$ is the length of the TK sequence).
[C] PDC structure cover based on similarity score, expressed in percent (= 100 x SCORE/$T_n$, where $T_n$ is the length of the PDC sequence).
[d] Measure of structural similarity based on Gaussian functions.
[e] Root-mean-square error of superposition in angstrom, calculated using all structurally equivalent C-alpha atoms.
[f] Sequence identity of TK and PDC in the equivalent regions, expressed in percent.

**Table S2 Design of single variants**

| Methods | Structure alignment (16) | Previous knowledge (4) |
|---|---|---|
| Mutants | H26I H26Q H66I H66T H100F H100L H100T H100Y L116I I189E I189N I189Q H261E H261I S385D D469H | H473N H473S D469E D469T |

**Table S3 Characteristics of the variant 4M/HL/DT**

| | Specific activity [a] (μmol $mg^{-1}min^{-1}$) | Conversion at 24 h [b] (%) | $K_m$ (mM) | $k_{cat}$ ($s^{-1}$) | $k_{cat}/K_m$[c] ($s^{-1}$ $M^{-1}$) |
|---|---|---|---|---|---|
| 4M/HL/DT | 0.048(0.005) | 46.8(2.0) | 90.4 (11) | 0.22(0.01) | 2.4 |

[a] Specific activity was tested towards 50 mM 3-FBA and 50 mM sodium pyruvate in 50 mM Tris-HCl solution at 30 °C with the enzyme concentration of 1.33 mg/mL.

[b] Yield was measured based on the consumption of substrates

[c] Kinetic parameters were measured towards 50 mM 3-FBA and sodium pyruvate with different concentration from 5 mM to 250 mM.

A

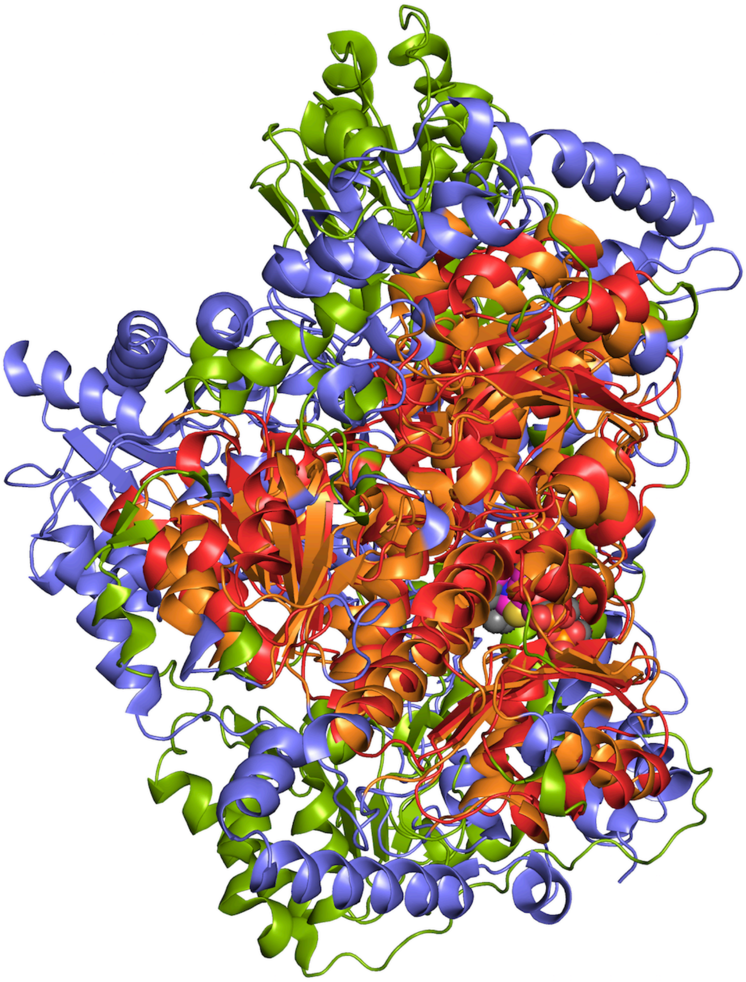

B

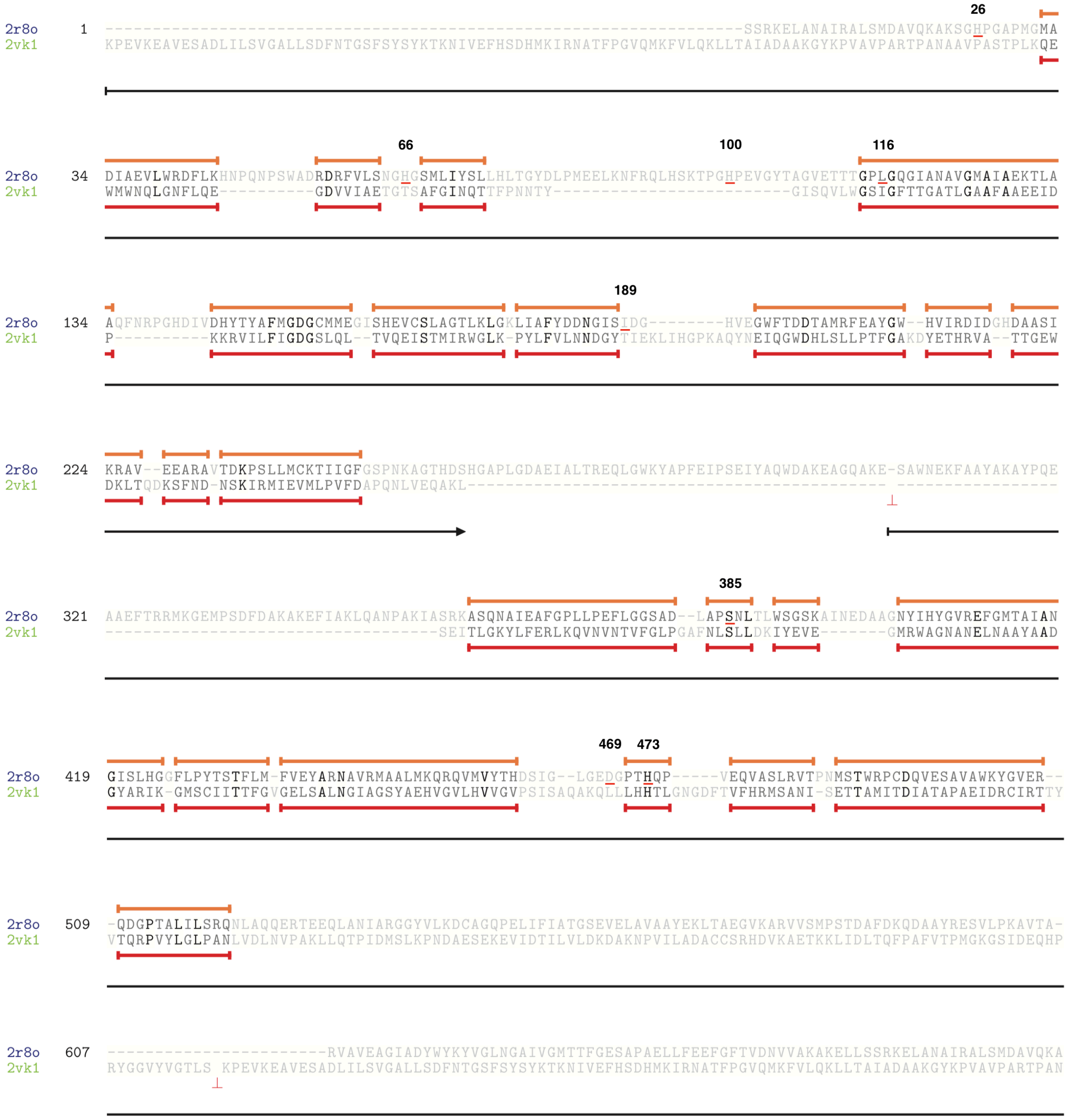

**Figure S1.** Structure alignment between transketolase (2R8O.pdb) and pyruvate decarboxylase (2VK1.pdb). A, structures of aligned TK and PDC. Blue, structure of transketolase; Green, structure of pyruvate decarboxylase. B, sequence alignment indicating the structurally equivalent residues. Pairs of residues that are structurally equivalent are coloured orange (TK) or red (PDC).

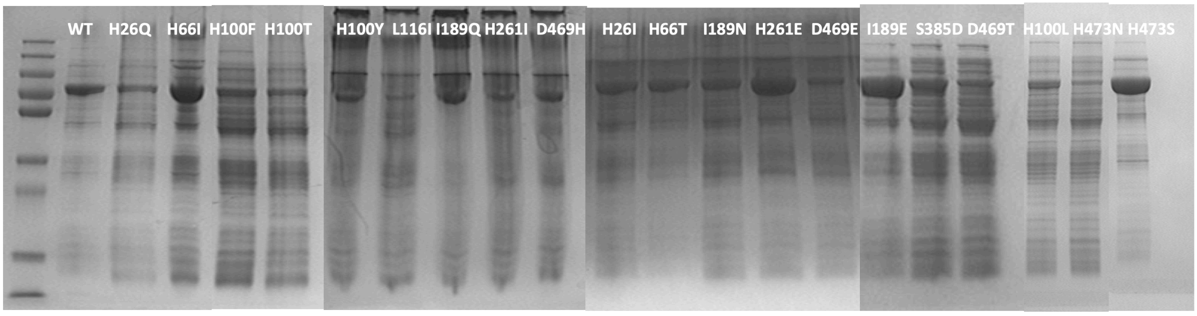


| | | | | | | | |
|---|---|---|---|---|---|---|---|
| **Variants** | WT | H26Q | H66I | H100F | H100T | H100Y | L116I |
| **Density** | 106.8 | 53.3 | 189.7 | 88.4 | 63.5 | 47.4 | 26.9 |
| **Variants** | I189Q | H261I | D469H | H26I | H66T | I189N | H261E |
| **Density** | 85.7 | 43.3 | 73.7 | 145.2 | 126.4 | 135.9 | 283.3 |
| **Variants** | D469E | I1898E | S385D | D469T | H100L | H473N | H473S |
| **Density** | 74.0 | 242.5 | 153.4 | 53.5 | 94.5 | 35.8 | 227.8 |

**Figure S2.** Densitometry to measure TK variants expression in the lysates with the protein concentration of 2 mg/mL. Density of target band for wild type and each variant was listed under the SDS-PAGE. The protein molecular weight marker (Up-250KD, 130KD, 95KD, 72KD, 55KD, 36KD, 28KD, 17KD, 10KD-Bottom); The size of the TK was expected to be 73 KDa on SDS/PAGE.

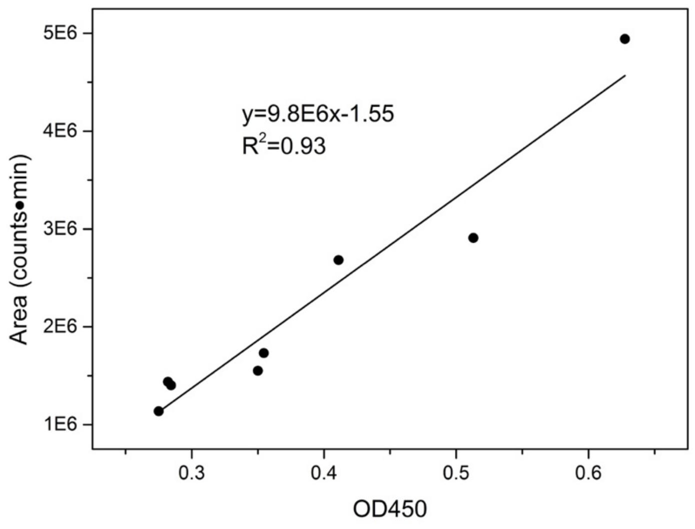


**Figure S3.** Correlation between colorimetric assay and the GC-MS measurement for the reactions yield.

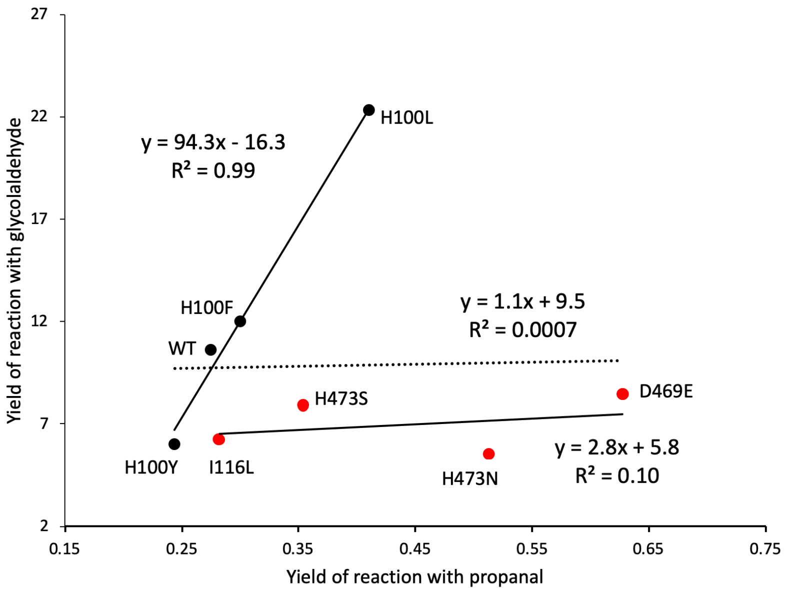


**Figure S4.** Activity correlation between reactions with sodium pyruvate as donor and with glycolaldehyde or propionaldehyde as acceptor. The yield was the concentration of DHB for the reaction between glycolaldehyde and pyruvate, and OD450 for the reaction between propanal and pyruvate.

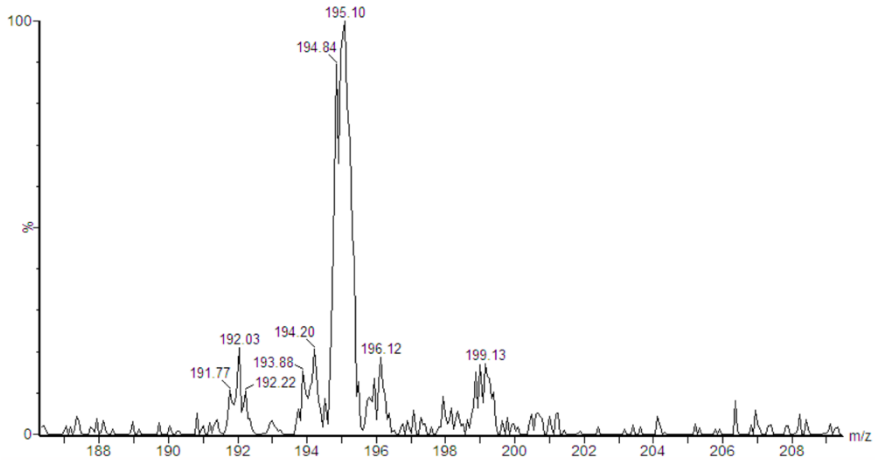


**Figure S5.** LC-MS analysis of the reaction product between 3-FBA **7** and sodium pyruvate **2d.**

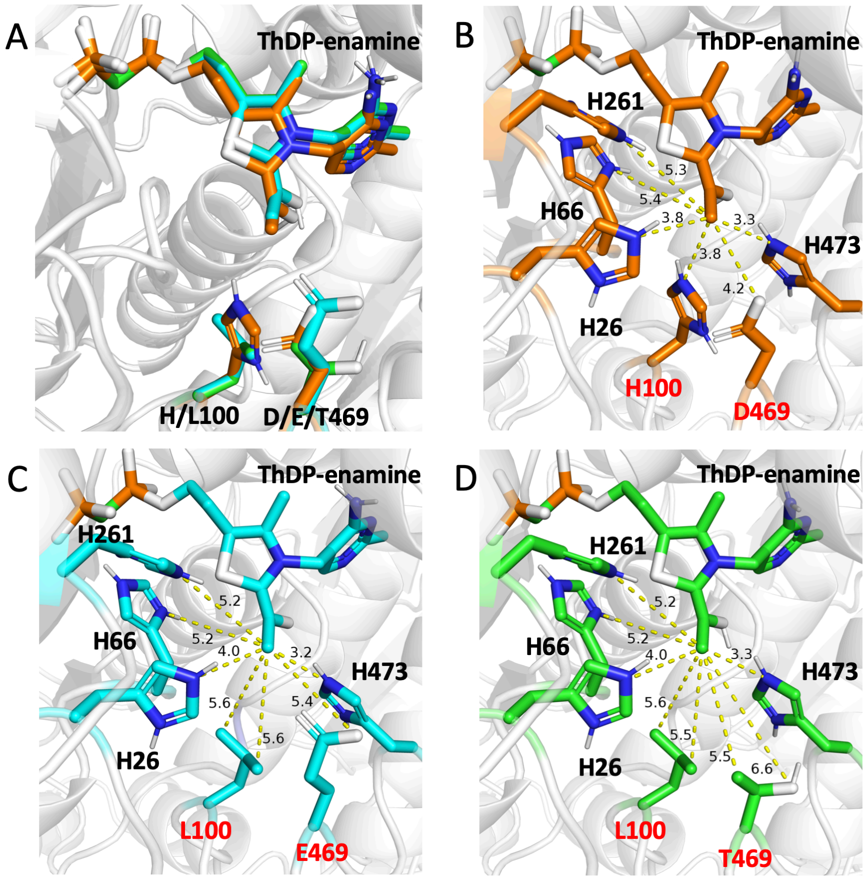


**Figure S6**. Pose of ThDP-enamine intermediate in the binding pockets of wild-type and mutant TKs. A, pose comparisons of ThDP-enamine in the WT and variants; B, ThDP-enamine docked into WT; C, ThDP-enamine docked into 4M/HL/DE; D, ThDP-enamine docked into 4M/HL/DT. Orange, WT; Cyan, 4M/HL/DE; Green, 4M/HL/DT

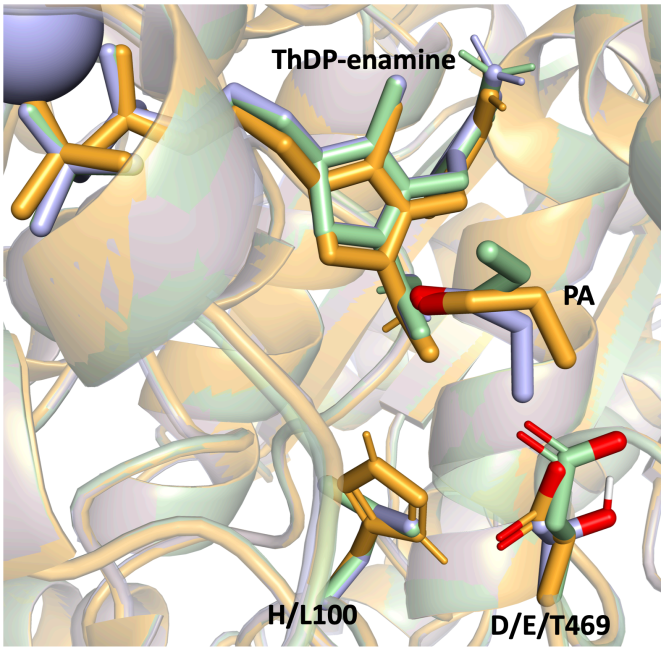


**Figure S7**. Structural alignment of TK wild type and two variants 4M/HL/DE, 4M/HL/DT. Light orange, WT; light blue, 4M/HL/DT; light green, 4M/HL/DE.

**General experimental for synthesis and analysis**

All chemicals were obtained from commercial suppliers and used as received. Small- scale reactions were heated using a BIOER Mixing block MB-102, and the scale-up reactions were heated using Heidolph MR Hei-Tec type heating mantle. Thin layer chromatography was carried out using Merck TLC Silica gel 60 $F_{254}$ plates and products were visualized using combinations of UV light (254 nm), potassium permanganate and phosphomolybdic acid staining solutions. Column chromatography was carried out using silica gel (particle size 40–60 µm). Infrared (IR) spectra were recorded using a Perkin Elmer Spectrum 100 FT-IR Spectrometer or a Bruker Alpha Platinum-ATR, operating in ATR mode. Mass spectra were obtained using a Waters Acquity UPLC SQD (using a linear gradient of 5-95% of acetonitrile over 5 min, with a C8 column, and flow rate of 0.6 mL/min) and Waters LCT Premier XE ESI Q-TOF mass spectrometer in the Department of Chemistry, UCL.

**General procedures for TK reactions between acceptor substrates pentanal or hexanal and the donor substrates sodium pyruvate or ketobutyric acid**

A stock "10x" cofactor solution of $MgCl_2$ (0.39 g) and ThDP (0.22 g) in water (10 mL) was adjusted to pH 7 and stored at 4 °C. 1 mL of the 10x cofactor solution was mixed with TK cell-free lysate (2 mL or 4 mL, as indicated) and water was added to bring the volume to 10 mL. The lysate mixture was incubated for 20 minutes. In a separate flask, a mixture of 100 mM of the donor and acceptor was prepared and adjusted to pH 7, then added to the enzyme mixture, to give a final substrate concentration of 50 mM of each. Reactions were then stirred at room temperature for 24 h or 48 h, as indicated, either in buffer at pH 7, or in an autotitrator programmed to add 1 M HCl when the pH increased above 7.

**Stereoselectivity measurements**

***Rac*-3-Hydroxyheptan-2-one**

**Step 1**

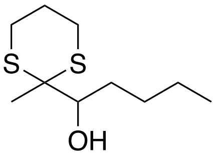


To solution of 2-methyl-1,3-dithiane[1] (479 µL, 4.32 mmol) in dry THF (16 mL) at -78 °C, *n*-butyllithium (2.5 M in hexane; 2.0 mL, 5.0 mmol) was added dropwise. The reaction was stirred at -78 °C for 30 min, at 0 °C for a further 30 min and then cooled to -78 °C. Pentanal (638 µL, 6.00 mmol) was added dropwise and the reaction mixture stirred at room temperature for 18 h. Water (16 mL) was added, followed by dichloromethane (16 mL) and the product extracted with dichloromethane (3 × 50 mL). The volume of the combined organic fraction was reduced under vacuum to approximately 50 mL, then washed with aqueous potassium hydroxide solution (7%, 20 mL) and sat. sodium chloride solution (20 mL), dried ($MgSO_4$), and the solvent was removed *in vacuo*. The crude product was purified by flash silica column chromatography (EtOAc/pet. ether 40-60, 1:9) to give 1-(2-methyl-1,3-dithian-2-yl)pentan-1-ol[2] as a pale yellow oil (379 mg, 40%). $R_f$ 0.30 (EtOAc/pet. ether 40-60, 1:4); $\nu_{max}$ (film, $cm^{-1}$) 3480, 2952; $^1$H NMR (600 MHz; $CDCl_3$) $\delta$ 0.91 (3H, t, *J* = 7.1 Hz, $CH_2CH_3$), 1.18-1.49 (7H, m, $CH_3$ and $(CH_2)_2$), 1.54-1.63 (1H, m, CHOHC*H*H), 1.80-1.86 (1H, m, $SCH_2C$*H*H), 1.90-1.96 (1H, m, CHOHCH*H*), 2.03-2.09 (1H, m, $SCH_2CH$*H*), 2.55-2.63 (2H, m, 2 × SC*H*H), 2.94-3.00 (2H, m, 2 × SCH*H*), 3.50-3.60 (1H, m, C*H*OH), 3.91 (1H, br d, *J* = 9.1 Hz, OH); $^{13}$C NMR (151 MHz; $CDCl_3$) $\delta$ 14.2, 21.9, 22.7, 24.4, 27.8, 29.5, 30.0, 32.2, 54.1, 71.3; *m/z* HRMS (ESI+) found $[MH]^+$ 221.1032, $C_{10}H_{21}OS_2$ requires 221.1034.

**Step 2**

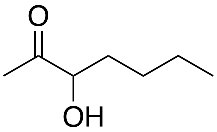


To a solution of 1-(2-methyl-1,3-dithian-2-yl)pentan-1-ol[2] (331 mg, 1.50 mmol) in water (40 mL), acetonitrile (8 mL) and THF (5 mL), iodomethane (2.99 mL, 48.2 mmol) and calcium carbonate (1.80 g, 18.0 mmol) were added and the reaction mixture heated at reflux for 4 h. The reaction was cooled to room temperature and the product extracted with diethyl ether (3 × 50 mL), dried ($MgSO_4$), filtered, and the solvent removed under vacuum. The crude product was purified by flash silica column chromatography (diethyl ether/pentane, 1:9) to give 3-hydroxyheptan-2-one[3] as a pale yellow oil (145 mg, 74%). $R_f$ 0.22 (EtOAc/pet. ether 40-60, 1:9); $\nu_{max}$ (film, $cm^{-1}$) 3420, 2928, 1709; $^1$H NMR (600 MHz; $(CD_3)_2CO$) $\delta$ 0.92 (3H, t, *J* = 7.2 Hz, $CH_2CH_3$), 1.28-1.43 (4H, m, $(CH_2)_2CH_3$), 1.50-1.58 (1H, m, CHOHCH*H*), 1.72-1.80 (1H, m, CHOHC*H*H), 2.17 (3H, s, $COCH_3$), 4.06-4.18 (2H, m, OH and C*H*OH); $^{13}$C NMR (151 MHz;

$(CD_3)_2CO$) δ 13.7, 22.7, 24.9, 27.4, 33.5, 77.2, 210.9; *m/z* HRMS (ESI+) found $[M+NH_4]^+$ 148.1332, $C_7H_{18}NO_2$ requires 148.1338.

**(3*S*)-Hydroxyheptan-2-one**

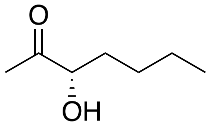


The co-factor solution (total volume 10 mL, pH 7) was prepared as described above in the general procedure for TK reactions. D469T or D469E TK cell-free lysate (2 mL) was added and the mixture was incubated for 20 min. Sodium pyruvate (0.110 g, 1.00 mmol) was dissolved in water (10 mL) with pentanal (0.106 mL, 1.00 mmol) and the pH adjusted to 7. This was then added to the enzyme suspension and the reaction was stirred in an autotitrator for 18 h while the reaction was followed by TLC analysis. The product was then extracted with $Et_2O$ (2 x 500 mL), and the organic layer was dried ($MgSO_4$), filtered and then dry-loaded onto silica before being purified by flash silica chromatography ($Et_2O$:pentane, 5:95) to give (3*S*)-hydroxyheptan-2-one as a pale green oil (0.006g, 5%). The compound is volatile so material was lost during the purification step. The reaction was also performed with D469E which gave the same isolated yield. NMR characterization data was consistent with that of the racemate; $^1$H NMR (600 MHz; ($CDCl_3$) δ 0.91 (3H, m, $CH_2CH_3$), 1.25-1.43 (4H, m, $(CH_2)_2CH_3$), 1.50-1.58 (1H, m, CHOHCH*H*), 1.80-1.85 (1H, m, CHOHC*H*H), 2.18 (3H, s, $COCH_3$), 4.16-4.22 (1H, m, C*H*OH); $^{13}$C NMR (151 MHz; ($CDCl_3$) δ 14.0, 22.6, 25.3, 27.0, 33.3, 77.0, 210.2. $[\alpha]^{20}_D$ = +33.3 (*c* 0.30, $Et_2O$), Lit. $[\alpha]^{15}_D$ = +12.5 (*c* 0.40, MeOH)[4].

The product stereoselectivity was established by GC analysis of the racemate which revealed 2 peaks with retention times (rt) of 5.5 min (*S*-isomer) and 5.8 min (*R*-isomer) example spectrum **A** below). The enzyme reaction gave a single peak at a rt 5.5 min only (S-isomer) (example spectrum **B** below) revealing an ee of >98%.

**A** GC trace of *rac*-3-Hydroxyheptan-2-one

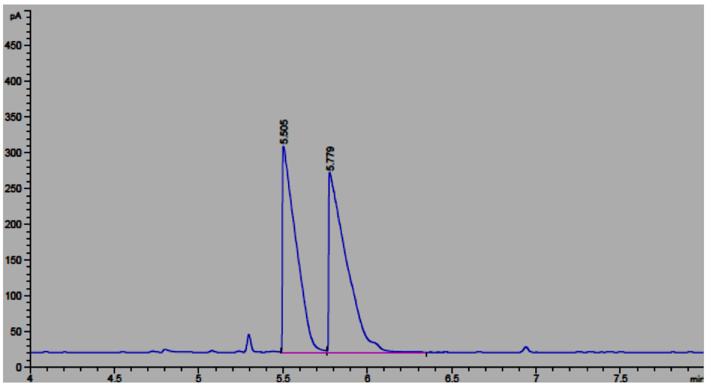

**B** GC trace of (3*S*)-hydroxyheptan-2-one from the reaction with D469T

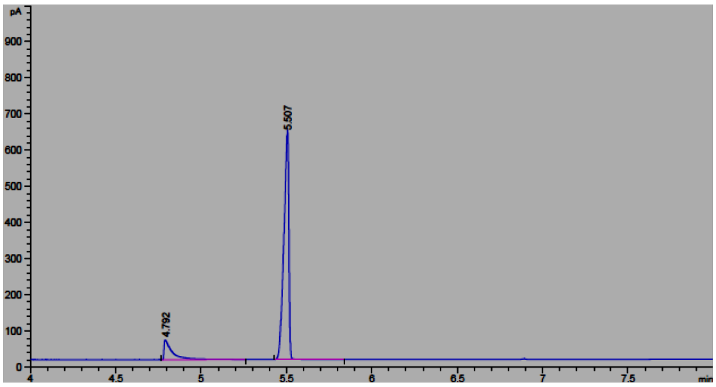


The product from the enzyme reaction was analysed further by derivatisation as the Mosher's ester.

**(2*S*,3*S*)-2-Oxoheptan-3-yl-3,3,3-trifluoro-2-methoxy-2-phenylpropanoate**

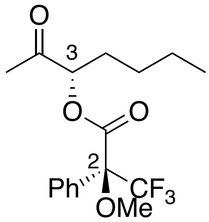


To a stirred solution of (3*S*)-hydroxyheptan-2-one (0.006 g, 0.05 mmol) in $CH_2Cl_2$ (5 mL) was added triethylamine (0.020 mL, 0.14 mmol) and (*R*)-MTPA-Cl (0.020 mL, 0.1 mmol) and the reaction stirred at room temperature for 18 h. The crude reaction mixture was then dry-loaded onto silica and purified by flash silica chromatography (EtOAc:hexane, 5:95) to give the titled compound as a colourless oil (1 mg, 6%). $R_f$ 0.5 (EtOAc/hexane, 1:9); $^1$H NMR (600 MHz; $CDCl_3$) $\delta$ 0.83 (3H, t, *J* = 7.1 Hz, $CH_2CH_3$), 1.20-1.28 (4H, m, $CH_2CH_2$), 1.75-1.84 (2H, m, $CH_2$CHO), 2.13 (0.04H, s, (2*R*,3*R*)-$COCH_3$), 2.21 (3H, s, (2*R*,3*S*)-$COCH_3$), 3.64 (3H, s, $OCH_3$), 5.17 (1H, dd, *J* = 4.5 and 4.1 Hz, C*H*OC=O), 7.43 (3H, m, Ar*H*), 7.64 (2H, m, Ar*H*); $^{13}$C NMR (151 MHz; $CDCl_3$) $\delta$ 13.8, 22.1, 26.6, 27.1, 29.8, 55.8, 80.5, 84.7 (q, $^2J_{CF}$ = 22.4 Hz), 127.6, 128.5, 129.8, 132.1, 166.6, 203.7; $^{19}$F NMR (282 MHz; $CDCl_3$) $\delta$ –72.1; *m/z* HRMS (CI+) found $[MH]^+$ 347.1477, $C_{17}H_{22}O_4F_3$ requires 347.1470.

***Rac*-3-Hydroxyoctan-2-one**

**Step 1**

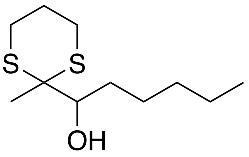

To solution of 2-methyl-1,3-dithiane[1] (479 µL, 4.32 mmol) in dry THF (16 mL) at -78 °C, *n*-butyllithium (2.5 M in hexane; 2.0 mL, 5.0 mmol) was added dropwise. The reaction was stirred at -78 °C for 30 min, at 0 °C for a further 30 min and then cooled to -78 °C. Hexanal (737 µL, 6.00 mmol) was added dropwise and the reaction mixture stirred at room temperature for 18 h. Water (16 mL) was added, followed by dichloromethane (16 mL) and the product extracted with dichloromethane (3 × 50 mL). The volume of the combined organic fraction was reduced under vacuum to approximately 50 mL, then washed with aqueous potassium hydroxide solution (7%, 20 mL) and sat. sodium chloride solution (20 mL), dried ($MgSO_4$), and the solvent was removed *in vacuo*. The crude product was purified by flash silica chromatography (EtOAc/pet. ether 40-60, 1:9) to give 1-(2-methyl-1,3-dithian-2-yl)hexan-1-ol as a pale yellow oil (365 mg, 39%). R*f* 0.30 (EtOAc/pet. ether 40-60, 1:4); $\nu_{max}$ (film, $cm^{-1}$) 3463, 2958; $^1H$ NMR (600 MHz; $CDCl_3$) $\delta$ 0.89 (3H, t, *J* = 7.1 Hz, $CH_2CH_3$), 1.18-1.48 (9H, m, $CH_3$ and $(CH_2)_3$), 1.56-1.65 (1H, m, CHOHC*H*H), 1.80-1.86 (1H, m, $SCH_2C$*H*H), 1.89-1.95 (1H, m, CHOHCH*H*), 2.03-2.09 (1H, m, $SCH_2CH$*H*), 2.55-2.63 (2H, m, 2 × SC*H*H), 2.94-3.00 (2H, m, 2 × SCH*H*), 3.51-3.61 (1H, m, C*H*OH), 3.91 (1H, br d, *J* = 9.6 Hz, OH); $^{13}C$ NMR (151 MHz; $CDCl_3$) $\delta$ 14.1, 21.9, 22.8, 24.4, 26.3, 27.8, 29.5, 30.1, 32.0, 54.0, 71.3; *m/z* HRMS (ESI+) found $[MH]^+$ 235.1190, $C_{11}H_{23}OS_2$ requires 235.1190.

**Step 2**

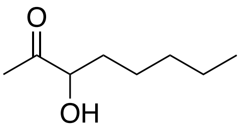

To a solution of 1-(2-methyl-1,3-dithian-2-yl)hexan-1-ol (352 mg, 1.50 mmol) in water (40 mL), acetonitrile (8 mL) and THF (5 mL), iodomethane (2.99 mL, 48.2 mmol) and calcium carbonate (1.80 g, 18.0 mmol) were added and the reaction mixture heated at reflux for 4 h. The reaction was cooled to room temperature and the product extracted with diethyl ether (3 × 50 mL), dried ($MgSO_4$), filtered, and the solvent removed under vacuum. The crude product was purified by flash silica column chromatography (diethyl ether/pentane, 1:9) to give 3-hydroxyoctan-2-one[5] as a pale yellow oil (154 mg, 71%). $R_f$ 0.22 (EtOAc/pet. ether 40-60, 1:9); $\nu_{max}$ (film, $cm^{-1}$) 3450, 2925, 1710, 1457; $^1H$ NMR (600 MHz; $(CD_3)_2CO$) $\delta$ 0.85-0.95 (3H, m, $CH_2C$*$H_3$*), 1.25-1.47 (6H, m, (C*$H_2$*)$_3CH_3$), 1.50-1.56 (1H, m, CHOHCH*H*), 1.72-1.78 (1H, m, CHOHC*H*H), 2.18 (3H, s, $COCH_3$), 4.06-4.14 (2H, m, OH and C*H*OH); $^{13}C$ NMR (151 Mz; $(CD_3)_2CO$) $\delta$ 13.8, 22.7 24.8, 25.0, 31.9, 33.8, 77.2, 210.9; *m/z* HRMS (ESI+) found $[M+NH_4]^+$ 162.1494, $C_8H_{20}NO_2$ requires 162.1494.

**(3*S*)-Hydroxyoctan-2-one**

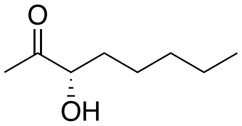

The co-factor solution (total volume 10 mL, pH 7) was prepared as described above in the general procedure for TK reactions. D469T or D469E TK free lysate (2 mL) was added and the mixture was incubated for 20 min. To sodium pyruvate (0.110 g, 1.00 mmol) in water (10 mL) and hexanal (0.122 mL, 1.00 mmol) the pH was adjusted to 7. This was then added to the enzyme suspension and the reaction was stirred in an autotitrator for 18 h while the reaction was followed by TLC analysis. The product was then extracted with $Et_2O$ (2 x 500 mL), and the organic layer was dried ($MgSO_4$), and then dry-loaded onto silica before being purified by flash silica chromatography ($Et_2O$:pentane, 5:95) to give (3*S*)-hydroxyoctan-2-one as a pale green oil (0.007g, 6%). The compound is volatile so some material was lost during the purification step. NMR characterization data was consistent with that of the racemate; $^1$H NMR (600 MHz; ($CDCl_3$) δ 0.89 (3H, m, $CH_2CH_3$), 1.25-1.47 (6H, m, $(CH_2)_3CH_3$), 1.50-1.56 (1H, m, CHOHCH*H*), 1.72-1.78 (1H, m, CHOHC*H*H), 2.20 (3H, s, $COCH_3$), 4.15-4.24 (1H, m, C*H*OH); $^{13}$C NMR (151 Mz; ($CDCl_3$) δ 14.1, 22.6 24.5, 25.3, 31.7, 33.6, 210.4. $[\alpha]^{20}_D$ = +8.5 (*c* 0.35, $Et_2O$), Lit. $[\alpha]^{20}_D$ = +57.3 (*c* 1.1, $CHCl_3$)[6].

The product stereoselectivity was established by GC analysis; rt 5.5 min (*S*-isomer) and 5.8 min (*R*-isomer). The enzyme reaction gave a single peak at a rt 5.5 min only (*S*-isomer) (example spectrum below) revealing an ee of >98%.

GC trace of (3*S*)-hydroxyoctan-2-one from the reaction with D469T

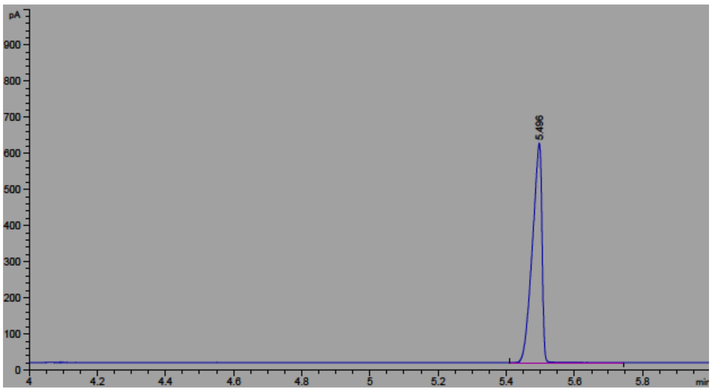


The product from the enzyme reaction was analysed further by derivatisation as the Mosher's ester.

**(2*S*,3*S*)-2-Oxooctan-3-yl-3,3,3-trifluoro-2-methoxy-2-phenylpropanoate**

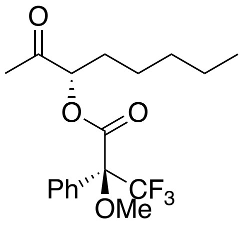


To a stirred solution of (3*S*)-hydroxyoctan-2-one (0.007 g, 0.05 mmol) in $CH_2Cl_2$ (5 mL) was added triethylamine (0.020 mL, 0.14 mmol) and (*R*)-MTPA-Cl (0.020 mL, 0.1 mmol) and the reaction stirred at room temperature for 18 h. The crude reaction mixture was then dry-loaded onto silica and purified by flash silica chromatography (EtOAc:hexane, 5:95) to give the titled compound as a colourless oil (1 mg, 5%)[7]. $R_f$ 0.5 (EtOAc/hexane, 1:9); $^1$H NMR (600 MHz; $CDCl_3$) $\delta$ 0.80-0.84 (3H, m, $CH_2CH_3$), 1.05-1.83 (8H, m, $(CH_2)_4$), 2.13 (0.05H, s, (2*R*,3*R*)-$COCH_3$), 2.22 (2.95H, s, (2*R*,3*S*)-$COCH_3$), 3.63 (3H, s, $OCH_3$), 5.18 (1H, dd, *J* = 4.6 and 4.1 Hz, C*H*OC=O), 7.43 (3H, m, Ar*H*), 7.64 (2H, m, Ar*H*); $^{13}$C NMR (151 MHz; $CDCl_3$) $\delta$ 13.9, 22.3, 24.8, 25.5, 30.0, 31.3, 55.7, 80.7, 84.7 (q, $^2J_{CF}$ = 28.1 Hz), 127.4, 128.5, 129.8, 132.1, 166.5, 203.7; *m/z* HRMS (CI+) found $[MH]^+$ 361.1636, $C_{18}H_{23}O_4F_3$ requires 361.1621.

**_Rac_-4-hydroxyoctan-3-one**

**Step 1**

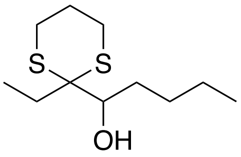

To solution of 2-ethyl-1,3-dithiane[1] (593 mg, 4.32 mmol) in dry THF (16 mL) at -78 °C, *n*-butyllithium (2.5 M in hexane; 2.0 mL, 5.0 mmol) was added dropwise. The reaction was stirred at -78 °C for 30 min, at 0 °C for a further 30 min and then cooled to -78 °C. Pentanal (638 µL, 6.00 mmol) was added dropwise and the reaction mixture stirred at room temperature for 18 h. Water (16 mL) was added, followed by dichloromethane (16 mL) and the product extracted with dichloromethane (3 × 50 mL). The volume of the combined organic fraction was reduced under vacuum to approximately 50 mL, then washed with aqueous potassium hydroxide solution (7%, 20 mL) and sat. sodium chloride solution (20 mL), dried ($MgSO_4$), and the solvent was removed *in vacuo*. The crude product was purified by flash silica column chromatography (EtOAc/pet. ether 40-60, 1:9) to give 1-(2-ethyl-1,3-dithian-2-yl)pentan-1-ol as a pale yellow oil (356 mg, 38%). R*f* 0.33 (EtOAc/pet. ether 40-60, 1:4); $\nu_{max}$ (film, cm$^{-1}$) 3477, 2961; $^1$H NMR (600 MHz; $CDCl_3$) δ 0.92 (3H, t, *J* = 7.1 Hz, $(CH_2)_2CH_3$), 1.07 (3H, t, *J* = 7.5 Hz, $CCH_2CH_3$), 1.19-1.49 (10H, m, 5 × $CH_2$), 1.65-1.71 (1H, m, CHOHC*H*H), 1.80-1.86 (1H, m, $SCH_2$C*H*H), 1.88-1.94 (1H, m, CHOHCH*H*), 2.02-2.08 (1H, m, $SCH_2$CH*H*), 2.60-2.66 (2H, m, 2 × SC*H*H), 2.94-3.00 (2H, m, 2 × SCH*H*), 3.66-3.76 (1H, m, C*H*OH), 3.95 (1H, br d, *J* = 9.3 Hz, OH); $^{13}$C NMR (151 MHz; $CDCl_3$) δ 9.4, 24.0, 24.6, 25.1, 25.9, 27.8, 29.2, 29.8, 31.1, 59.7, 71.0; *m/z* HRMS (ESI+) found $[MH]^+$ 235.1188, $C_{11}H_{23}OS_2$ requires 235.1190.

**Step 2**

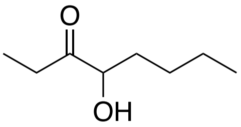

To a solution of 1-(2-ethyl-1,3-dithian-2-yl)pentan-1-ol (352 mg, 1.50 mmol) in water (40 mL), acetonitrile (8 mL) and THF (5 mL), iodomethane (2.99 mL, 48.2 mmol) and calcium carbonate (1.80 g, 18.0 mmol) were added and the reaction mixture was heated at reflux for 4 h. The reaction was cooled to room temperature and the product extracted with diethyl ether (3 × 50 mL), dried ($MgSO_4$), filtered, and the solvent removed *in vacuo*. The crude product was purified by flash silica column chromatography (diethyl ether/pentane, 1:9) to give 4-hydroxyoctan-3-one[8] as a pale yellow oil (149 mg, 69%). $R_f$ 0.24 (EtOAc/pet. ether 40-60, 1:9); $\nu_{max}$ (film, cm$^{-1}$) 3470, 2930, 1709, 1459; $^1$H NMR (600 MHz; $(CD_3)_2CO$) δ 0.91 (3H, t, *J* = 7.1 Hz, $CH_2CH_3$), 1.02 (3H, t, *J* = 7.3 Hz, $COCH_2CH_3$), 1.28-1.46 (4H, m, $(CH_2)_2CH_3$), 1.50-1.58 (1H, m, CHOHCH*H*), 1.72-1.79 (1H, m, CHOHC*H*H), 2.53-2.67 (2H, m, $COCH_2$), 4.04 (1H, d, *J* = 5.2 Hz, OH), 4.06-4.10 (1H, m, C*H*OH); $^{13}$C NMR (151 Mz; $(CD_3)_2CO$) δ 7.2, 13.7, 22.7, 27.5, 30.7, 33.8, 76.7, 213.4; *m/z* HRMS (ESI+) found $[M+NH_4]^+$ 162.1493, $C_8H_{20}NO_2$ requires 162.1494.

***(4S)*-4-Hydroxyoctan-3-one**

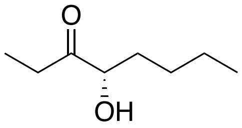

The co-factor solution (total volume 10 mL, pH 7) was prepared as described above in the general procedure for TK reactions. D469T TK cell-free lysate (2 mL) was added and the mixture was incubated for 20 min. To 2-ketobutyric acid (0.102 g, 1.00 mmol) in water (10 mL) and pentanal (0.106 mL, 1.00 mmol) the pH was adjusted to 7. This was then added to the enzyme suspension and the reaction was stirred in an autotitrator for 18 h while the reaction was followed by TLC analysis. The product was then extracted with $Et_2O$ (2 x 500 mL), and the organic layer was dried ($MgSO_4$), and then dry-loaded onto silica before being purified by flash silica chromatography ($Et_2O$:pentane, 5:95) to give (4*S*)-hydroxyoctan-3-one[9] as a pale green oil (0.006 g, 4%). The compound is volatile so some material was lost during the purification step. NMR characterization data was consistent with that of the racemate; $^1$H NMR (600 MHz; ($CDCl_3$) $\delta$ 0.89 (3H, t, $J$ = 7.1 Hz, $CH_2CH_3$), 1.00 (3H, t, $J$ = 7.3 Hz, $COCH_2CH_3$), 1.25-1.46 (4H, m, $(CH_2)_2CH_3$), 1.49-1.55 (1H, m, CHOHCH*H*), 1.70-1.78 (1H, m, CHOHC*H*H), 2.54-2.60 (2H, m, $COCH_2$), 4.04 (1H, d, $J$ = 5.2 Hz, OH), 4.06-4.10 (1H, m, C*H*OH); $^{13}$C NMR (151 Mz; ($CDCl_3$)) $\delta$ 7.2, 13.7, 22.7, 27.5, 30.7, 33.8, 76.7, 213.4. $[\alpha]^{20}_D$ = +83.3 (*c* 0.15, $CDCl_3$).

The product stereoselectivity was established by GC analysis; rt 5.5 min (*S*-isomer) and 5.7 min (*R*-isomer). The enzyme reaction gave a single peak at 5.5 min only (*S*-isomer) (example spectrum below) revealing an ee of >98%.

GC trace o**f** (4*S*)-hydroxyoctan-3-one from the reaction with D469T

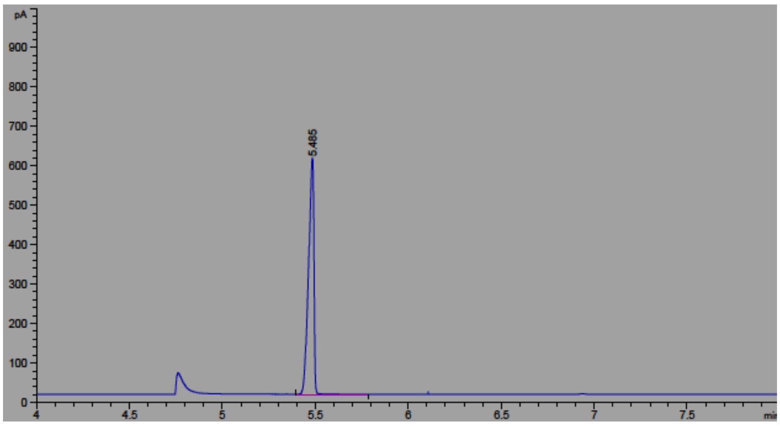


The product from the enzyme reaction was analysed further by derivatisation as the Mosher's ester.

**(2*S*,4*S*)-3-Oxooctan-4-yl-3,3,3-trifluoro-2-methoxy-2-phenylpropanoate**

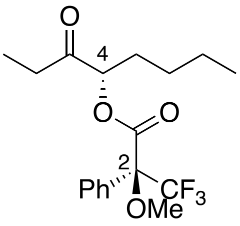


To a stirred solution of (4*S*)-hydroxyoctan-3-one (0.006 g, 0.04 mmol) in $CH_2Cl_2$ (5 mL) was added triethylamine (0.020 mL, 0.14 mmol) and (*R*)-MTPA-Cl (0.020 mL, 0.1 mmol) and the reaction stirred at room temperature for 18 h. The crude reaction mixture was then dry-loaded onto silica and purified by flash silica chromatography (EtOAc:hexane, 5:95) to give the titled compound as a colourless oil (1 mg, 6%). $R_f$ 0.52 (EtOAc/hexane, 1:9); $^1$H NMR (600 MHz; $CDCl_3$) $\delta$ 0.82 (3H, t, *J* = 7.0 Hz, $CH_2CH_3$), 1.10 (3H, t, *J* = 7.3 Hz, $COCH_2CH_3$), 1.18-1.84 (6H, m, $(CH_2)_3$), 2.51 (2H, m, $COCH_2CH_3$), 3.64 (3H, s, $OCH_3$), 5.15-5.20 (1H, m, CHOC=O), 7.42-65 (5H, m, Ar*H*); $^{13}$C NMR (151 MHz; $CDCl_3$) $\delta$ 7.4, 14.3, 25.4, 27.1, 29.8, 33.9, 55.8, 80.1, 84.7 (q, $^2J_{CF}$ = 28.1 Hz), 127.6, 128.9, 129.8, 132.1, 166.5, 206.5; $^{19}$F NMR (282 MHz; $CDCl_3$) $\delta$ –72.1; *m/z* HRMS (CI+) found $[MH]^+$ 361.1627, $C_{18}H_{24}O_4F_3$ requires 361.1633.

***Rac*-4-hydroxynonan-3-one**

**Step 1**

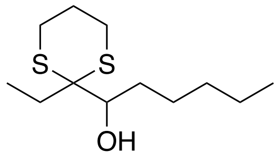

To solution of 2-ethyl-1,3-dithiane[1] (593 mg, 4.32 mmol) in dry THF (16 mL) at -78 °C, *n*-butyllithium (2.5 M in hexane; 2.0 mL, 5.0 mmol) was added dropwise. The reaction was stirred at -78 °C for 30 min, at 0 °C for a further 30 min and then cooled to -78 °C. Hexanal (737 µL, 6.00 mmol) was added dropwise and the reaction mixture stirred at room temperature for 18 h. Water (16 mL) was added, followed by dichloromethane (16 mL) and the product extracted with dichloromethane (3 × 50 mL). The volume of the combined organic fraction was reduced under vacuum to approximately 50 mL, then washed with aqueous potassium hydroxide solution (7%, 20 mL) and sat. sodium chloride solution (20 mL), dried ($MgSO_4$), and the solvent was removed *in vacuo*. The crude product was purified by flash silica column chromatography (EtOAc/pet. ether 40-60, 1:9) to give 1-(2-ethyl-1,3-dithian-2-yl)hexan-1-ol as a pale yellow oil (397 mg, 40%). R*f* 0.34 (EtOAc/pet. ether 40-60, 1:4); $\nu_{max}$ (film, $cm^{-1}$) 3478, 2953; $^1H$ NMR (600 MHz; $CDCl_3$) $\delta$ 0.92 (3H, t, *J* = 7.1 Hz, $(CH_2)_4CH_3$), 1.07 (3H, t, *J* = 7.5 Hz, $CCH_2CH_3$), 1.21-1.48 (8H, m, 4 x $CH_2$), 1.65-1.69 (1H, m, CHOHC*H*H), 1.80-1.86 (1H, m, $SCH_2$C*H*H), 1.88-1.92 (1H, m, CHOHCH*H*), 2.02-2.06 (1H, m, $SCH_2$CH*H*), 2.60-2.66 (2H, m, 2 × SC*H*H), 2.94-3.00 (2H, m, 2 × SCH*H*), 3.70-3.76 (1H, m, C*H*OH), 3.97 (1H, d, *J* = 9.6 Hz, OH); $^{13}C$ NMR (151 MHz; $CDCl_3$) $\delta$ 9.4, 23.9, 24.5, 25.1, 26.0, 26.7, 27.8, 29.2, 29.8, 31.5, 59.7, 71.0; *m/z* HRMS (ESI+) found $[MH]^+$ 249.1344, $C_{12}H_{25}OS_2$ requires 249.1347.

**Step 2**

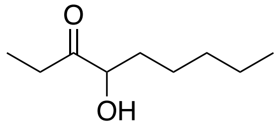

To a solution of 1-(2-ethyl-1,3-dithian-2-yl)hexan-1-ol (373 mg, 1.50 mmol) in water (40 mL), acetonitrile (8 mL) and THF (5 mL), iodomethane (2.99 mL, 48.2 mmol) and calcium carbonate (1.80 g, 18.0 mmol) were added and the reaction mixture was heated at reflux for 4 h. The reaction was cooled to room temperature and the product extracted with diethyl ether (3 × 50 mL), dried ($MgSO_4$), filtered, and the solvent removed *in vacuo*. The crude product was purified by flash silica column chromatography (diethyl ether/pentane, 1:9) to give 4-hydroxynonan-3-one[10] as a pale yellow oil (190 mg, 80%). $R_f$ 0.25 (EtOAc/pet. ether 40-60, 1:9); $\nu_{max}$ (film, $cm^{-1}$) 3447, 2933, 1709, 1459; $^1H$ NMR (600 MHz; $(CD_3)_2CO$) $\delta$ 0.90 (3H, t, *J* = 7.1 Hz, $CH_2CH_3$), 1.02 (3H, t, *J* = 7.3 Hz, $COCH_2CH_3$), 1.28-1.47 (6H, m, $(CH_2)_3CH_3$), 1.48-1.58 (1H, m, CHOHCH*H*), 1.71-1.79 (1H, m, CHOHC*H*H), 2.52-2.66 (2H, m, $COCH_2$), 4.05 (1H, d, *J* = 5.2 Hz, OH), 4.08-4.13 (1H, m, C*H*OH); $^{13}C$ NMR (151 Mz; $(CD_3)_2CO$) $\delta$ 7.2, 13.8, 22.7, 25.0, 30.7, 31.9, 34.1, 76.7, 213.4; *m/z* HRMS (ESI+) found $[M+NH_4]^+$ 176.1645, $C_9H_{22}NO_2$ requires 176.1645.

**(4*S*)-4-Hydroxynonan-3-one**

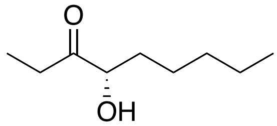

The co-factor solution (total volume 10 mL, pH 7) was prepared as described above in the general procedure for TK reactions. D469E cell-free lysate (2 mL) was added and the mixture was incubated for 20 min. To 2-ketobutyric acid (0.102 g, 1.00 mmol) in water (10 mL) and hexanal (0.123 mL, 1.00 mmol) the pH was adjusted to 7. This was then added to the enzyme suspension and the reaction was stirred in an autotitrator for 24 h while the reaction was followed by TLC analysis. The product was then extracted with $Et_2O$ (2 x 500 mL), and the organic layer was dried ($MgSO_4$), and then dry-loaded onto silica before being purified by flash silica chromatography ($Et_2O$:pentane, 5:95) to give (4*S*)-hydroxyoctan-3-one as a pale green oil (13 mg, 8%). The yield was comparable using D469T and the compound is volatile so some material was lost during the purification step. NMR characterization data was consistent with that of the racemate; $^1$H NMR (600 MHz; ($CDCl_3$)) δ 0.90 (3H, t, *J* = 7.1 Hz, $CH_2CH_3$), 1.02 (3H, t, *J* = 7.3 Hz, $COCH_2CH_3$), 1.28-1.47 (6H, m, $(CH_2)_3CH_3$), 1.48-1.58 (1H, m, CHOHCH*H*), 1.71-1.79 (1H, m, CHOHC*H*H), 2.52-2.66 (2H, m, $COCH_2$), 4.05 (1H, d, *J* = 5.2 Hz, OH), 4.08-4.13 (1H, m, C*H*OH); $^{13}$C NMR (151 MHz; ($CDCl_3$)) δ 7.6, 14.0, 22.5, 24.5, 31.1, 31.6, 33.8, 76.2, 212.9 $[\alpha]^{20}_D$ = +95.3 (*c* 0.15, $CDCl_3$).

**NMR spectra**

**(*rac*)-4-Hydroxyoctan-3-one**

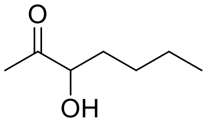


**[1]H NMR**

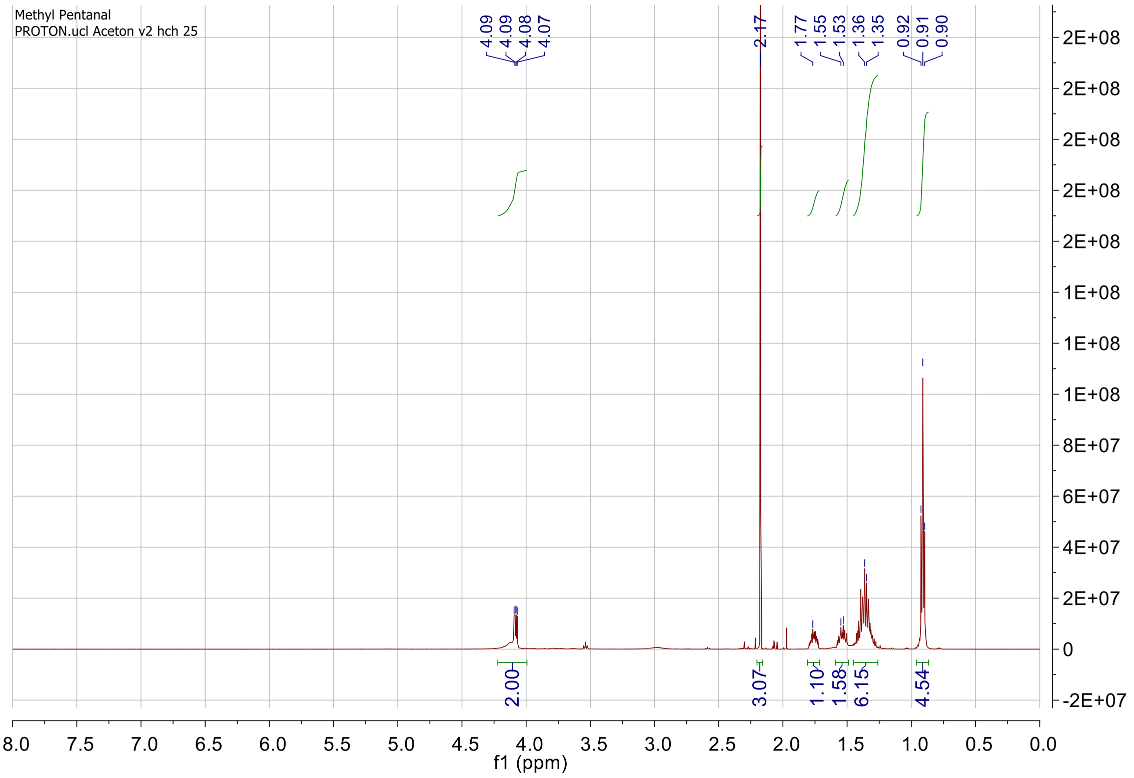


**[13]C NMR**

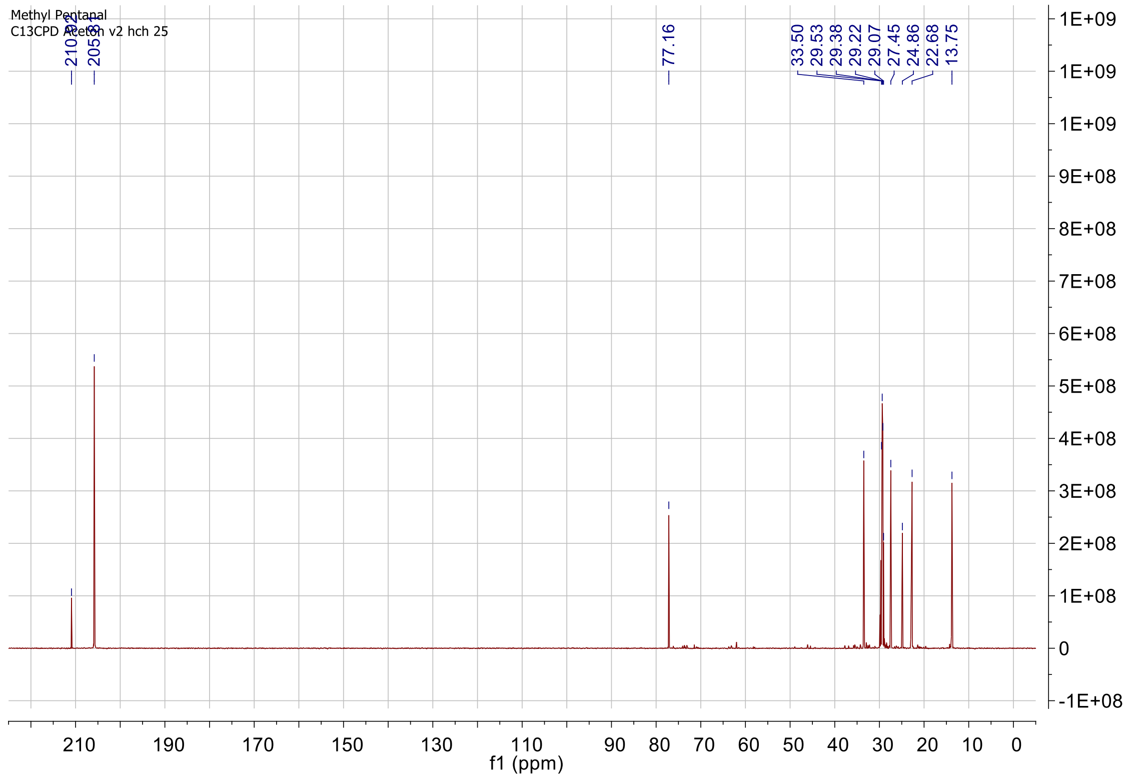

### (3*S*)-Hydroxyheptan-2-one

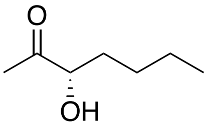

#### $^{1}$H NMR

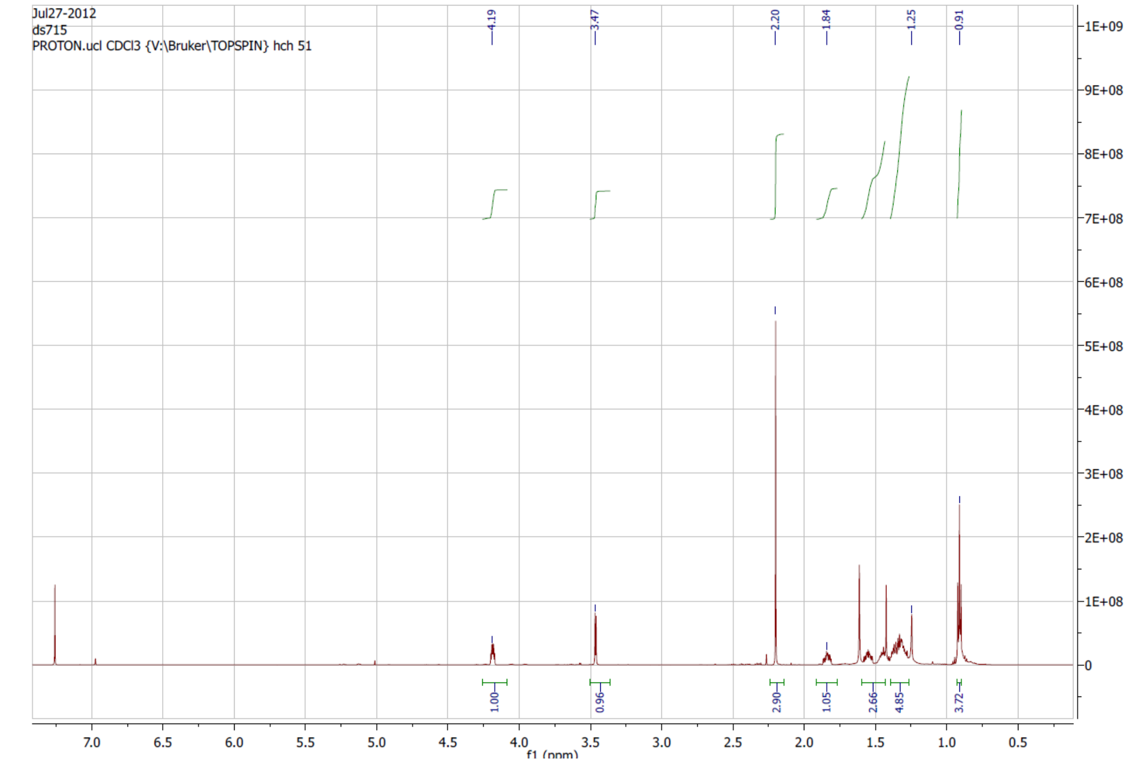


#### $^{13}$C NMR

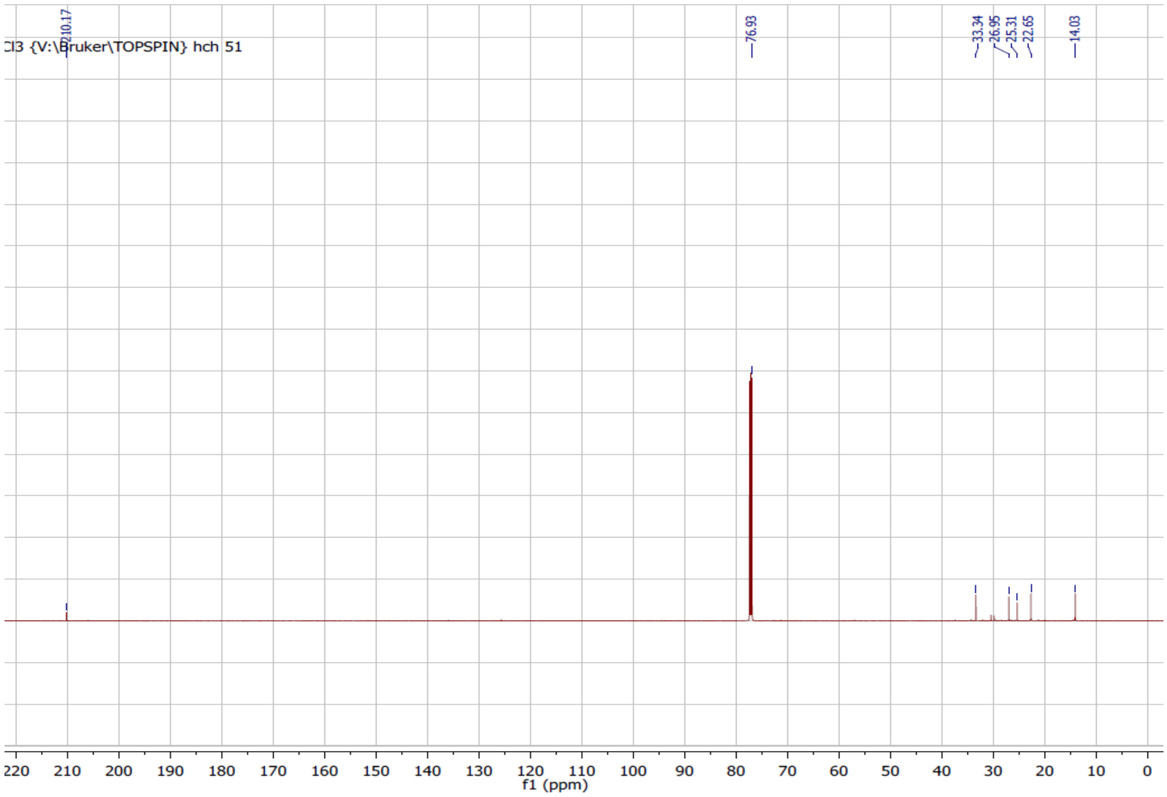

### (2*S*,3*S*)-2-Oxoheptan-3-yl-3,3,3-trifluoro-2-methoxy-2-phenylpropanoate

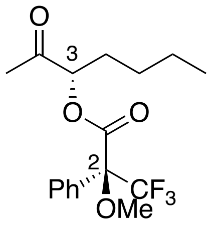

### [1]H NMR

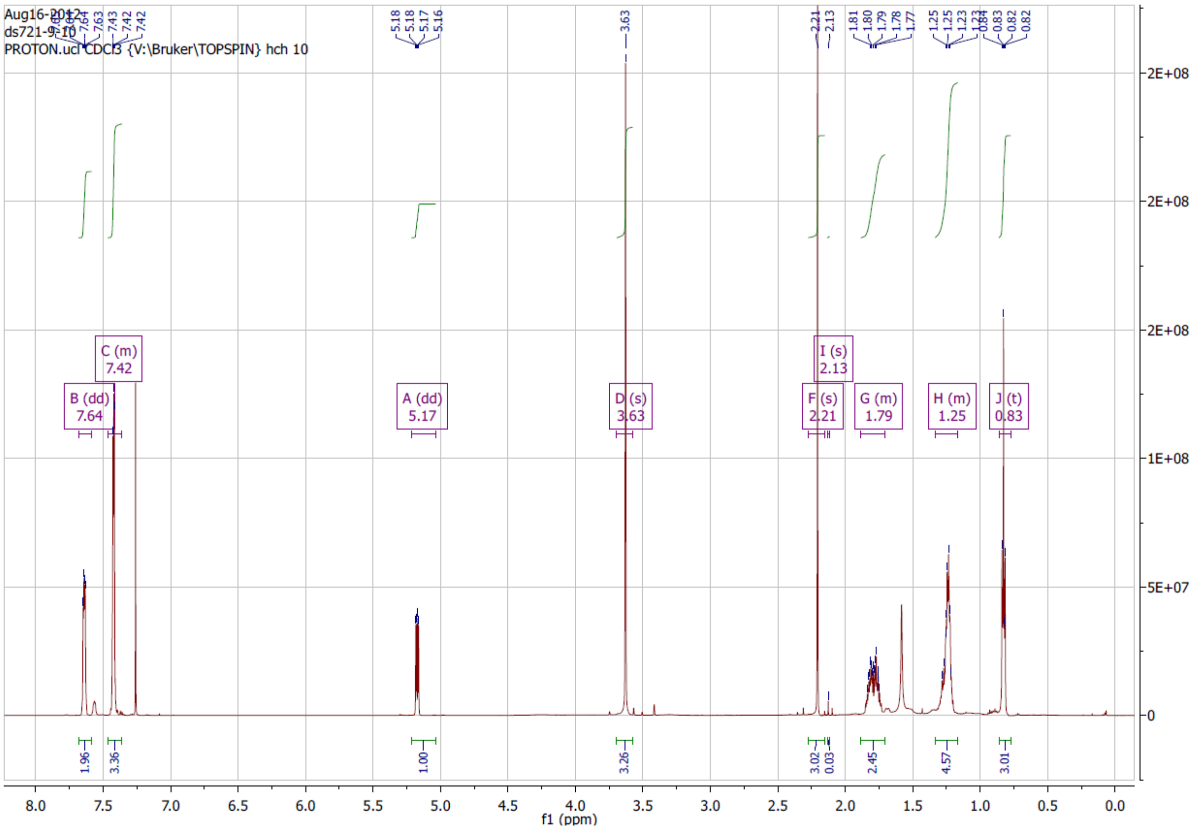

### [1]H NMR 2.0 to 2.3 ppm

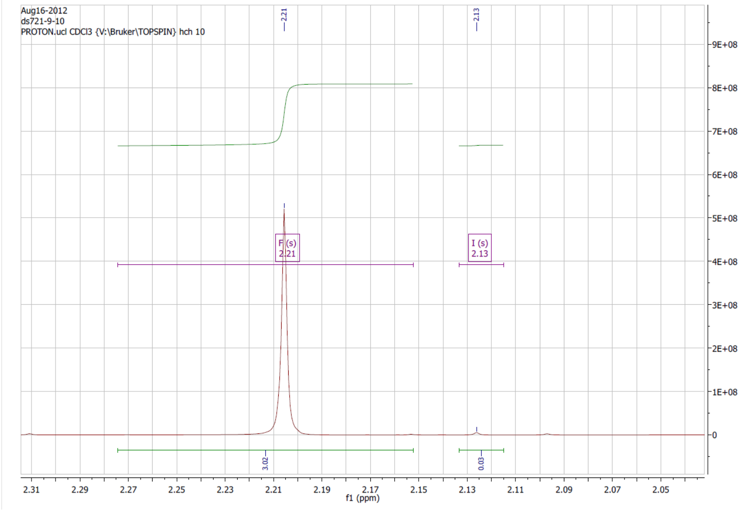

### $^{13}C$ NMR

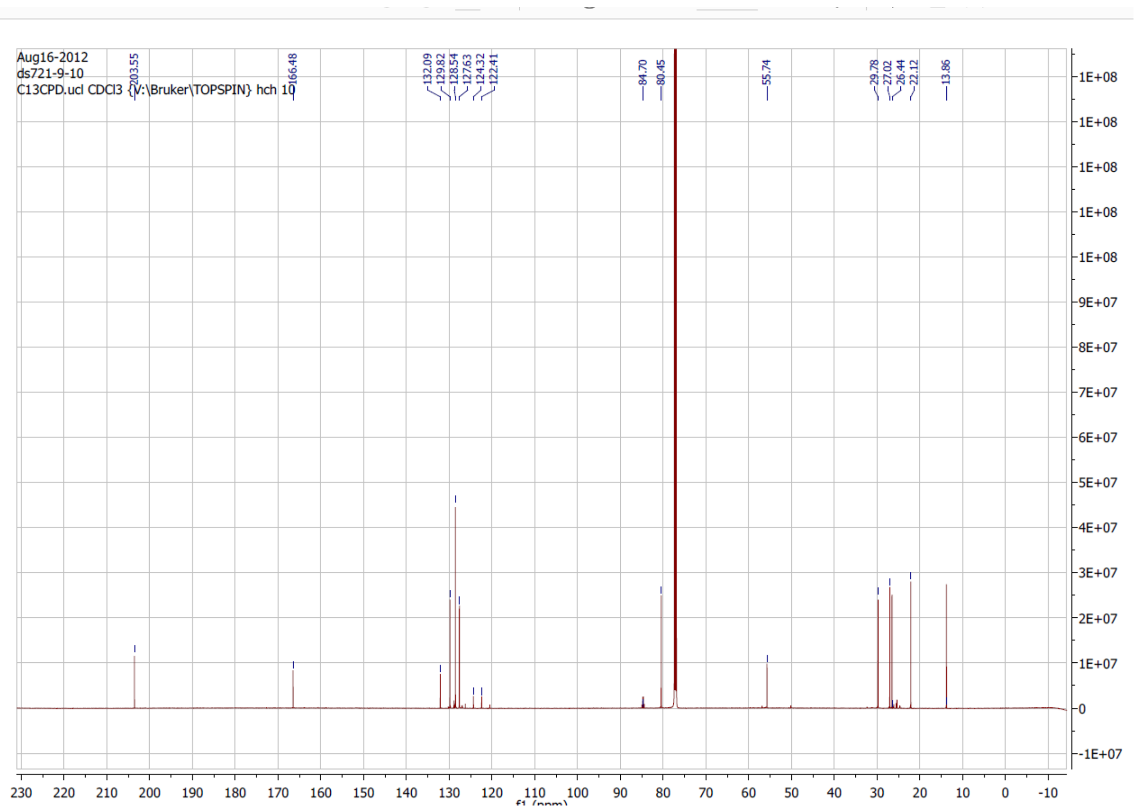

Aug16-2012
ds721-9-10
C13CPD.ucl CDCl3 {V:\Bruker\TOPSPIN} hch 10
203.55
166.48
132.09
129.82
128.54
127.63
124.32
122.41
84.70
80.45
55.74
29.78
27.02
26.44
22.12
13.86
f1 (ppm)

### (*rac*)-3-Hydroxyoctan-2-one

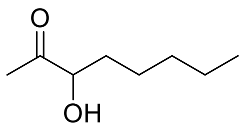


#### $^{1}$H NMR

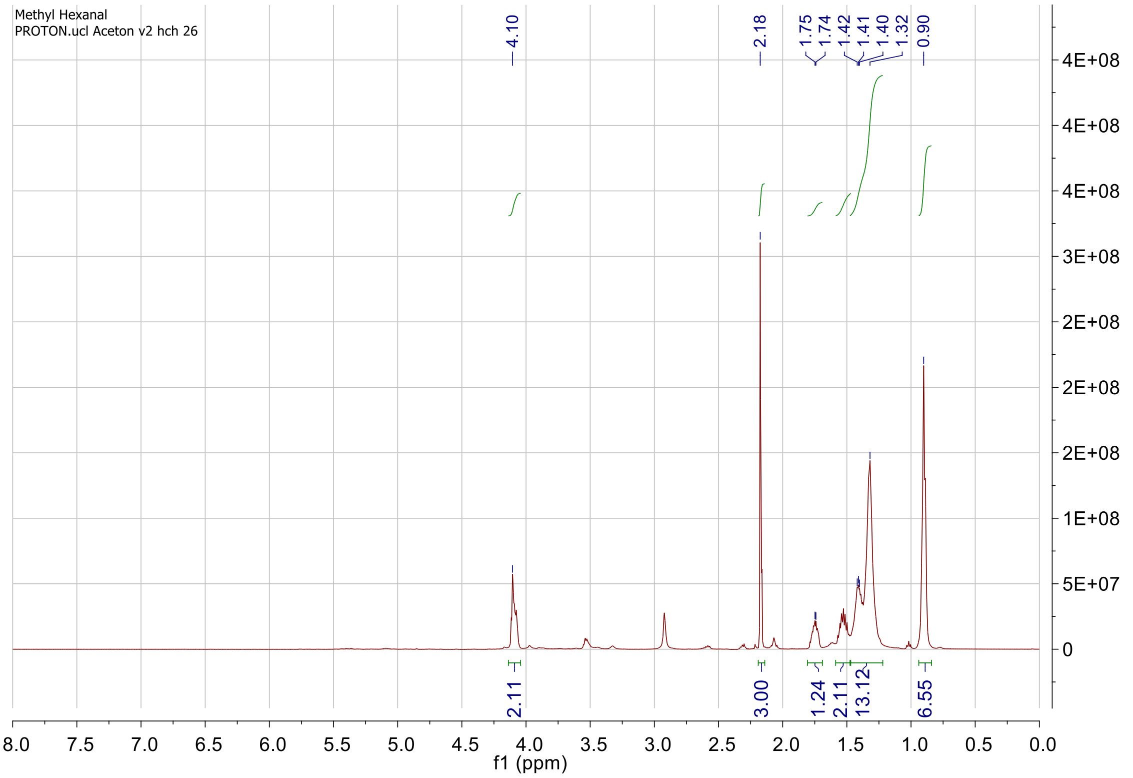


#### $^{13}$C NMR

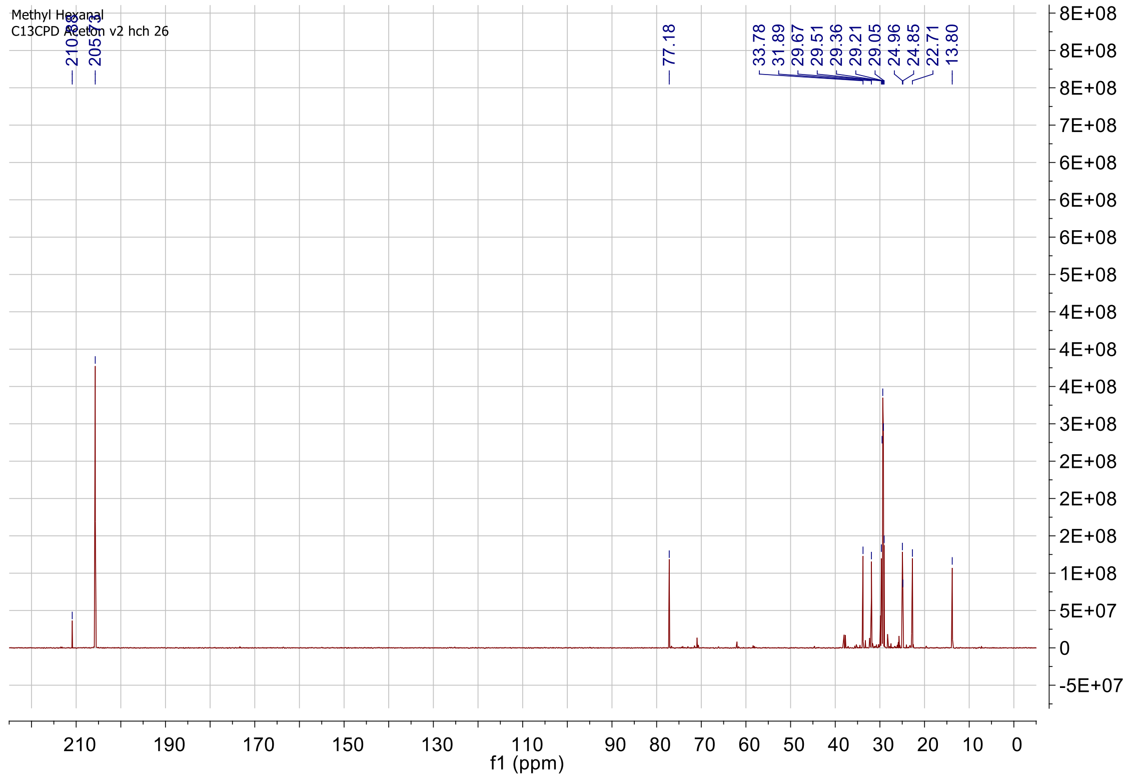

**(3*S*)-Hydroxyoctan-2-one**

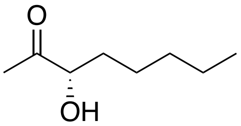

**[1]H NMR**

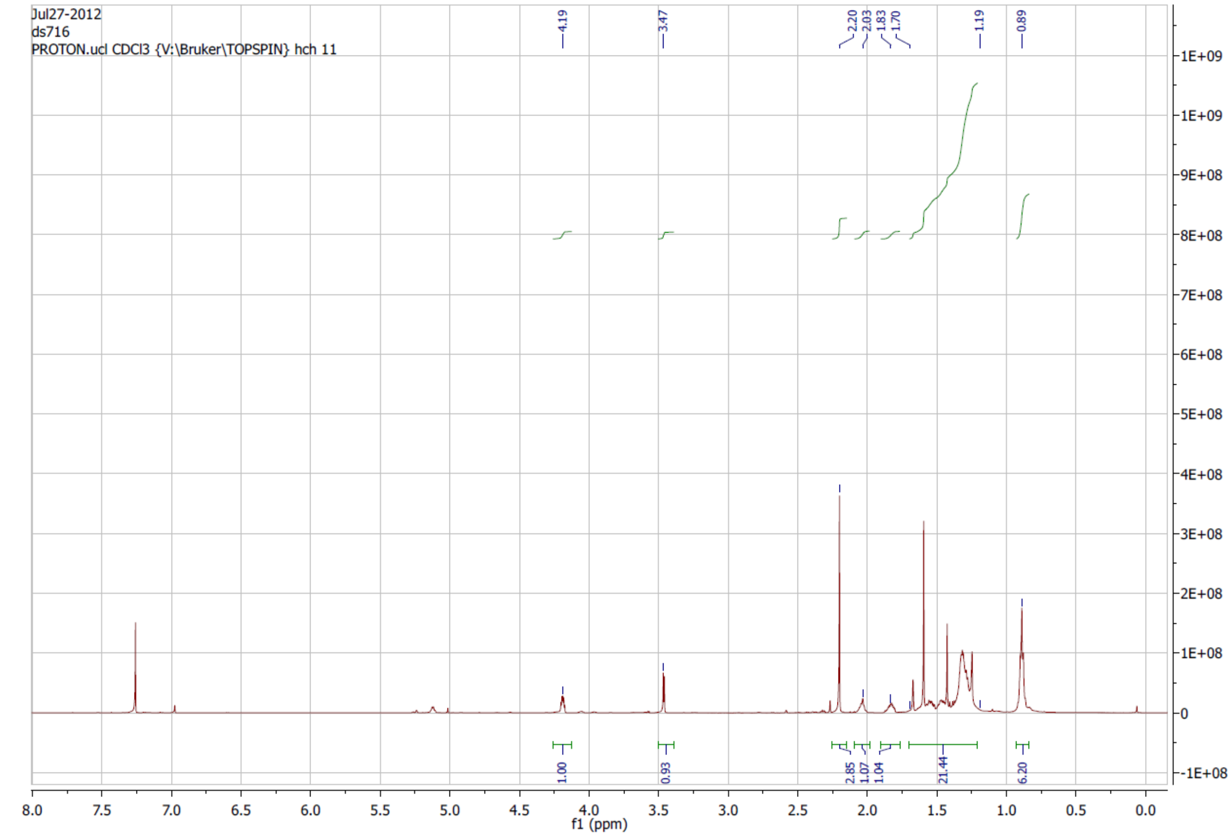


**[13]C NMR**

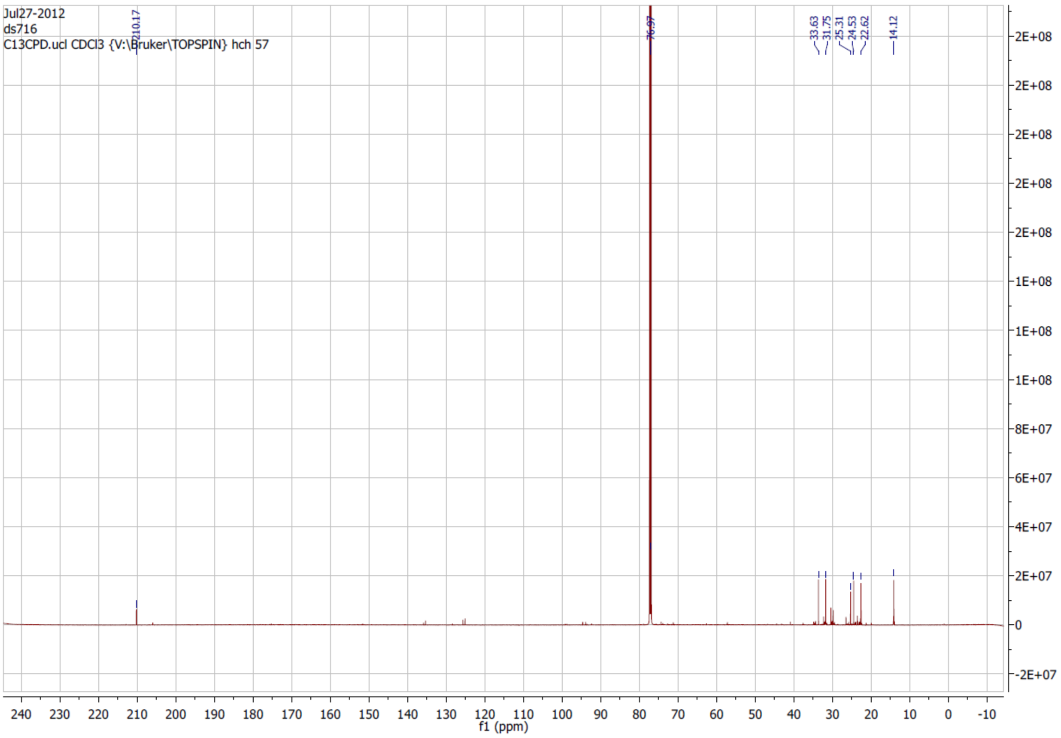

### (2*S*,3*S*)-2-Oxooctan-3-yl-3,3,3-trifluoro-2-methoxy-2-phenylpropanoate

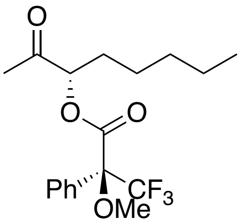

### $^{1}$H NMR

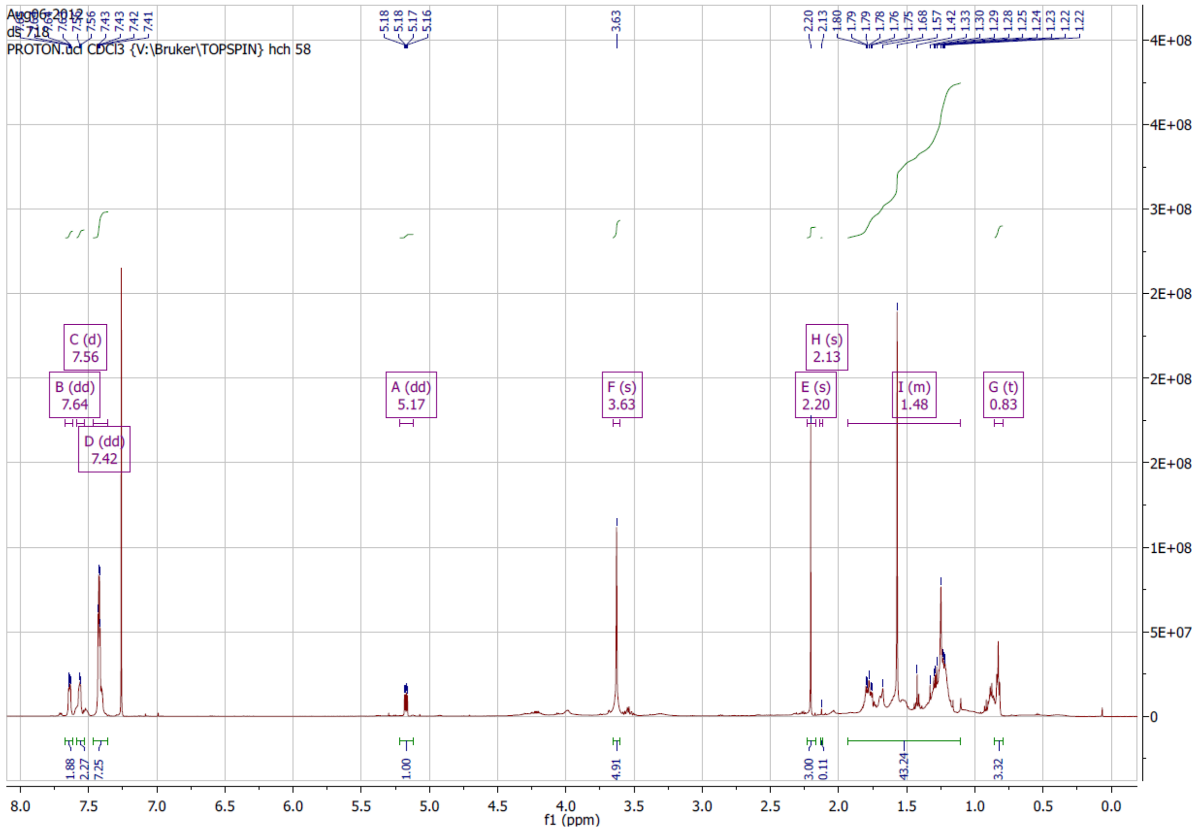


### $^{1}$H NMR 2.0 to 2.3 ppm

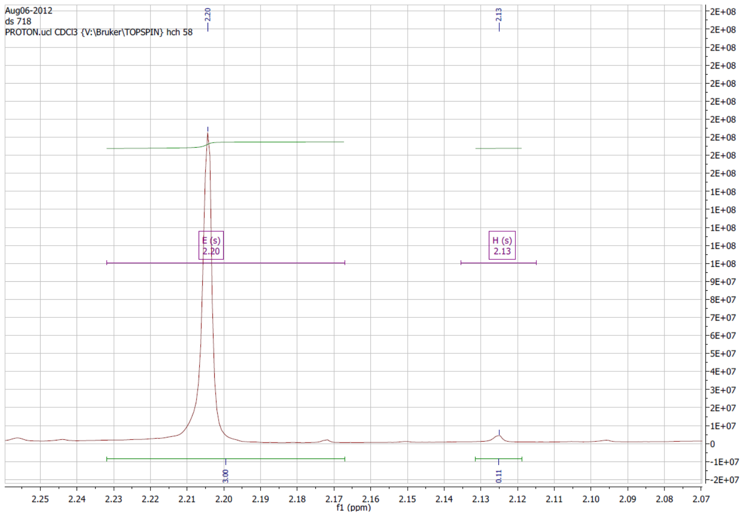

### $^{13}C$ NMR

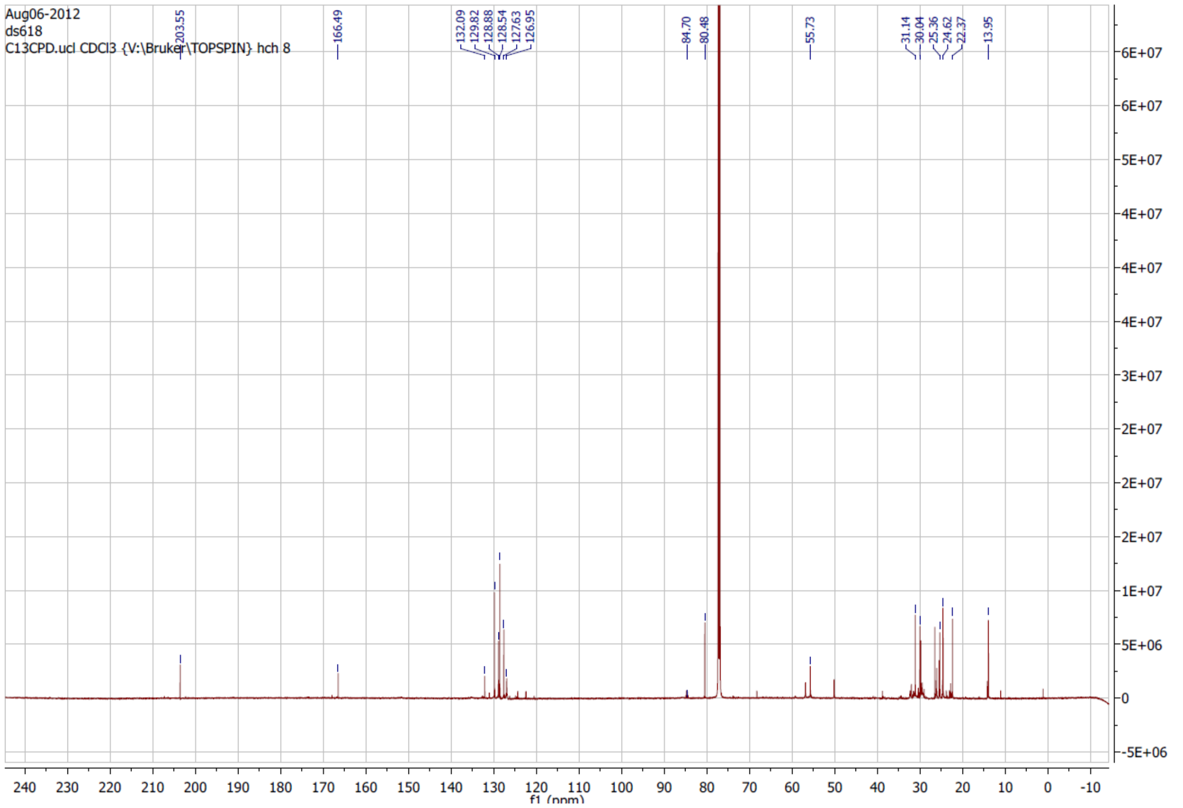

Aug06-2012
ds618
C13CPD.ucl CDCl3 {V:\Bruker\TOPSPIN} hch 8
203.55
166.49
132.09
129.82
128.88
128.54
127.63
126.95
84.70
80.48
55.73
31.14
30.04
25.36
24.62
22.37
13.95
f1 (ppm)

### (*rac*)-4-Hydroxyoctan-3-one

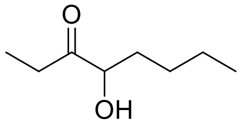

#### $^{1}$H NMR

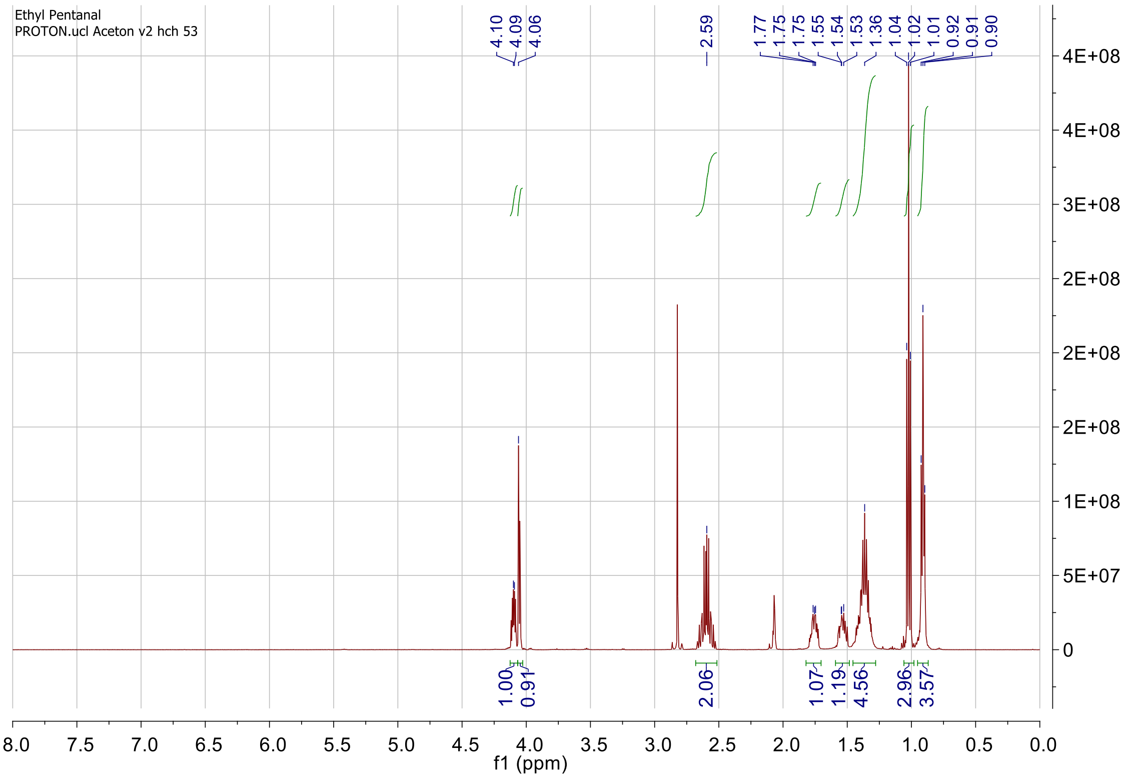

#### $^{13}$C NMR

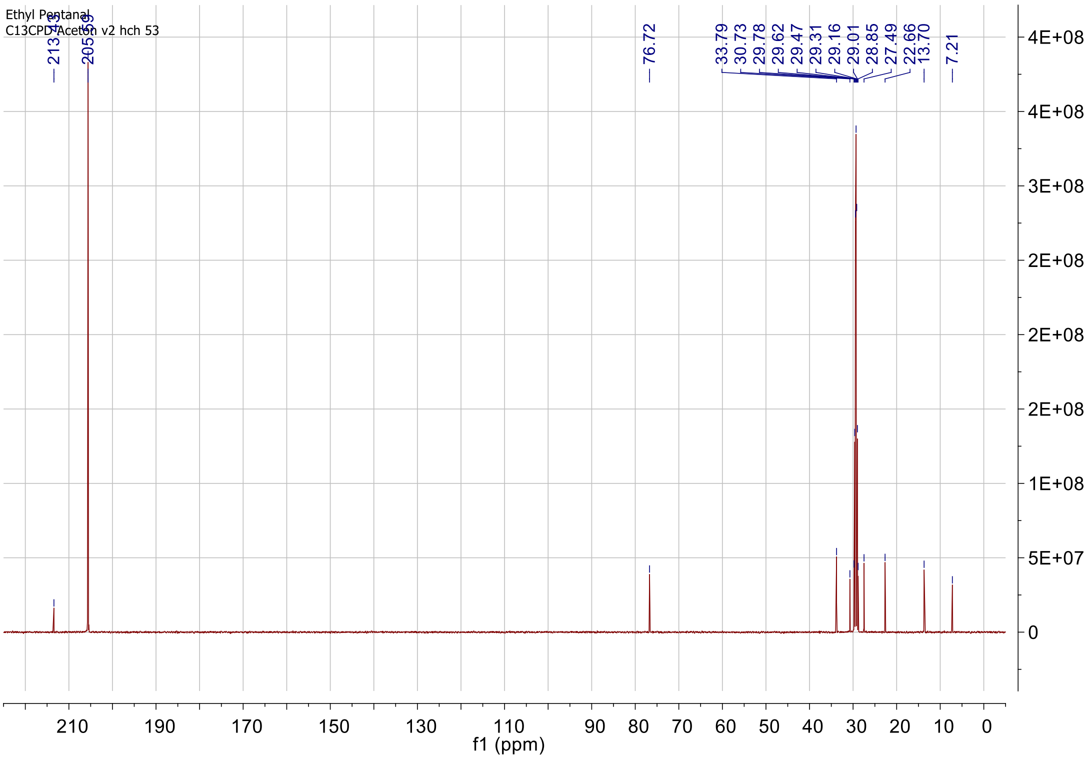

### *(4S)*-4-Hydroxyoctan-3-one

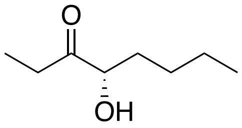

### [1]H NMR

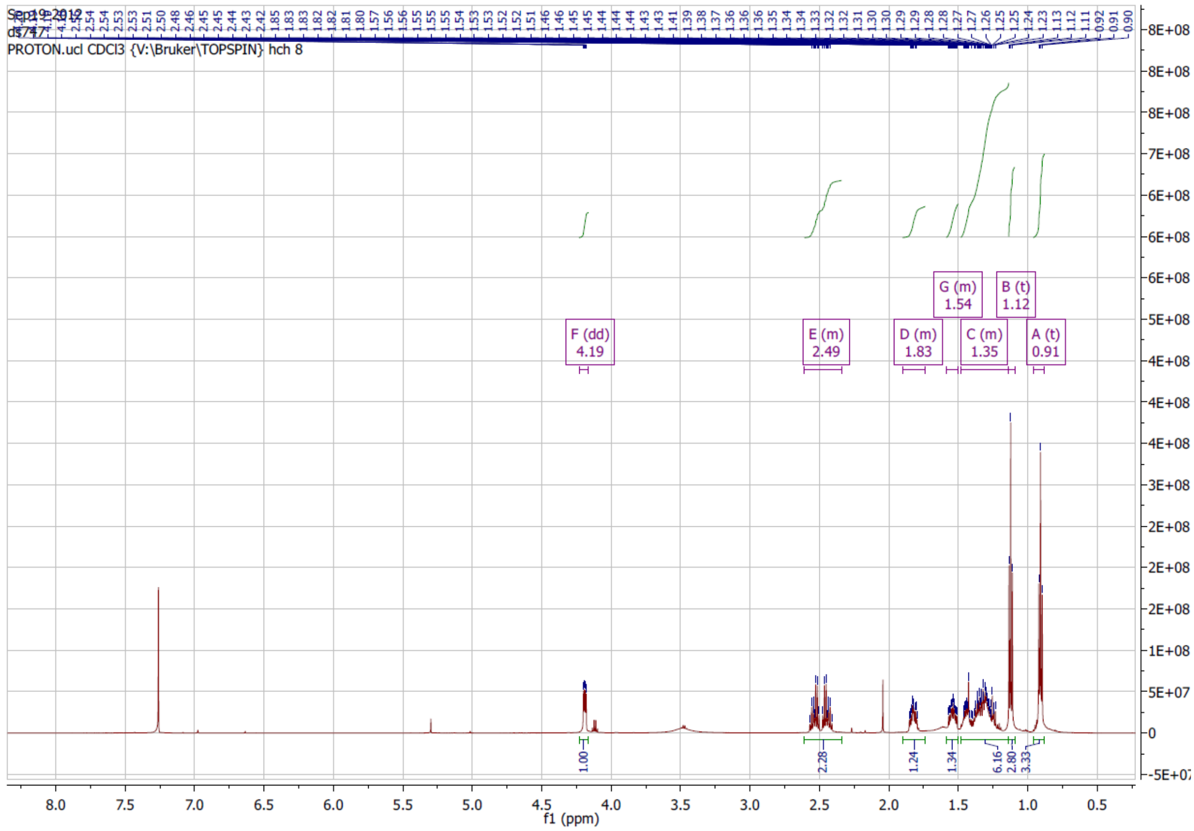

### (2*S*,4*S*)-3-Oxooctan-4-yl-3,3,3-trifluoro-2-methoxy-2-phenylpropanoate

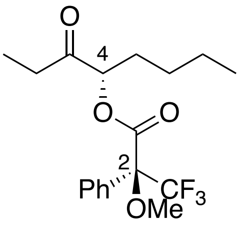


### [1]H NMR

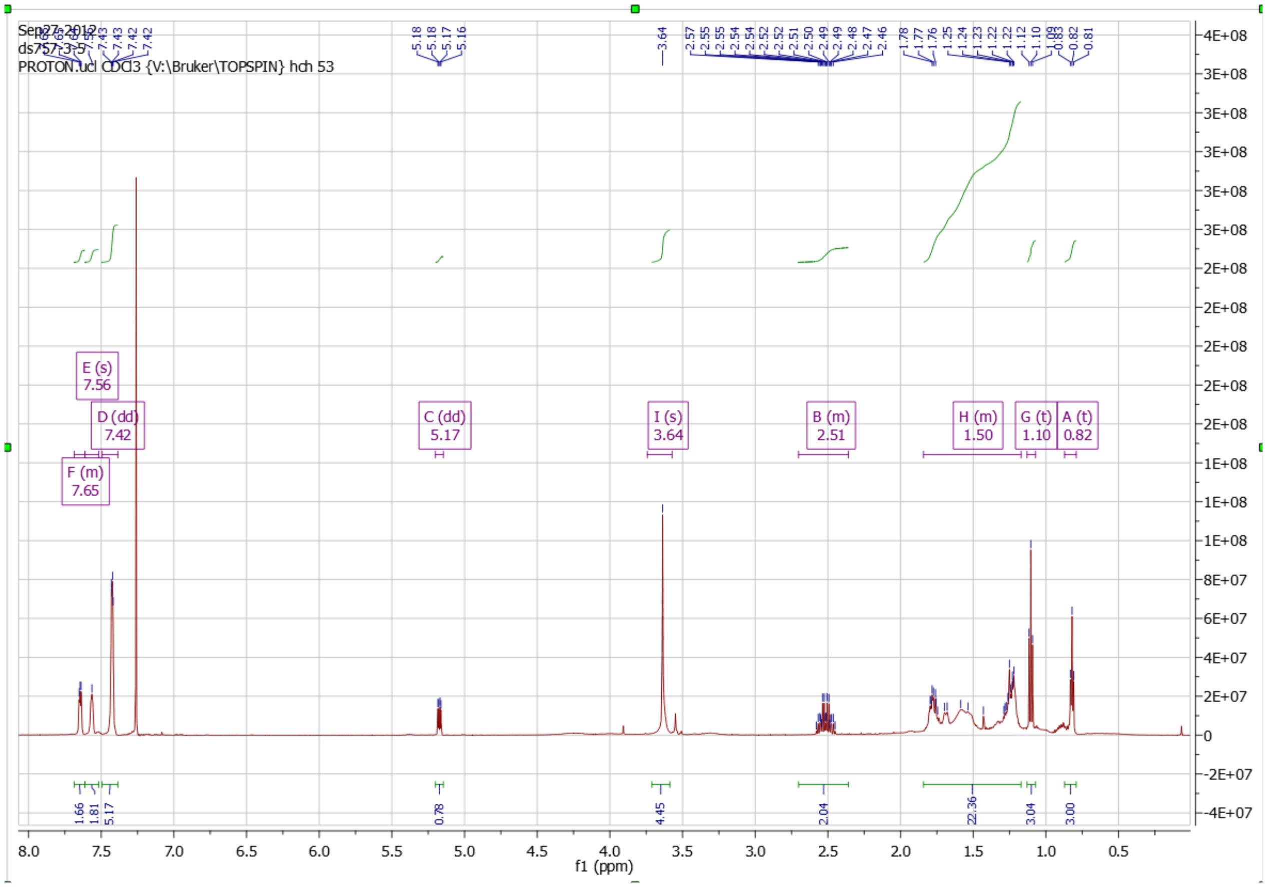

### [1]H NMR 0.7 to 1.2 ppm

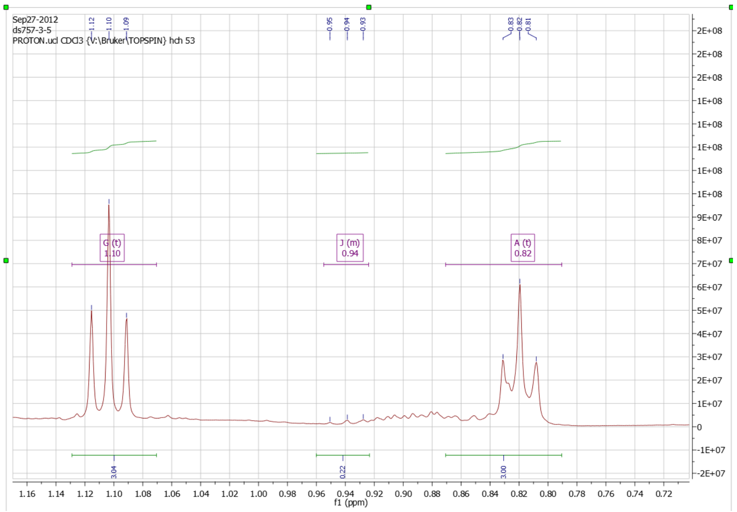

### $^{13}$C NMR

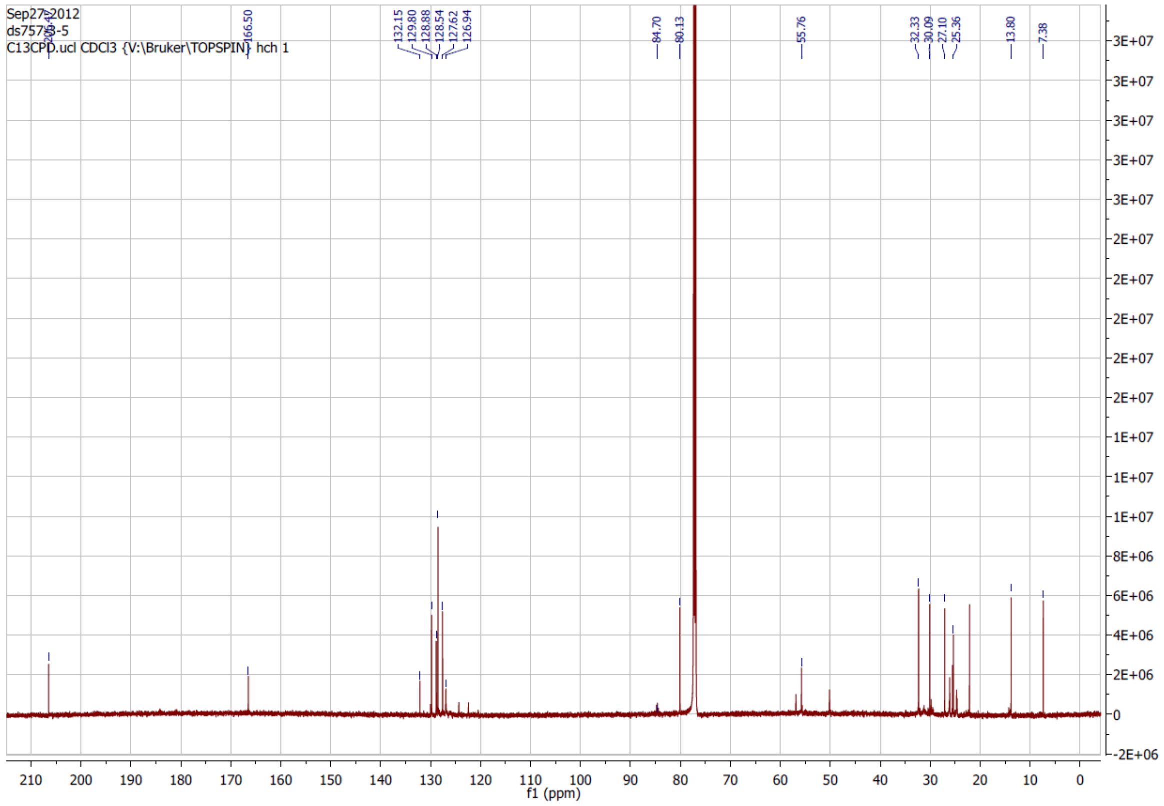

Sep27-2012
ds757-3-5
C13CPD.ucl CDCl3 {V:\Bruker\TOPSPIN} hch 1
166.50
132.15
129.80
128.88
128.54
127.62
126.94
84.70
80.13
55.76
32.33
30.09
27.10
25.36
13.80
7.38
f1 (ppm)

### (*rac*)-4-Hydroxynonan-3-one

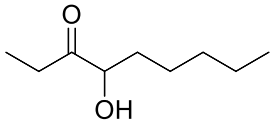


### [1]H NMR

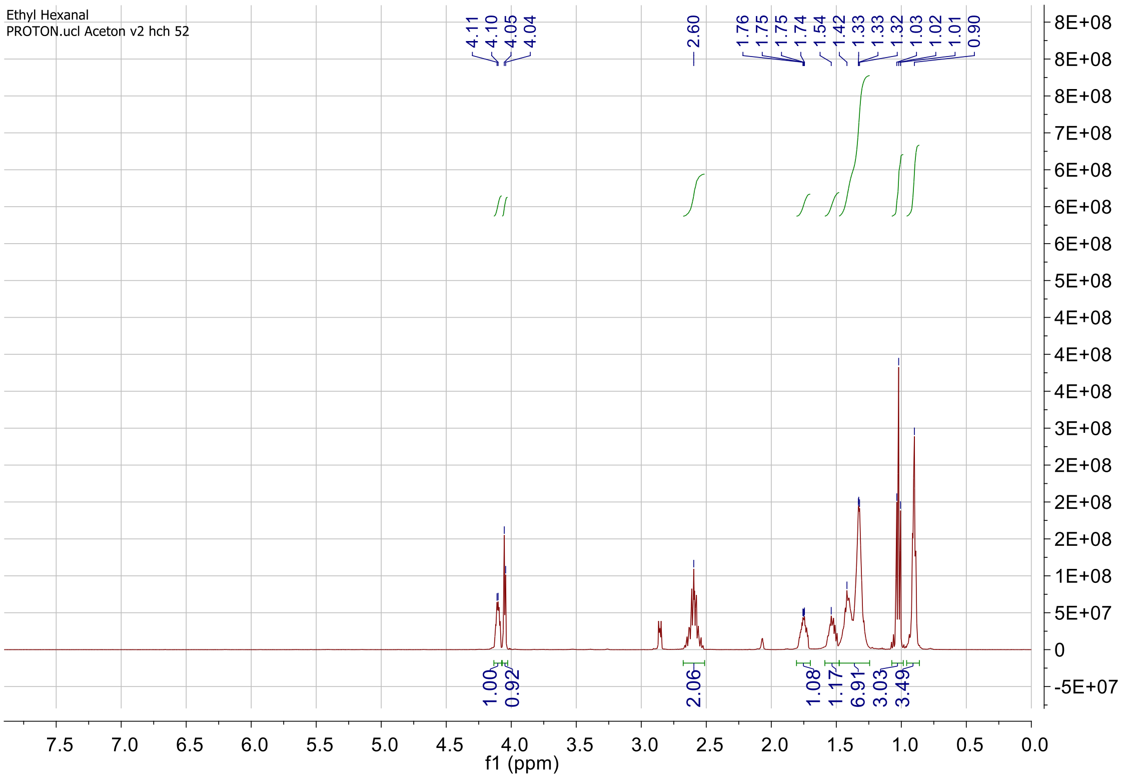


### [13]C NMR

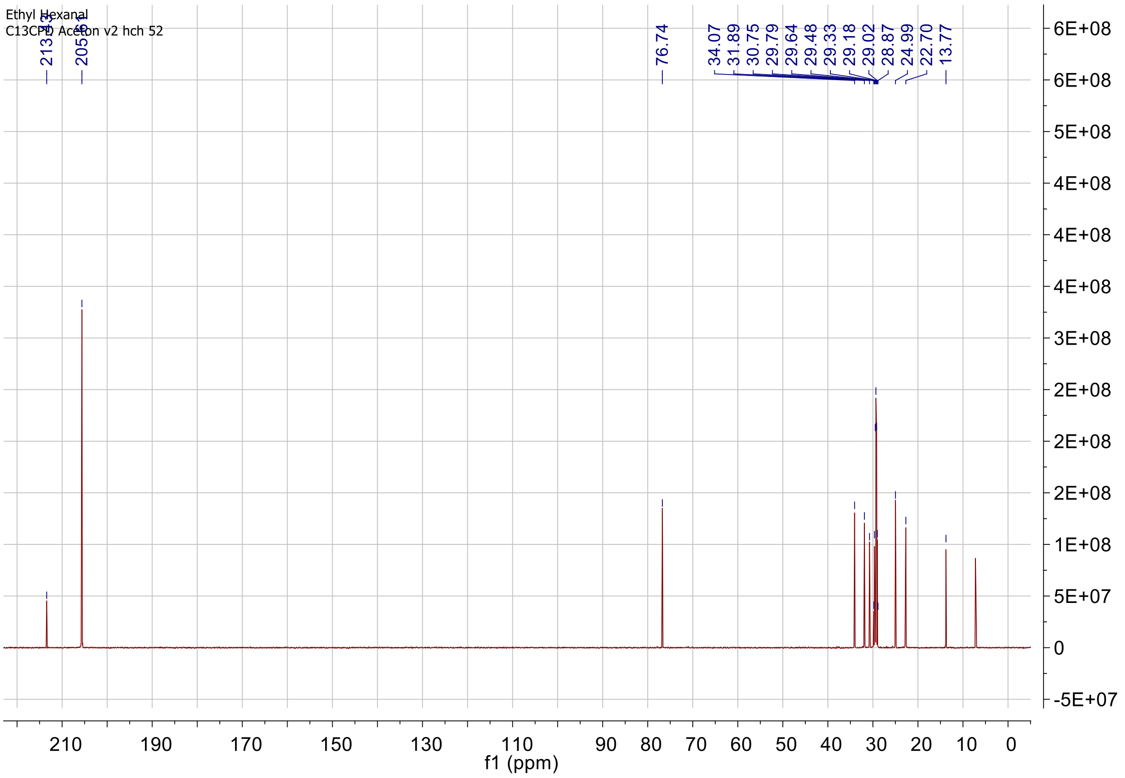

**(4*S*)-4-Hydroxynonan-3-one**

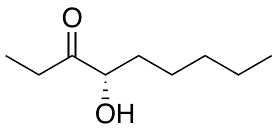


### $^1$H NMR

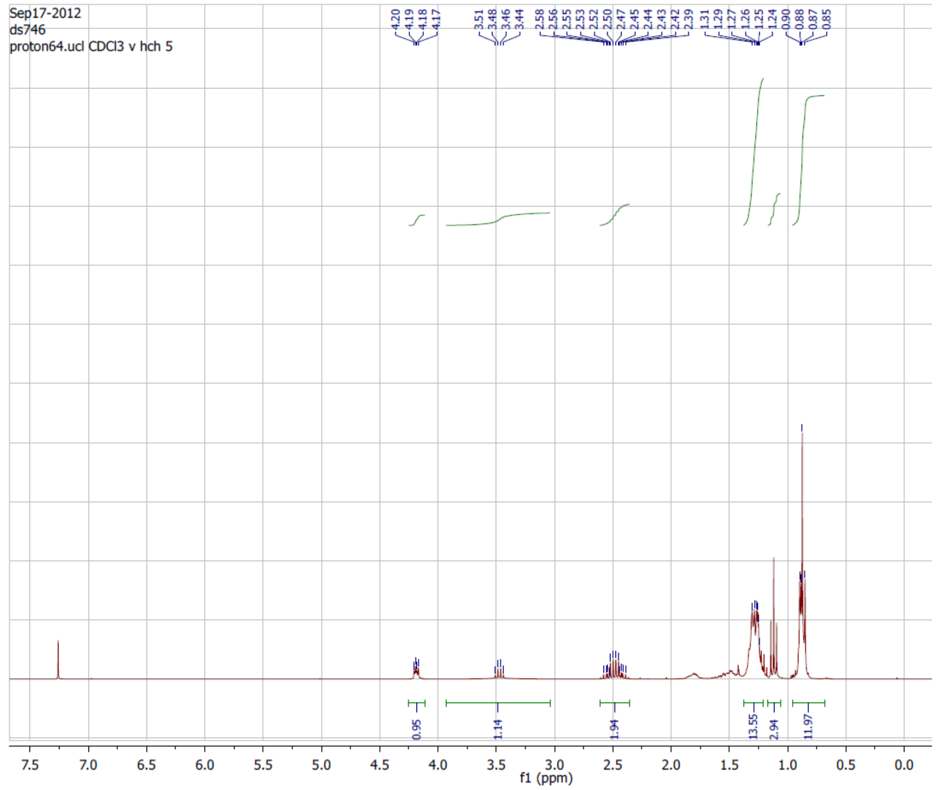


### $^{13}$C NMR

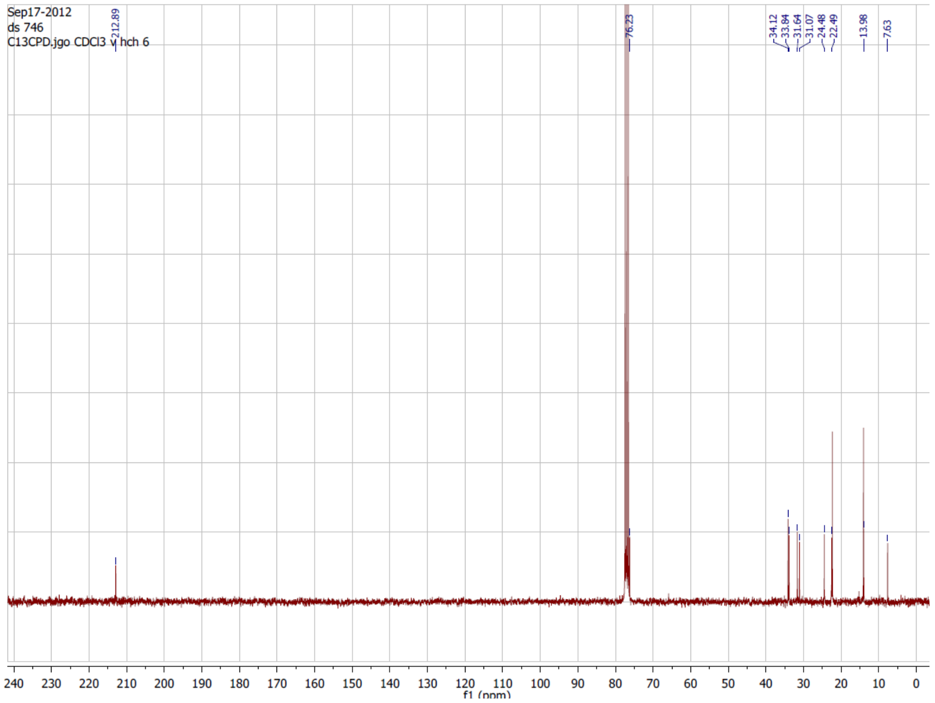

### 3-(1-hydroxy-2-oxopropyl) benzoic acid

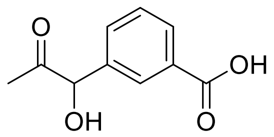


### $^1$H NMR

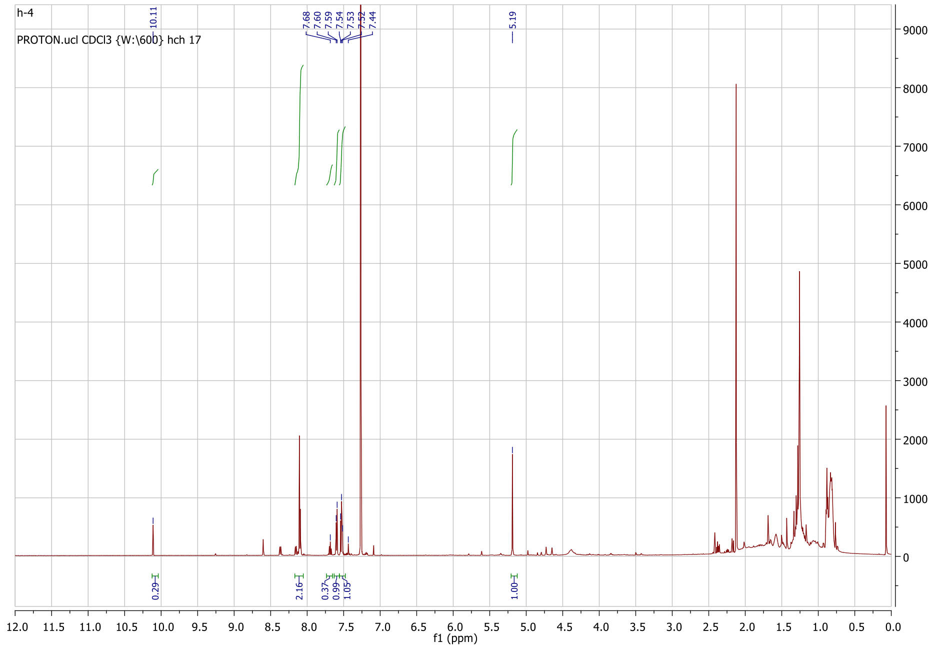


### $^{13}$C NMR

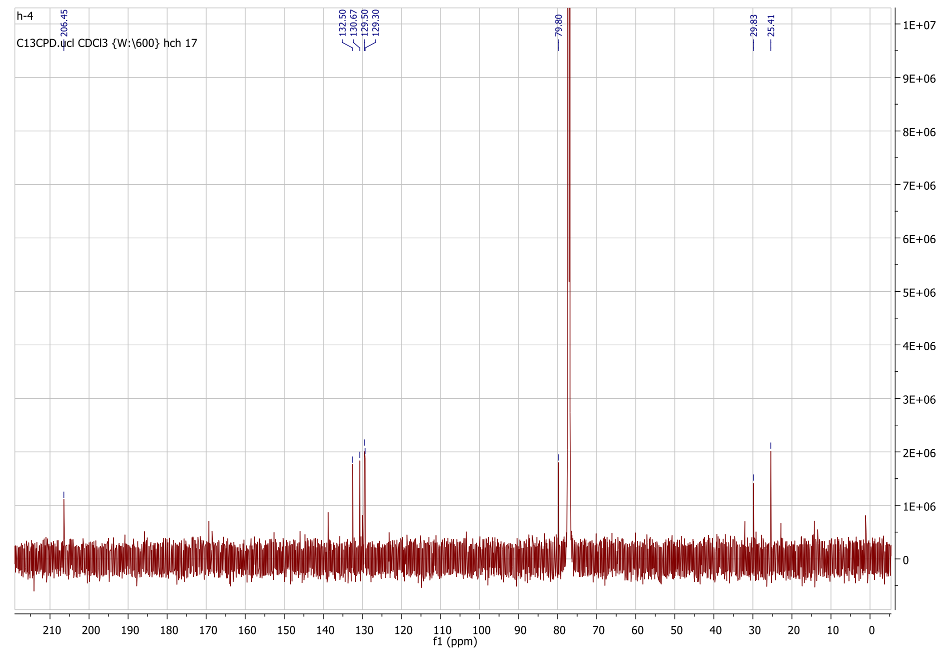
